# Proximal Pulmonary Artery Stiffening as a Biomarker of Cardiopulmonary Aging

**DOI:** 10.1101/2025.06.10.658632

**Authors:** Ruben De Man, Zhongyu Cai, Pramath Doddaballapur, Nicole Guerrera, Liqin Lin, Aurelien Justet, Nebal S. Abu Hussein, Cristina Cavinato, Micha Sam B. Raredon, Paul Heerdt, Inderjit Singh, Xiting Yan, Min-Jong Kang, Danielle R. Bruns, Patty J. Lee, George Tellides, Jay D. Humphrey, Naftali Kaminski, Abhay B. Ramachandra, Edward P. Manning

## Abstract

The geroscience hypothesis suggests that understanding underlying ageing mechanisms will enable us to delay aging and lessen age-related disability and diseases. While hallmarks of ageing list multiple contributing factors, role of mechanics has only been recently recognized and increasingly appreciated. Here, we use mouse models of ageing to investigate changes in mechanics of the proximal pulmonary artery, lung and right ventricle function in ageing. We found an age-related decline in the capacity to store energy and increased circumferential stiffness of the proximal pulmonary artery with age that associated with a reorientation of collagen towards the circumferential direction, decreased exercise ability, and decreased function of the lung and right ventricle. The observed compromised mechanics in proximal pulmonary artery is consistent across multiple mouse models of accelerated ageing. Further, transcriptional changes in proximal pulmonary artery indicate that aging is associated with senescence of perivascular macrophages, adventitial fibroblasts, and medial smooth muscle cells. Older pulmonary arteries increase expression of genes associated with ECM turnover (including genes in the TGFβ pathway) and increased intercellular signaling amongst perivascular macrophages, fibroblasts and smooth muscle cells. Our results provide promising biomarkers of ageing for diagnosis and potential pathways and molecular targets for targeting anti-ageing therapies.

## Main

Age is the greatest risk factor for chronic pulmonary diseases in adults, including lung cancer, pulmonary fibrosis, and chronic obstructive pulmonary disease^1^. By extension, slowing the aging process should delay age-related diseases and prolong healthspan, also known as the geroscience hypothesis^2^. While it is well known that lung function declines in older adults, the etiology of this age-related decline remains unclear. Lung function can be defined by various metrics such as diffusion capacity, forced expiratory volume, or forced vital capacity, all of which diminish with age. The decline in these functional parameters associates with dyspnea, a common respiratory complaint of older adults and a geriatric syndrome that associates with worsening frailty and increased mortality. Yet, more than 30% of dyspnea in adults over the age of 60 is unexplained and often ascribed to “growing old” or “healthy aging^3^.” Therefore, a better understanding of the mechanisms underlying healthy aging of the cardiopulmonary system is clinically critical and important for prolonging lung function and healthspan of the older adults. Ventricular-vascular-ventilation coupling is central to lung function^4, 5^. Importantly, the large, proximal pulmonary artery (PA) which couples the pulmonary vascular bed with the right ventricle (RV), is a critical regulator of blood flow to the lungs and afterload to the right ventricle. The role of proximal vessel remodeling in age-related decline of pulmonary function is largely unexplored.

Age-related arterial stiffening is a recognized hallmark of aging for systemic arteries but has not been studied in detail in the pulmonary circulation^6–8^. In the aorta, structural stiffening is largely due to wall thickening resulting from an excessive deposition of collagen, often in the adventitia, in response to age-related fragmentation of elastin in the media^9–13^. Large artery stiffening associates with increased pulse wave velocity (PWV), a risk factor increasingly recognized for adverse, age-related cardiovascular outcomes^8, 14–19^. The positive feedback loop, encompassing wall thickening, arterial stiffening, and increasing pulse wave velocity, is hypothesized to lead to end organ dysfunction due in part to the associated increased penetration of pulsatility into the organ microcirculation^8, 12, 20^. It is well accepted that proximal PA geometry and properties are critical to regulating blood flow to the distal pulmonary circulation^21, 22^ despite accounting for a minority of the total compliance of the entire pulmonary vascular bed^23, 24^. Adverse vascular remodeling has been observed to increase with age in the distal pulmonary arteries^25^ and to associate with lung and RV dysfunction^5, 21, 26^. Nevertheless, the precise mechanisms by which the proximal PA may contribute to possible positive feedback via adverse remodeling and increased PWV remains poorly understood. We suggest, however, that stiffening of proximal PAs holds prognostic value in progressive pulmonary diseases such as pulmonary hypertension and chronic obstructive pulmonary disease^27–29^.

This concept is supported, in part, by our recent documentation that the human proximal PA stiffens with age, though not due to wall thickening or excessive collagen deposition as in the aorta^30^. Based on these findings, and known effects of aortic stiffening in systemic arteries, we hypothesize that age-related proximal PA remodeling associates with an increase in material stiffening and diminished capacity to store elastic energy of the arterial wall, which contributes to the decline in lung and RV function. A goal of this study was to determine a role, if any, of the proximal PA in coupling the RV and distal pulmonary vasculature as a cardiopulmonary unit in aging^4^. To test this hypothesis, we performed biomechanical experiments to quantify structural and functional characteristics of the pulmonary vasculature, lung, and heart from young (∼3 months) and old (24 months; natural aging model) adult male mice as well as from genetically modified mice that exhibit accelerated aging (Fibulin-5 null, Hutchinson-Gilford Progeria Syndrome) to assess similarities and differences in specific hallmarks of aging and age-related stressors in the cardiopulmonary system of aging models^31, 32^. We also quantified age-related changes in the mechanical properties of the lungs (using PFT), RV (using echocardiography) and proximal PA mechanics (using *ex vivo* biaxial mechanical testing). We narrowed our studies on the reduced ability of the proximal PA to store elastic energy and the effect this has on lung and RV function. This, in turn, focused our investigation on the role of extracellular matrix (ECM) in age-related stiffening of the pulmonary vasculature, a niche of “fibroaging” that has been largely unexplored^33^. We aimed to integrate multiple methods of investigating tissue, cellular, and molecular mechanisms to understand better any age-related remodeling of the ECM of the proximal pulmonary vasculature.

An overall goal of this study is to identify potential biomarkers of cardiopulmonary aging that may be translated to humans. Biomarkers are biological markers that can be quantified and measured to characterize normal or pathologic biological process as well as responses to therapeutic interventions^34, 35^. Therefore, in addition to characterizing healthy aging in mice, we hoped to identify measurable biological signs of age-related changes in the cardiopulmonary system of mice that may hold clinical meaning when measured in humans.

## Results

### Cardiopulmonary function is significantly decreased in older mice

We first evaluated overall cardiopulmonary function by measuring daily running distances. The older mice had significant cardiopulmonary impairment compared with younger mice (p = 0.02, Figure 1A). Cardiac function in older mice displayed significantly impaired RV contractility relative to younger mice (s’, p < 0.01). The RV is capable of homeometric (contractility) and heterometric (volume) adaptations to preserve stroke volume if faced with increased afterload, thus we compared RV free wall thickness and RA volume in young and old mice to determine if either or both of these adaptations occur with age^4^. We found a trend toward increased RV free wall thickening (p = 0.15) and a significant increase in RA dilatation (p = 0.04), as shown in Figure 1B, signs of RV compensation to delay RV heart failure. Additional right heart echocardiogram values are in Supplemental Table S1. We found that lungs in older mice expanded to significantly greater volumes, at similar amounts of inspiratory pressure, than lungs in young mice (p<0.05, Figure 1C). Further, older lungs had significantly increased compliance (0.094 mL/cmH_2_O vs 0.062 mL/cmH_2_O, p < 0.01) and increased inspiratory capacity (1.07 mL vs 0.66 mL, p<0.01).

**Figure 1.**
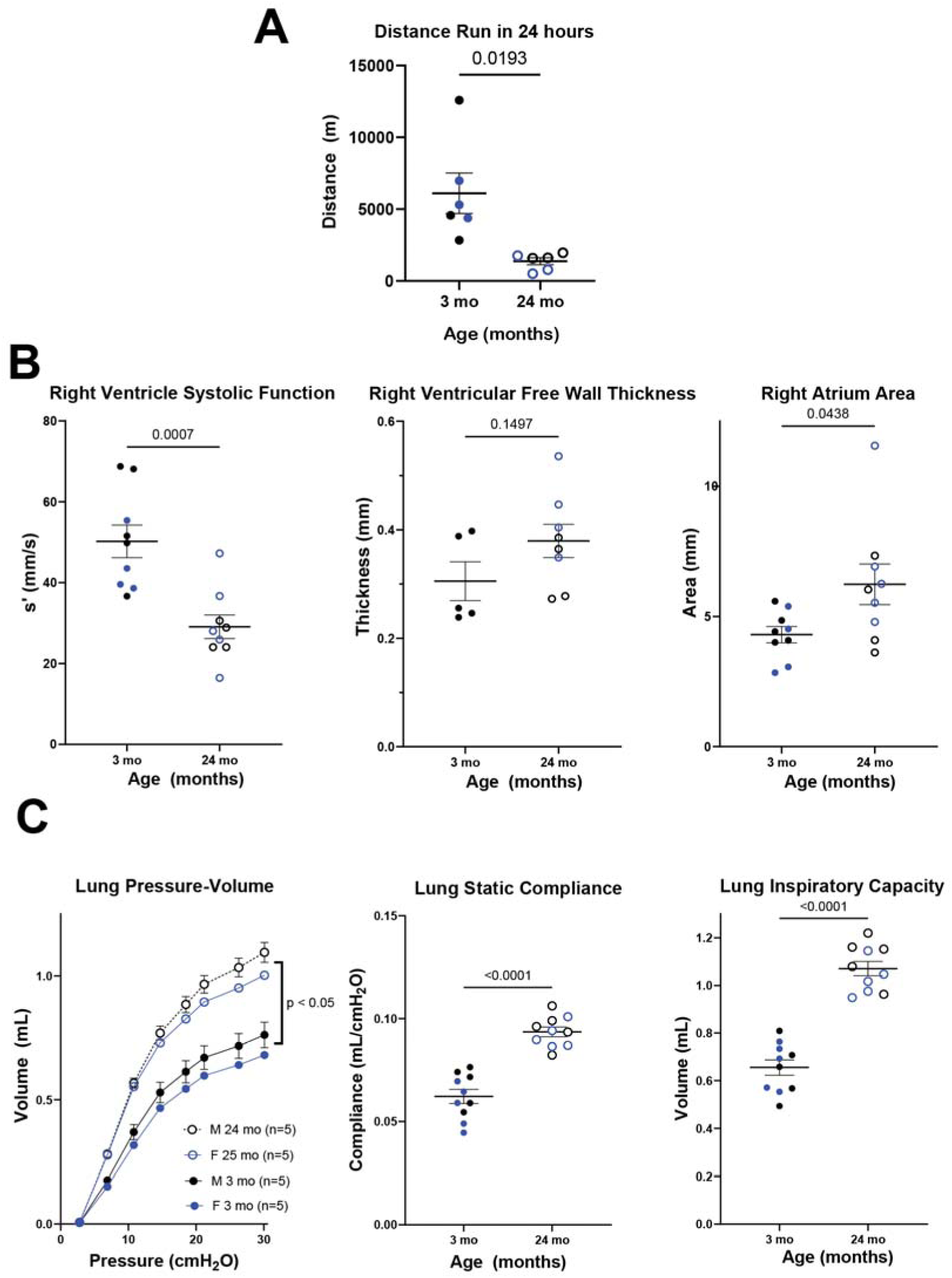
Overall cardiopulmonary function is significantly decreased in old compared to young mice. A) Average distance traveled per day (in meters) on running wheels over the span of a month for each mouse. B) Right ventricular (RV) systolic function in old mice was decreased significantly despite signs of RV compensation with a trend toward increased RV free wall thickness (hypertrophy) and significantly increased right atrium area, despite variability in aged subjects (suggesting chronic RV volume overload). C) Old mice have significantly increased lung volume for a given pressure increased static compliance and inspiratory capacity. Black = male, blue = female, solid circles = young mice, hollow circles = old mice.

### Old proximal PAs are stiffer and have less elastic stored energy than young proximal PA, which results in increased pulse wave velocity

We hypothesized that proximal PAs stiffen with age as observed in humans,^30^ thus we compared biomechanical differences between young and old mice. *Ex vivo* biomechanical testing revealed an aging-related leftward shift in the circumferential stress-stretch curve (Figure 2). Pressure increases with age in the right circulation^36^. Hence, we compared the proximal PA mechanics of the young mice at 15 mmHg with those of old mice at 25 mmHg, as these values represent *in vivo* loading conditions. We observed a reduced distension, despite an increase in inner radius, and a lower stored energy in the old mice (p = 0.0025), both consistent with a significant thickening of the aged wall (p = 0.04). We observed a significant increase in circumferential material stiffness (p< 0.0002) suggesting maladaptation in aging, though no significant change in axial stiffness, highlighting differential remodeling along the two primary in-plane directions. The decrease in stored energy was not significant when evaluated at age-appropriate mean pressure suggesting that the age-related remodeling was at least partially homeostatic (Supplemental Table S3). There are possibly opposite trends in male and female mice, where stored energy tends to decrease in older male mice but not female mice. Importantly, however, there was a significant reduction (p < 0.0001) in the distensibility of the proximal PA, noting that this metric combines information from both the circumferential and axial directions and thus it is a convenient metric of the passive function of the artery. We observed nearly a two-fold increase in PWV (p < 0.0001) associated with changes in proximal PA microstructure. Additional biomechanical data are provided in Supplemental Table S4.

**Figure 2.**
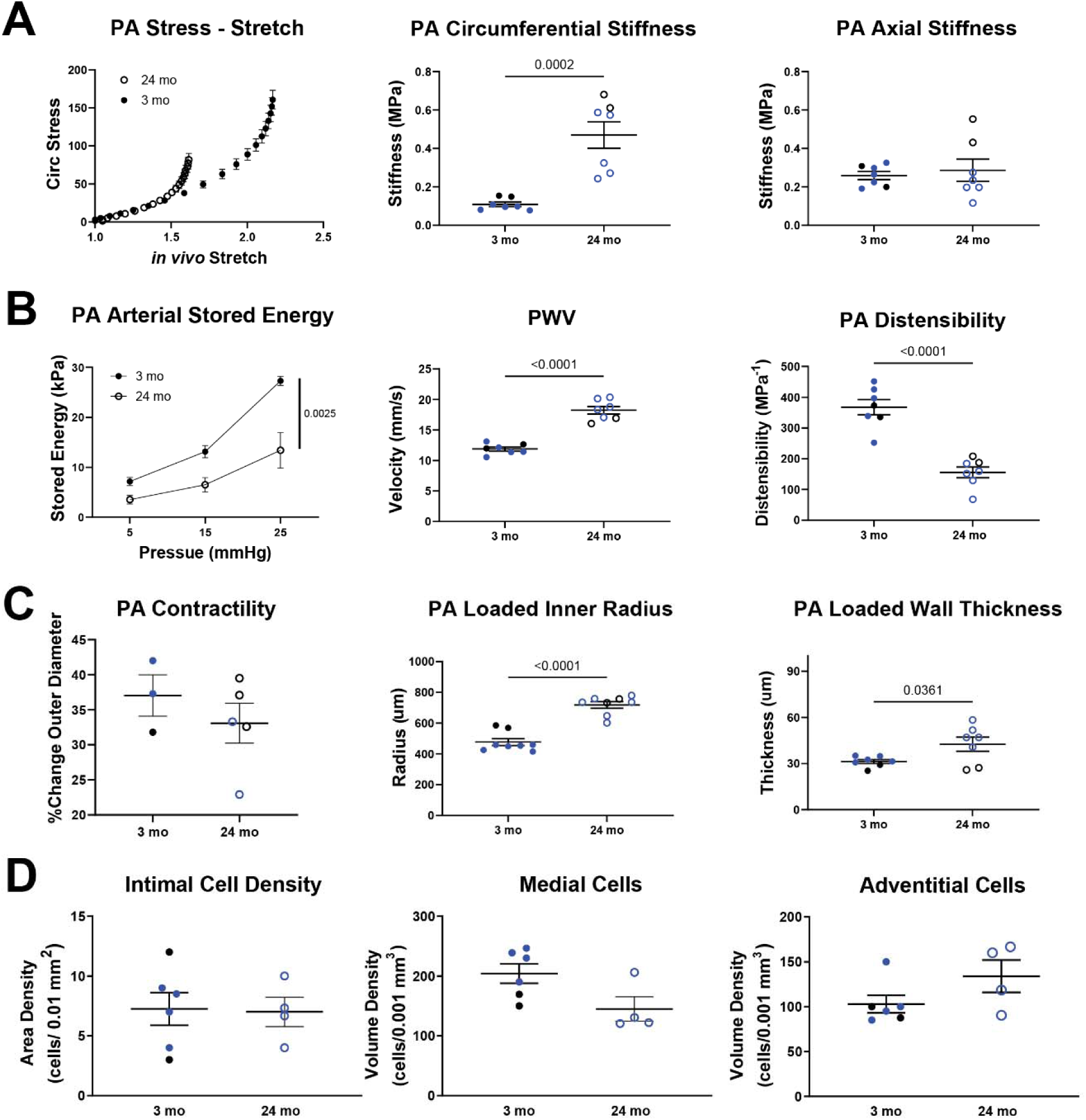
Old proximal PAs are stiffer and have less elastic stored energy than young proximal PA, which results in increased PWV. A) Stress in the walls of old proximal PAs increases at a greater rate than young proximal PAs as deformation increases, suggesting increased stiffness. We found that circumferential material stiffness of old PAs is greater than young PAs though axial stiffness is similar. B) Increased material stiffness of old PAs associates with decreased arterial stored energy, increased PWV, and decreased distensibility. C) The contractility of old PAs did not significantly decrease compared with young PAs. Significant age-related remodeled occurred, including nearly doubling in size of the vessel diameter and a modest increase in arterial wall thickness. D) There were no significant changes in the density of resident cells within the arterial wall when comparing old versus young samples to explain these changes in geometry or mechanical properties. Black = male, blue = female, solid = young, hollow = old.

### Proximal PA stiffness associates with decreased RV systolic function and lung mechanics

We previously found that biomechanical changes in the proximal PA (PA) of mice and humans associated with pulmonary and RV dysfunction, specifically increased stiffness of the proximal PA correlates with reduced measures of RV function, including ejection fraction, s’, and TAPSE^30, 37, 38^. Therefore, we performed linear regressions to compare changes in arterial stiffness with changes in lung and right heart mechanics. The decreased stored energy of the proximal PA of old mice associated with decreased *s’* (R^2^ = 0.53, p = 0.0006) and a trend toward increased RV free wall thickening (R^2^ = 0.18, p = 0.15), all signs of early RV dysfunction (Figure 3). Decreased stored energy of proximal PA of older mice also associated with increased lung compliance (R^2^ = 0.80, p < 0.05) and inspiratory capacity (R^2^ = 0.84, p< 0.05), all signs of impaired lung function (Figure 3, Supplemental Figure S2). We have previously associated these changes with loss of gas exchange units in the lungs of mice whose pulmonary arteries stiffened due to hypoxia^39^, and we found a similar association between proximal PA stiffening (with subsequent increase in PWV that is hypothesized to induce adverse remodeling/damage to cells of the pulmonary capillaries) and diffusion capacity of lungs of older mice (Supplemental Figure S2). Similar to the aorta,^9–11^ there was an increase in loaded thickness of the wall of the proximal PA (Figure 2), yet there was no significant increase in the proportion of adventitial collagen to overall collagen (medial + adventitial) in older mice compared with younger mice (p = 0.37). Next, we quantified the orientation of the adventitial collagen under in vivo loading conditions, noting that marked reorientation was observed previously due to hypoxia.^40^ The adventitial collagen in proximal PA of old mice had greater orientation toward the circumferential direction than younger arteries (old = 8.7 degrees ± 3.5 vs young = 2.0 ± 1.0, p = 0.0442, Figure 3). We also observed a trend of smooth muscle cell re-orientation toward the circumferential direction (old = 42.7 ± 4.3 vs young = 60.6 ± 8.8, p = 0.0727, Figure 3).

**Figure 3.**
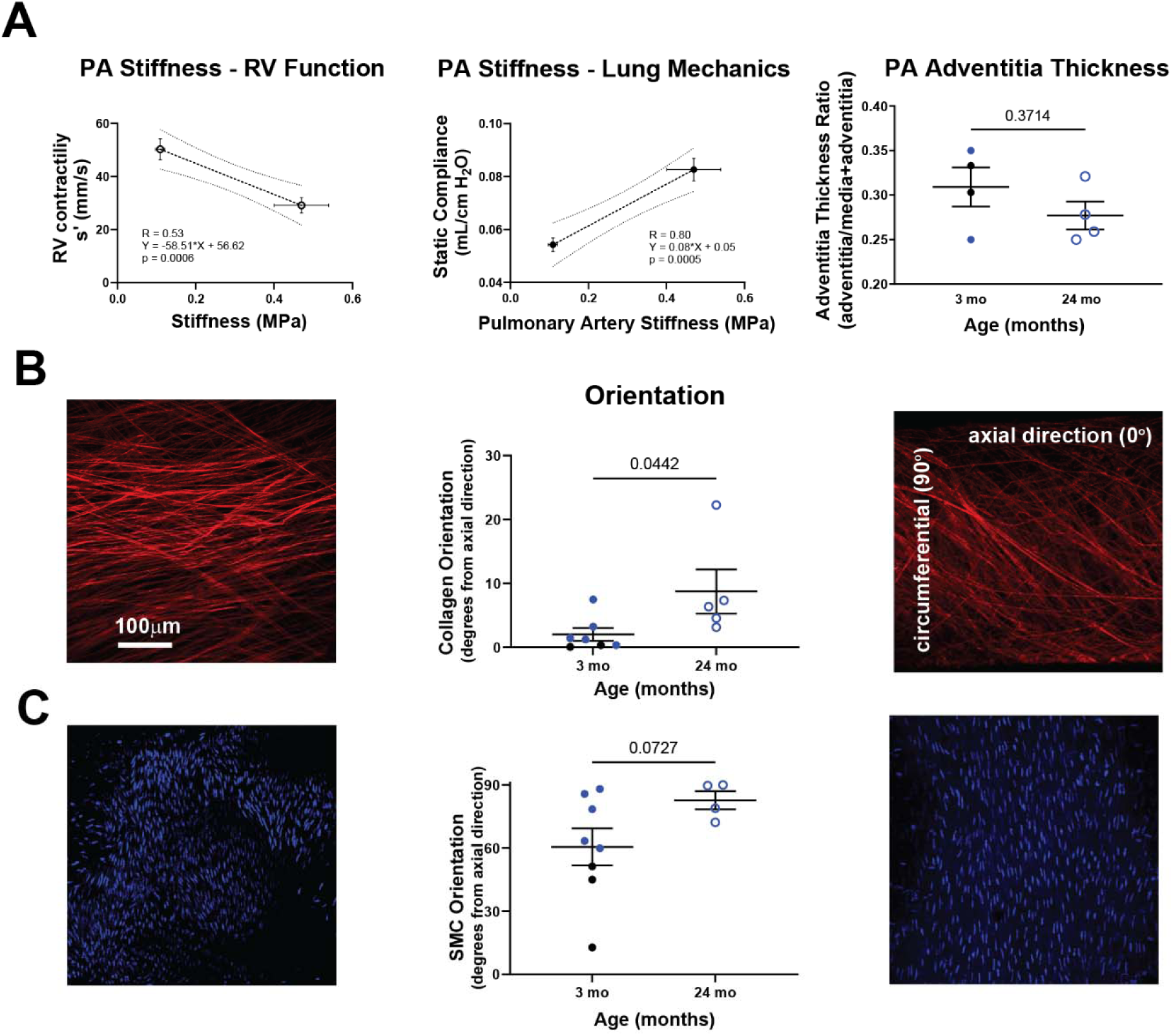
Proximal PA stiffness associates with declines in RV systolic function and lung mechanics, a result of microstructural remodeling of the adventitia rather than excessive collagen deposition and mere wall thickening. A) A linear regression of PA stiffening data with RV systolic function data (s’) and lung mechanics (static compliance) suggest that proximal PA stiffening associates with decline in RV and lung function. Additional timepoints are required to determine if this age-dependent relationship is truly linear or if a non-linear relationship exists. B) Microstructural analysis of the proximal PA wall shows that there is a significant re-orientation of collagen fibers from the axial toward the circumferential direction that associates with age-related proximal PA stiffening. C) SMCs in the media also shift their orientation toward the circumferential direction. Black = male, blue = female, young = solid, old = hollow.

### Age-associated transcriptional changes of the mouse proximal PA include ECM remodeling of the arterial adventitia

To investigate cellular and molecular mechanisms associated with the age-related remodeling observed in the proximal PA, we analyzed transcriptomic changes of resident pulmonary arterial cells and their intercellular communication. 20,950 cells were isolated from the proximal pulmonary arteries from n = 4 old mice (three males 20, 24, and 27 months and one female 28 months) and n = 4 young mice (two males and one female, each 3 months old), which revealed 18 different cell clusters (Supplemental Figure S3) including resident cells within the pulmonary arterial wall (ECs, SMCs, FBs, and MΦs), cells from surrounding tissue (large airway epithelial cells), and cells that may be associated with the arterial wall or surrounding tissue such as lymph nodes or thymus (B cells, T cells). There were no significant changes in the proportions of resident cells of young and old mice (Supplemental Figure S3).

The differential gene expression of adventitial FBs, SMCs, MΦs, and ECs revealed differential expression of 4,882 genes that positively correlated with age-related differences between young and old mice and 7,005 genes that negatively correlated with key biological processes and molecular pathways (Figure 4, Supplemental Figure S4). Common biological processes across multiple cell types in old cells include morphogenesis of cardiovascular structures (FB GO:0003007, GO:0003143; SMC GO:0060977; EC GO:0048514), cell cycling and proliferation (FB GO:2000134; SMC GO:0030308; MΦ GO:0016567, GO:0042981; EC GO:1901978), and cell motility and matrix interactions (FB GO:0007626; SMC GO:0035426, GO:0030335; EC GO:0120180, GO:0010810, GO:0010595). FBs and SMCs expressed several genes that associate with ECM turnover (Bmp6, Smad3, Smad6, Adamts12, Col8a1, Fgf1,Smad4, Fzd4,Ntn4)^41^. FBs from old cells had decreased expression of genes associated with aerobic respiration and carbon metabolism, suggesting that old FBs may rely on alternative sources of cellular energy such as lipid or fatty acid metabolism. We found increased expression of genes associated with inter- and intra-cellular signaling (ECM-SMC signaling; mTOR, IL-10, and Wnt signaling in SMCs; retinoic acid, cytokine, and IFNγ, and phosphorylation mediated signaling in perivascular macrophages; VEGF, cytokine and IFNγ signaling in ECs).

**Figure 4.**
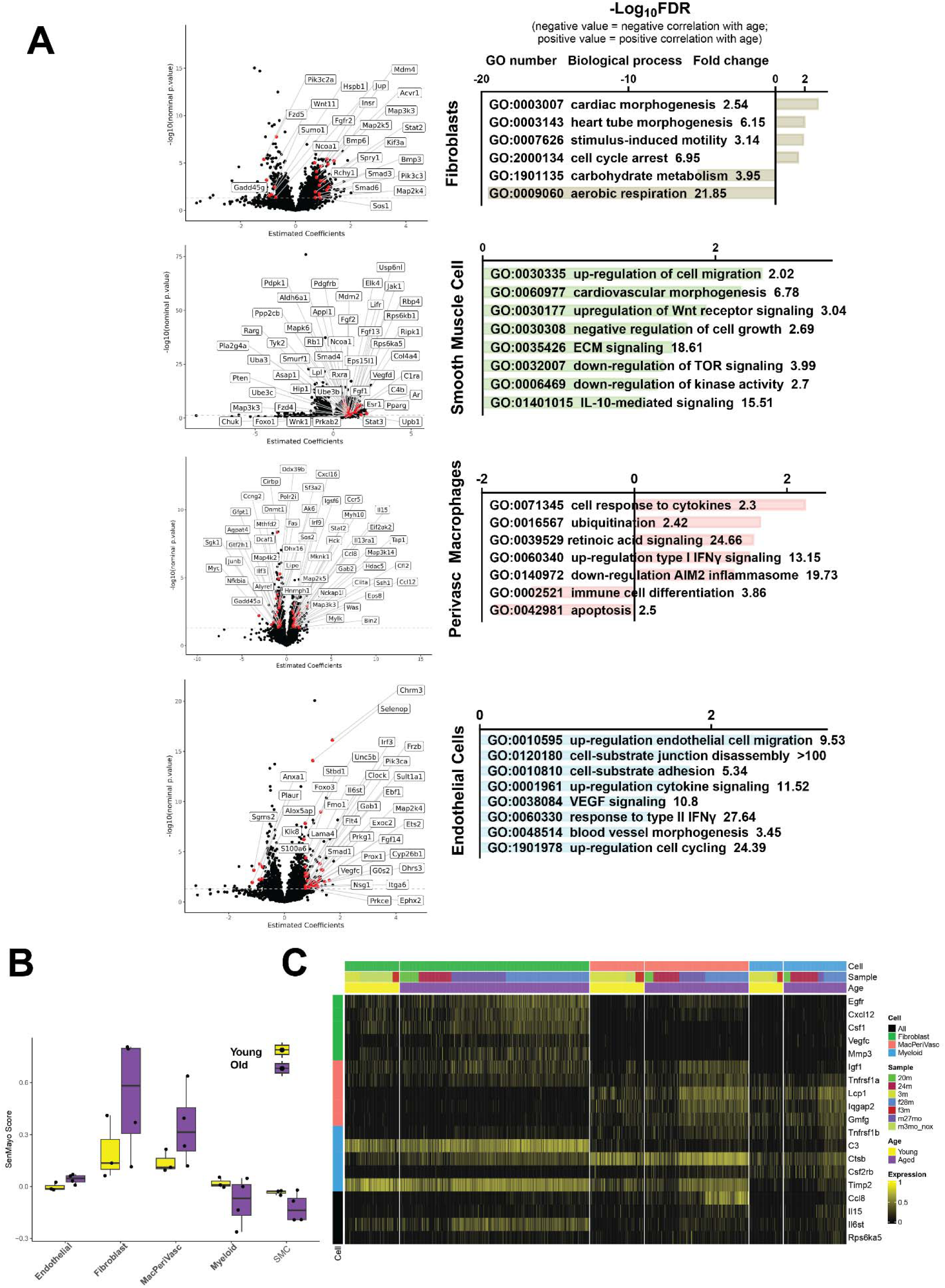
Single Cell RNA sequencing data identifies DEGs and biological processes that associate with ECM remodeling of the arterial wall with significant increased senescence of perivascular macrophages and fibroblasts. A) Volcano plots of the DEG analysis (left panels) demonstrate changes in gene expression between old and young PAs. Select DEGs were highlighted for each resident cell type. GO term enrichment analysis (right panels) of the top DEGs showed enrichment in age-related biological processes that provide cellular-molecular insights into age-related changes of the ECM in proximal PAs. B) SenMayo scores for each cell, grouped by cell type and split by age group. SenMayo scores averaged per sample. C) Heatmap showing SenMayo genes that are increased in aged resident pulmonary artery cell types. The top 5 SenMayo genes are plotted for each cell type.

### Aging of the cardiopulmonary system associates with senescence of perivascular M***Φ***s, FBs, and SMCs that reside in the walls of the proximal PA

Based on observed alterations in cellular proliferation, metabolism, and cytokines, we hypothesized that senescence plays an important role in age-related changes. To further explore the role of senescence in aging of the proximal PA, we used the SenMayo senescence signature^42^. We calculated a senescence score for all cells in the dataset and designated cells with a score exceeding 0.25 indicating senescence. We identified 352 cells expressing this signature throughout our population of resident cells (Supplemental Figure S3). Additionally, a higher proportion of cells from old mice expressed canonical senescence markers such as CDKN1A (p21) (26.3% vs 21.7%) and CDKN2A (0.8% vs 0.4%).

Given the senescent signature in the overall cell population, we analyzed the senescent signature of resident ECs, SMCs, and FBs within the wall of the proximal PA, and subdivided the myeloid cell population into three separate phenotypes of MΦs: we reserved “myeloid” for recruited cells such as circulating monocytes (characterized by genes *Ccr2, Cd24a*, Figure 4), MΦs associated with lung parenchyma (MacLung, *Chil3, Fabp5*), and resident perivascular MΦs (MacPeriVasc, *C1qa, Pf4* comparable to *FOLR2+* MΦs in humans). Resident cells of the proximal PA of old mice had significantly higher senescence scores than younger mice (adventitial FBs 0.248 vs 0.104, p < 0.05; perivascular MΦs 0.163 vs 0.093, p < 0.05 Figure 4). To determine which genes were driving the differences in senescence score, we examined differential expression of SenMayo genes for each cell type. Among FBs, perivascular MΦs, and myeloid cells, only 4 of the 125 SenMayo genes were consistently increased across cell types (*Ccl8*, *Il15*, *Il6st*, *Rps6ka5*; Figure 4). SenMayo genes that were increased exclusively in FBs included *Egfr*, *Cxcl12*, *Csf1*, *Vegfc*, and *Mmp3*. In contrast, SenMayo genes that increased in MΦs, and myeloid cells included pro-inflammatory molecules such as *Tnfrsf1a*, *C3*, and *Ctsb*.

### Older proximal pulmonary arteries exhibit altered intercellular communication patterns associated with ECM remodeling and inflammation

Based on changes in gene expression associated with signal transduction, we hypothesized that altered intercellular interactions may regulate FBs activity and play a key role in promoting adventitial homeostasis. Therefore, to further investigate the cellular-molecular nature of adventitial remodeling that contributes to age-related stiffening of the proximal PA, we investigated possible intercellular communications of recruited myeloid and resident cells. We found the strongest intercellular communication amongst resident perivascular MΦs, adventitial FBs, and SMCs (Figure 5), cell types associated with increased senescent signature in tissue from older mice. We identified gene signaling pairs, elaborated below, that were significantly increased between the different types of MΦs and FBs in the older pulmonary arteries compared to younger arteries. Notably, there was considerable overlap in the cell signaling changes from perivascular MΦs to FBs and SMCs (122 signaling mechanisms out of 270). Mechanisms that increased with aging included TGFβ and cytokine signaling mechanisms (*Tgfb1-Itgb8, Tgfb1-Itgb6, Tgfb1-Tgfbr1, Tgfa-Eng, Tgfb1-Acvrl1, Bmp2-Acvr2b, Bmp2-Bmpr1a, Bmp2-Acvr2a, Bmp2-Bmpr2, Il18-Il1rl2, Ccl2-Ackr2, Cxcl12-Itgb1*), PI3k-Akt and growth factor pathways (*Igf1-Insr, Nrg2-Erbb2, Pdgfc-Pdfrb, Igf1-Igf1r, Fgf9-Fgfr1, Egf-Egfr*), ECM organization (*Mmp9-Ephb2, Adam9-Itgb5, Adam9-Itgb1, Adam9-Itgav, Ecm1-Cachd1*), and response to hypoxia (*Vegfb-Nrp1*). Similar increases in cytokine signaling and growth factor pathways were observed in outgoing myeloid cell signaling to FBs and SMCs (complete table in Supplementary Table S7). Intercellular outgoing signaling from the endothelium to the myeloid cells was also characterized (Figure 5). Analysis of signaling from ECs to SMCs revealed an age-associated increase in signaling mechanisms related to ECM-receptor interactions and focal adhesion (*Col4a1-Itgb8, Col4a1-Itgav, Fgf9-Fgfr1, Lama3-Sdc2, Lama4-Itgav*).

**Figure 5.**
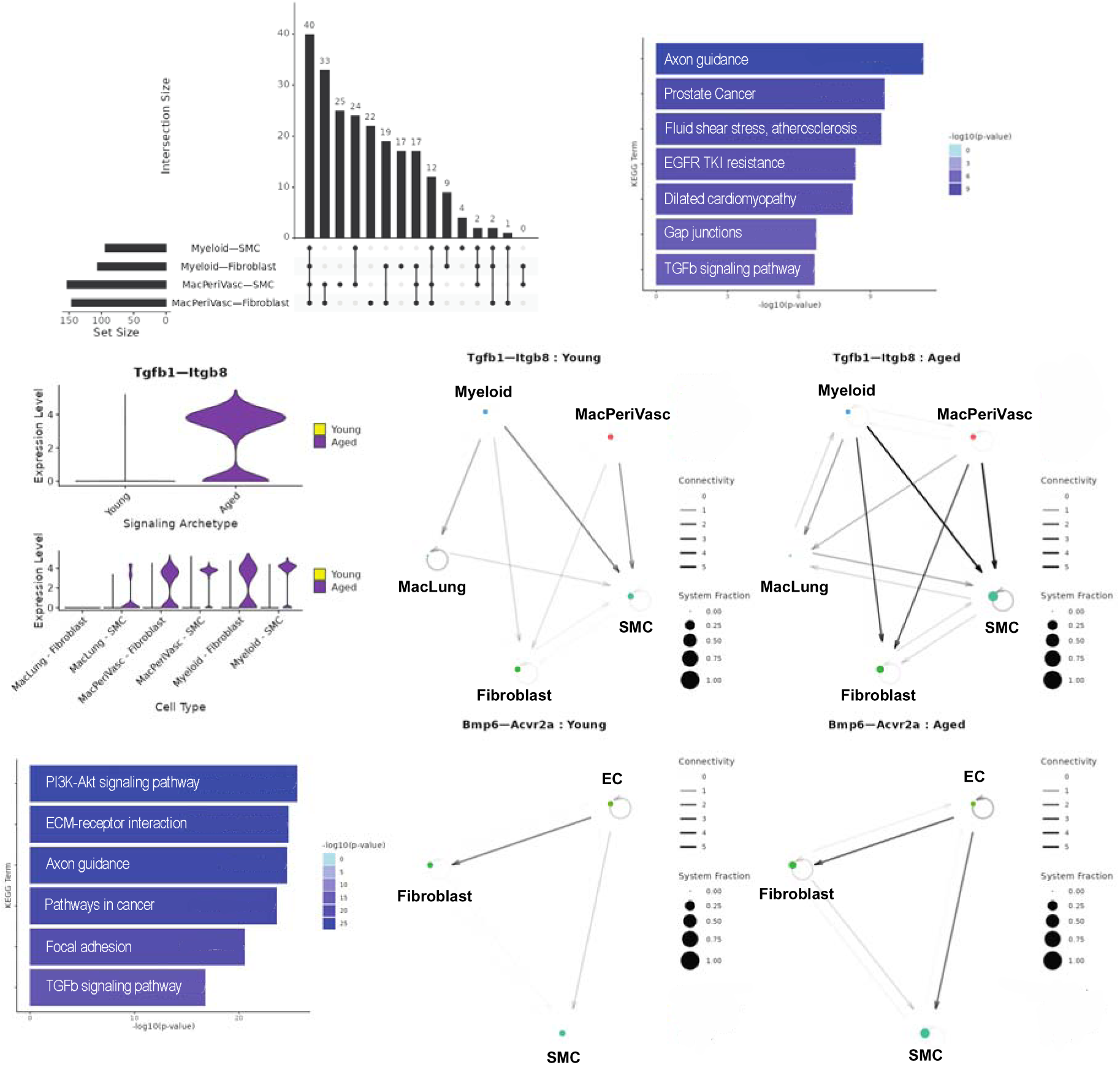
NICHES reveals age-related changes in myeloid-to-mesenchymal cell signaling. (A) Upset plot showing overlap between cell signaling mechanisms that are differentially expressed in aging. (B) KEGG enrichment terms and associated p-values for signaling mechanisms that are increased with age for MacPeriVasc to FB and SMC signaling. (C) Expression level of Tgfb1-Itgb8 in old versus young samples. (D) Circuit plot showing Tgfb1-Itgb8 signaling in young and aged cell types. Node size is proportional to cellular fraction and edge width is proportional to connectivity. (E) KEGG enrichment terms and associated p-values for signaling mechanisms that are increased with age for Endothelial to SMC signaling. (F) Circuit plot showing Bmp6-Acvr2a signaling in young and aged cell types.

### Proximal PAs of progeroid mice are significantly stiffer, less distensible and exhibit greater pulse wave velocity

To investigate additional factors of aging and their relationship with proximal PA remodeling, we used genetically modified mice to quantify mechanical changes for two forms of accelerated aging: impaired elastogenesis to model loss of elastin in large arteries commonly associated with aging (*Fbln5^-/-^*) and nuclear stress associated with accelerated aging as a result of Hutchinson Guilford Progeria Syndrome (HGPS; *Lmna^G609G/G609G^*). Age-related changes in elastic fibers are observed naturally over decades in humans but not in WT mice due to their relatively short lifespans compared to the half-life of vascular elastin (∼50 years). Consistent with natural aging, biomechanical testing of the proximal PA from these accelerated aging models also revealed leftward shifts in the circumferential stress-stretch behaviors. Comparisons at *in vivo* relevant pressures (15 mmHg for young and 25 mmHg for aged groups) revealed a significant reduction in elastic stored energy in the progeria mice (p = 0.0029, Figure 6) though proximal PA of 3 mo old mice with impaired elastogenesis remained similar to young C57BL/6J mice (p = 0.999). In particular, the ability of the pulmonary arteries to store elastic energy ordered, across all groups, as 3 mo C57BL/6J > 3 mo *Fbln5^-/-^* > 24 mo C57BL/6J > 6 mo HPGS. Similar to the 24 mo group, the distensibility of the HGPS and *Fbln5^-/-^* mice were significantly reduced, (p< 0.001) with increases in pulse wave velocity (p< 0.05, Figure 6).

**Figure 6.**
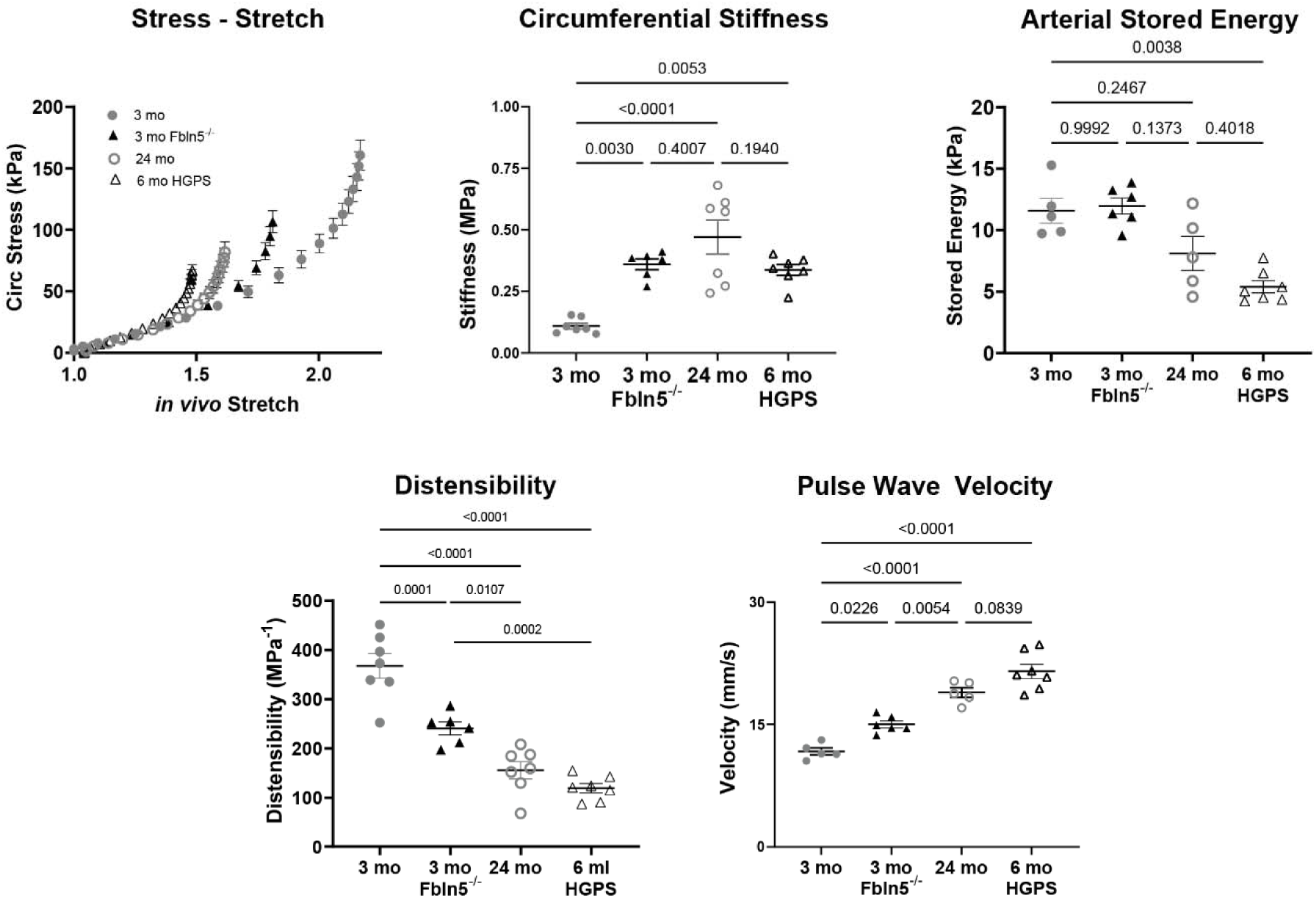
Proximal PAs of progeroid mice (3 mo Fbln5-/-, 6 mo HPGS) resemble proximal PAs from old C57BL6 mice and are significantly stiffer, less distensible and greater PWV than young C57BL6 mice. A) Stress in the walls of old and progeroid proximal PAs increases at a greater rate than young proximal PAs. Circumferential material stiffness of the arterial walls of old, 3 mo Fbln5-/- and progeroid proximal PAs is significantly increased compared with young proximal PAs. Similarly, distensibility significantly decreases in old, 3 mo Fbln5-/- and progeroid PAs and PWV significantly increases.

## Discussion

Based on clinical observations that the cardiopulmonary system declines with aging, we hypothesized that aging alters the underlying mechanical properties. To test this hypothesis, we compared cardiopulmonary mechanics in the RV, proximal PA, and lungs of young and old mice. We found that the proximal PA plays an important role in age-related changes – it stiffens with increasing age which associates with decreased mobility, diminished RV contractility, and diminished lung function. Therefore, stiffening of the proximal PA may be a biomarker of aging of the cardiopulmonary system in mice. We also found that such aging associated with senescence of perivascular MΦs, adventitial FBs, and medial SMCs within the proximal PAs. Older pulmonary arteries increase expression of genes associated with ECM turnover (including genes in the TGFβ pathway, proteases such as *Adamts12*, and *Col8a1*) and increased intercellular signaling amongst perivascular MΦs, FBs, and SMCs.

### The proximal pulmonary artery plays an important role in the aging cardiopulmonary system

There is a paucity of literature on the mechanisms of aging that account for the concomitant declines in function of both the right heart and lung, a gap in knowledge that we believe this study helps fill. We previously related increased structural stiffness of the proximal PA in humans, reduced RV and lung function. We also hypothesized that material stiffness of the proximal PA increases with age^30^. This study on mice corroborates age-related increased proximal PA material stiffness associates with reduced RV and lung function, similar to phenomena in the systemic circulation^12, 19, 43^. We showed that maladaptations of the proximal PA also associate with reduced RV and lung dysfunction in the mouse as a result of chronic hypoxic exposure or loss of elastin proteostasis^38, 40, 44^. Our findings suggest proximal PA stiffening contributes to age-related functional decline of the RV and lungs^36, 45–50^.

Decreased distensibility of the proximal PA increases PWV, which is known to negatively affect arterioles and capillaries in end-organs due to increased pulsatile pressure variations^51^. Repeated injury of the microcirculation can lead to end-organ dysfunction^14, 15, 17, 51^. In the pulmonary circulation, this negatively affects ECs lining capillaries that surround lung alveoli, thus impairing the fundamental units of gas exchange in the body. Reflected waves due to increased arterial wall stiffening also negatively impacts ventricular contractility^8, 12, 52^. Although decreased distensibility of the proximal PA has been associated with decreased exercise ability in individuals with pulmonary hypertension^53, 54^, we believe that this is the first study to find this relationship in aging. Further, a recent study associated decreasing axial (or longitudinal) strain with increasing severity of pulmonary hypertension in humans^29^. Our study further extends this finding to aging and provides mechanistic insights to the phenomenon.

Age-related changes leading to aortic stiffness and associated hemodynamic sequelae (thickening of the wall, decreased *in vivo* axial stretch, and loss of arterial elastic stored energy) are well described in humans and mice^10, 12, 55^. We found similar age-related remodeling in the proximal PA. Whereas age-related mechanical changes of the aorta associate with increases in collagen content and fiber thickness, we found that collagen degradation and re-deposition in a direction oriented more toward the circumferential direction in the proximal PA drive the observed age-related mechanical changes. The resulting increases in pulse pressure, ventricular hypertrophy, and diastolic dysfunction appear to be the same in both the pulmonary and systemic circulation. The lack of age-related changes in elastin, likely due to the short lifespan of mice compared to humans, suggest that other mechanisms of aging are at play providing potential therapeutic targets to prolong cardiopulmonary healthspan.

We also previously found that induced elevation of pulmonary arterial pressure (by chronic hypoxia) has profound effects arterial wall remodeling, resulting in similar degrees of arterial wall stiffening, decreased stored energy, and collagen fiber re-orientation after five weeks of hypoxia as what we observe in two years of aging. Adverse tissue-level remodeling of the proximal PAs has been prevented when nifedipine preserved blood pressure during hypoxic exposure^56^, suggesting that the vascular remodeling was in response to the pressure rise. However, several reports suggest that hypoxia stimulates early monocyte / macrophage activity^57^ thus driving pulmonary vascular remodeling and the development of pulmonary hypertension^58–60^. We found that perivascular MΦs are important to age-related remodeling of the proximal PA, potentially identifying this cell type as a therapeutic target for age-related changes in the cardiopulmonary system.

Stiffer proximal PAs and associated decreases in stored energy were also evident in multiple progeroid murine models, including HGPS and compromised elastogenesis. Mutagenically-induced cell stress in progeria mice mimics advanced-aging phenotype of the proximal PA. Our group has previously demonstrated that mice with Hutchinson-Gilford progeria syndrome have accelerated aortic stiffening with left heart dysfunction^61^. To the authors’ knowledge the present study is the first report of accelerated stiffening of the pulmonary vasculature in Hutchinson-Gilford progeria syndrome.

Stiffer large arteries have an impaired ability to store elastic energy during systolic distension, which compromises the ability of the artery to augment flow during diastole that, in turn, is expected to associate with impaired lung perfusion, lung and RV function, and overall poorer cardiopulmonary performance, all of which we observed in this study. Non-invasive measures of the proximal PAs are possible through cardiac MRI and echocardiography and therefore may be a non-invasive biomarker for biological aging of the cardiopulmonary system. Proximal PA stiffening may be a potential therapeutic target to delay aging, prolong healthspan, and prevent age-associated chronic diseases of the cardiopulmonary system^34, 35^.

### Aging of proximal PA in mice associates with ECM turnover, cellular senescence in the arterial wall, and altered intercellular signaling

Senescence plays a critical role in age-related changes in the physiology of the lungs^47, 62, 63^. Yet, there is significant variance between senescence of various tissues and organs^64^. We therefore used a generalized, transcriptomic signature of senescence to identify potential biological roles of senescent cells in age-related changes of the proximal PA^65, 66^, which to the best of our knowledge, has not been done before. Our findings are nevertheless consistent with previous studies of the whole human lung (not including the proximal PAs) that found increased prevalence of senescence in immune cells, specifically in monocytes and alveolar MΦs^67^. We also found increased senescence of perivascular MΦs as well as FBs and SMCs that reside within the wall of extra-lobar arteries. Senescence and age-related transcriptomic changes of these cells associated with ECM remodeling (degradation by proteases and collagen deposition in orientations more toward the circumferential direction), a novel, mechanistic result.

In our prior studies of aging of the human lung, including the distal pulmonary arteries, we found that genes associated with senescence expressed by FBs included the senescent gene *IGF1* as well many genes for collagens and matrix metalloproteinases, suggesting an association between senescence of lung FBs and dysregulation of ECM associated with aging of the lung^68–70^. Therefore, there may be similar associations between senescence of SMCs and FBs of the proximal and distal pulmonary arteries involving IGF signaling and adventitial ECM remodeling. We found increased senescence of ECs in the proximal PAs of older mice similar to findings of increased senescence of ECs in distal pulmonary arteries of the aged human lung^68, 69^ that others have associated with age-related lung diseases such as COPD^71^. We conclude that senescence plays an important role in age-related remodeling of both the proximal and distal pulmonary arteries of the lung, and perivascular MΦs play an important role in regulating age-related remodeling of the proximal vessels in mice. The latter corroborates our findings that perivascular MΦs play a critical role in regulating proximal PA remodeling in mice exposed to chronic hypoxia^39^.

Age-related proximal PA stiffening shares similarities with stiffening due to hypoxic exposure though at a much slower rate and by different cellular and molecular mechanisms. We found that increased material stiffness of the wall due to aging or chronic hypoxia associated with re-orientation of the adventitial collagen fibers toward the circumferential direction^39, 40^. We also found that these changes result from altered intercellular signaling amongst SMCs, FBs, and MΦs^40, 72^. A major difference between age-related stiffening of the proximal PA compared to chronic hypoxia is the preserved contractility of SMCs in older mice despite reduced cell numbers. In hypoxic environments, SMCs undergo a metabolic shift from aerobic respiration to glycolysis, which significantly alters intercellular communications amongst SMCs, FBs, and MΦs^39^. mTOR inhibition with rapamycin showed limited success in preserving SMC contractility and preventing proximal PA stiffening in hypoxia,^39^ but mTOR inhibition may prove more beneficial in delaying age-related stiffening. It remains unclear if all SMCs in the media change phenotypes or only certain proportions. Our transcriptomic data revealed intra- and intercellular signaling changes that suggests perivascular MΦs play an important role in regulating remodeling of the ECM by adventitial FBs and medial SMCs, similar to results that we observed in response to chronic hypoxia^39, 40^. These findings mirror studies of the aging lung (not including the proximal PA) that reported age-associated changes in expression of ECM components including collagens, and laminins^73, 74^. In this study we found that transcriptional changes that associate with ECM remodeling of the proximal PA also associate with an increased senescence signature in perivascular MΦs and FB populations and increased signaling amongst other resident cell types in the arterial wall.

### Hallmarks of cardiopulmonary aging include altered biomechanical properties

We identified key hallmarks of aging^31^ in the cardiopulmonary system of mice, compromised proteostasis of collagen, increased senescence of MΦs and FBs, genomic instability, and alterations in intercellular communication amongst cells that reside in the walls of the proximal pulmonary artery. These age-related changes provide mechanistic insights into stiffening of the proximal pulmonary artery and reduced function of the RV and lungs. This supports a recently proposed addition of altered mechanical properties as a hallmark of aging^32^. The mechanisms underlying age-related changes in mechanics of the murine pulmonary artery appear to be independent of both elastic fiber damage and fibrosis—the excessive deposition of collagen that one expects in the adventitia of older, stiffer aortas^10, 75^. Rather, the remodeling of collagen within the adventitia resulted mainly from changes in microstructure rather than changes in quantity.

The loss of proteostasis of adventitial collagen suggests that cellular and molecular mechanisms are key to understanding age-related pulmonary artery remodeling. Therefore, we focused on the roles of FBs and MΦs regulation of FBs, which are mostly found in the adventitia. There were no significant changes in the proportions of these cell types when comparing resident cell populations from old and young pulmonary arteries, but there were signs of altered gene expression and altered intercellular communications that associate with remodeling of the ECM (TGFβ mediated) and inflammatory processes associated with aging (Il6). We also found both higher proportions of cells expressing key senescence genes CDKN1A and CDKN2A and increased senescence transcriptional signatures in the cells that resided in the walls of proximal pulmonary artery (MΦs, ECs, SMCs, and FBs). Three major myeloid phenotypes reside in the walls of proximal pulmonary arteries, and there is increased intercellular signaling between perivascular MΦs and stromal cells (FBs, SMCs). Upregulation of TGFβ and cytokine signaling in old PAs suggests dysregulation of cytokine and growth factor signaling contribute to age-related structural changes structural in the proximal PA, as described regarding senescence and other pathologies such as cancer^76, 77^. These findings highlight the importance of intercellular communications in adventitial collagen remodeling beyond mere transcriptional changes.

### Limitations

Our study is limited by several factors. First, we compared cells and tissues from young (∼3 months) and older (∼24 months) mice. While this proved to be sufficient to identify age-related changes in the cardiopulmonary system, it is insufficient to investigate mechanisms of the progressive process of aging. Future studies should sample ages well distributed between the two ages we considered. Further, the variation in the populations of cell types dissociated from the proximal PAs appears to be larger than the effect size that we can measure. The tortuosity of the *Fbln5^-/-^* mouse aorta and fragility and small size of HGPS mice posed challenges to *in vivo* pressure measurements. Many mechanical measurements are pressure-specific, and lack of *in vivo* measurements is a drawback of the study. This was ameliorated by using representative *in vivo* pressure, based on literature data. Age-related changes in the aorta appear to be more pronounced than induced hypertension and independent of sex as a biological variable,^11^ however, we found some post-hoc differences in proximal PA mechanics that may be sex dependent, such as elastic stored energy of the arterial wall. This study is insufficiently powered to determine this difference using sex as a biological variable.

We quantitatively explored the expression of senescence gene signatures in resident cells by designating each cell a senescence score based on the SenMayo gene list. SenMayo includes many senescence-associated secretory phenotype (SASP) genes and has been validated experimentally in human and mouse models in a variety of cell and tissue types, though not specifically for the proximal PA. While overall senescence was noted in MΦs and FBs, distinct SenMayo genes increased in each cell type, suggesting heterogeneity in senescence transcriptional programs^78^. Our results suggest that MΦs and myeloid cells exhibited greater expression of proinflammatory SASP genes in aging, whereas FBs had greater expression of genes in developmental pathways. The heterogeneity of senescence programs in different cell types within this tissue is an area that warrants further study.

### Conclusions

This study presents two major finding that warrant further investigation. First, it is critical to determine if the hallmarks of aging identified in the proximal PAs of mice are valid in humans. In line with the geroscience hypothesis, our mechanistic insights, if present in humans, may provide therapeutic targets to delay age-related declines in RV and lung functions and mitigate age-related disability and diseases^2^. Second, it would be beneficial to evaluate whether mechanical metrics such as distensibility, stiffness, stored energy, and PWV can serve as reliable biomarkers of aging of the cardiopulmonary system in humans. Success in these areas may reduce frailty and prolong healthspan of older individuals.

## Methods

### Mice

The study was approved by the Yale University Institutional Animal Care and Use Committee. Young (∼3-5 months; labeled as 3mo in figures) and old (24 months), female and male C57BL/6J WT mice (young and old mice from Jackson Laboratory, Bar Harbor, ME and old mice from the NIA aging mice colony) were housed in an antigen-free and virus-free animal care facility under a 12-hour light and dark cycle. Mice of similar strains from different environments may differ in phenotype^79, 80^. To minimize environmental differences, mice from Jackson Laboratory were inbred locally and some were allowed to age locally. Mice from the NIA Aging Colony were obtained at 20 months of age and aged further locally. Mice were fed a standard rodent chow and had free access to water. For *ex vivo* testing and transcriptomic analyses, animals were euthanized either with an overdose of urethane or Beuthanasia-D by intraperitoneal injection followed by exsanguination and harvest of the hearts, lungs, and pulmonary arteries. To explore mechanisms further, we used two genetically modified mice that display accelerated vascular aging: fibulin 5 null (*Fbln5^-/-^*), which are deficient in the elastin-associated glycoprotein fibulin 5^81, 82^, and *Lmna^G609G/G609G^*, which harbor a mutated form of the nuclear envelop scaffolding protein lamin-A that results in the Hutchinson-Gilford Progeria Syndrome^61, 83^.

### Voluntary Exercise

We measured the daily running distance of mice using in-cage running wheels (Actimetrics Wireless Low-profile Running Wheel Model ACE-557-WLP) with ClockLab Data Collection Software from Lafayette Instrument. These devices are designed to not interfere with normal housing conditions. We allowed the mice two days to acclimate to their cage with the running wheel and measured the distance run in meters over 24 hours on the third day. All mice participated.

### Biomechanical Measurement and Analysis

Specimens were excised from the main pulmonary artery to the first branch of the right (RPA) and left (LPA) pulmonary artery and prepared as described previously^84^. After flushing blood with a Hanks buffered saline solution^85^, perivascular tissue and fat were gently removed, and the LPA, ligamentum arteriosum. and small branch vessels were ligated with suture. The RPA was cannulated on custom glass micropipettes and secured with ligatures beyond the main pulmonary artery bifurcation and the first branch of the pulmonary at the other end. The specimen was submerged in Hanks buffered saline solution at room temperature to eliminate smooth muscle contractility and biaxially tested using a custom computer-controlled testing device^86^. To promote reproducibility and rigor, we used the same seven passive testing protocols as described previously,^87^ namely, pressure-distension tests at three fixed axial lengths (1.05, 1 and 0.95 times the *in vivo* length) and axial force-extension tests at four fixed constant pressures (5, 15, 25 and 40 mmHg). Diameter was measured using a videoscope, length was imposed with a micro-stepper motor, and pressure and force were measured with standard transducers. We used a 2-D formulation to model the passive mechanical behavior; the residual stresses tend to homogenize the stress field, thereby rendering mean values as good estimates of overall wall stress^88^. We modeled the wall using a hyperelastic constitutive formulation consisting of neoHookean term to capture elastin-dominated contributions and four-fiber families with Fung-like exponential behaviors to capture collagen and smooth muscle contributions. Importantly, from this relation we can derive clinically and mechanobiologically relevant quantities such as biaxial wall stress and stiffness at different loads or deformations of interest. Details of the parameter estimation of these passive constitutive functions can be found elsewhere^84, 87^. Methods for measuring contractility in a biaxial setup can be found in our previous work^40, 84^. Here we report contractility at a constant distending pressure of 15 mmHg and subject-specific *in vivo* stretch given vasoactive stimulation with 100 mM potassium chloride (KCl) and 100 mM phenylephrine (PE) within an oxygenated Krebs physiological solution maintained at 37°C and 7.4 pH.

PWV is an integrated measure of the structural stiffness of an artery; it depends on both the geometry and material properties and can be well-approximated based on vessel distensibility *D*, namely

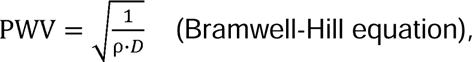

where *ρ* is the mass density of the contained fluid (∼1050 kg·m^-3^) and *D* (in Pa^-1^or kg^-1^m·s^2^) is defined by the normalized change in arterial inner diameter from end-systole to end-diastole divided by the change in end-systolic and end-diastolic pressures.^89^ PWV in the pulmonary artery and its effects on RV hemodynamics can be measured non-invasively using the aforementioned variables^90–92^.

### Lung Mechanics Measurement and Analysis

Mice were anesthetized using urethane (1 g/kg administered in 10% solution with sterile water) and tracheostomized, then connected to the Flexivent system (FlexiVent®, SCIREQ©, Montreal, QC, Canada). Succinylcholine (1 mg/kg) was administered via intraperitoneal injection to eliminate spontaneous breathing. FlexiVent perturbations and oscillations were performed and analyzed using the FlexiWare Version 7.6 software, Service Pack 6 to obtain lung pressure-volume loops, static compliance (Cst), and inspiratory capacity (IC). Maneuvers and perturbations continued until acquiring three suitable measurements. A coefficient of determination of 0.95 was the lower limit for suitable measurements. An average of three measurements for each metric was calculated per mouse.

### Ultrasonography

Noninvasive investigation of cardiac function was performed in additional mice using transthoracic echocardiograms under light anesthesia (1.5% isoflurane) while maintaining physiological temperature^93^. Standardized cardiac views were obtained with a high-resolution ultrasound system (Vevo 2100, VisualSonics, Toronto, ON, Canada) equipped with an ultrahigh frequency (40 MHz) linear array transducer. B-mode two-dimensional (2D) images of the RV and right atrium (RA) were obtained from an apical four-chamber view, and the pulmonary artery was obtained from a parasternal short-axis view at the level of the aortic valve. In addition, M-mode and tissue Doppler imaging (TDI) of the lateral tricuspid annulus were obtained from the apical four-chamber view. The pulmonic valve was imaged at the level of the leaflet tips with pulsed wave Doppler. RV outflow tract (RVOT) diameter, tricuspid annular plane systolic excursion (TAPSE), RV systolic myocardial velocity (s’), pulmonary artery acceleration time (PAT), pulmonary ejection time (PET), two-dimensional end systolic (ES) and diastolic (ED) right ventricle area, and RA area were measured offline using Vevo Lab software (version 3.2.6, VisualSonics) by an experienced sonographer. Fractional area change of the right ventricle was computed as (ED-ES)/EDLJ×LJ100^94^.

### Multiphoton Imaging and Collagen Fiber Orientation Analysis

A titanium-sapphire laser (Chameleon Vision II, Coherent) was used to image representative regions of pulmonary arteries at *in vivo* relevant loading conditions (*in vivo* stretches and pressures identical to those used during passive mechanical testing, described below). A LaVision Biotec TriMScope microscope was tuned at 820 nm and equipped with a water immersion 20X objective lens (NA. 0.95). The backward scattering second harmonic generation signal from fibrillar collagens was detected within the wavelength range 390-425 nm; the auto-fluorescent signal arising from elastin was detected at 500-550 nm, and the fluorescent signal of cell nuclei labelled with Syto red stain was detected above 550 nm. An in-plane field of view (axial-circumferential plane) of 500 µm x 500 µm and a volume of about 0.05 mm^3^ were used; this provides a much greater volume of tissue for imaging than via standard histology and hence averaging over greater numbers of cells and ECM. The in-plane resolution was 0.48 µm/pixel and the out-of-plane (radial direction) step size was 1 µm/pixel. 3D images acquired concurrently for the three signals (collagen, elastin, and cell nuclei) were post-processed using MATLAB R2019b and ImageJ 1.53a. The first processing step relied on the near cylindrical shape of the samples to fit a circle to the two-dimensional mid-thickness profile of the arterial wall and transform each circumferential-radial slice of the 3D images from polar coordinates (angle and radius) to Cartesian. This allowed a layer-specific microstructural analysis to focus on collagen fiber alignment and cell volume density analyses, as described previously^9, 95^.

### Single Cell RNA Sequencing (scRNA-seq) and Analyses

We analyzed viable cells from the main pulmonary arteries of young and old mice. Following euthanasia, the heart, lungs, and pulmonary arteries were excised with surrounding tissue. The arteries were mechanically chopped and placed in 1mg/ml collagenase (Roche) and 3 U/ml elastase (Worthington). Following cellular dissociation, we barcoded unique mRNA molecules of each cell using our 10x Genomics Chromium platform (3’ v3.1 kit) and a droplet-based microfluidic system as previously described^96^, then performed reverse transcription, cDNA amplification, fragmentation, adaptor ligation, and sample index PCR according to the manufacturer’s protocol. High sensitivity DNA bioanalyzer traces of cDNA after barcoding and of the final cDNA library were evaluated for quality control. The final cDNA libraries were sequenced on a HiSeq 4000 Illumina platform in our core facility aiming for 150 million reads per library. Raw sequencing reads were demultiplexed based on sample index adaptors, which were added during the last step of cDNA library preparation. Possible adaptor and/or primer contamination were removed using *Cutadapt*. We processed the trimmed reads using the scRNA-seq implementation of STAR (STARsolo), where reads were mapped to the murine reference genome GRCm38 release M22 (GRCm38.p6), collapsed and counted, and summarized to a gene expression matrix. Data were analyzed and visualized using the Seurat R package^97^. Specifically, we clustered the cellular transcriptomes and visualized them in a uniform manifold approximation and projection (UMAP) space to delineate cell types (Supplemental Figure S3).

Differentially expressed genes (DEGs) between old and young mice were identified using a generalized linear mixed effects (GLME) model assuming that the read counts follow negative binomial distribution. Only genes expressed in >5% of cells in at least three samples were included for the DEG analysis. Genes that did not achieve model convergence were filtered out. Eventually, we selected genes with a nominal p-value < 0.05 and fold-chage>2 as significant DEGs. To understand the biological function of the significant DEGs, we conducted pathway enrichment analysis using Enrichr^98–100^ for four cell types (endothelial cells (ECs), SMCs, fibroblast (FBs), myeloid). A senescence score was calculated for each cell in the dataset using the extensively validated SenMayo gene list and the AddModuleScore function in Seurat R package^42^. Our implementation compared the average expression of each SenMayo gene to the expression of a set of 50 random control genes. SenMayo scores were compared using a Wilcoxon rank sum test. To identify cells exhibiting a senescence-like phenotype, we set a SenMayo threshold above which cells were designated “high SenMayo” cells.

To analyze the intercellular communications amongst SMCs, FBs, and macrophages (MΦs), connectomic analysis in single-cell RNAseq data was performed using the NICHES R pacakge, which computes cell-cell interactions by multiplying expression of ligand (in sender cell) with expression of receptor (in receiving cell)^101^. Single-cell RNAseq data was imputed with the ALRA algorithm using genes that were expressed in at least 50 cells. Ligand-receptor lists from FANTOM5 were used to filter for biologically relevant signaling mechanisms. Cell-to-cell signaling was calculated using NICHES for each sample individually, and later merged across samples. The resulting dataset was filtered to include only cell pairs with non-zero connectivity for at least 100 mechanisms. Cell signaling data was scaled and followed by PCA and UMAP calculation. Differential connectivity analysis was performed using a Wilcoxon Rank Sum test. Ligand-receptor pairs were analyzed for gene enrichment using the gProfiler R package.

### Statistics

For binary variables with only two categories, such as young or old, significance of differences between the two categories in morphological, mechanical, functional, and microstructural properties were assessed by a two-tailed unpaired Two-Sample Welch’s *t* test. For categorical variables with more than two categories (e.g., levels of stretch or pressure), significance of across-categories differences in outcomes were assessed using a two-factor analysis of variance (ANOVA) test. A global test across all levels was performed and then pair-wise comparisons were conducted with post-hoc tests using Bonferroni correction. A *p <* 0.05 level of significance was used, with data reported as mean ± standard error from the mean (SEM). All statistical tests were performed using GraphPad Prism version 7.01 for Windows, GraphPad Software, La Jolla California USA, www.graphpad.com. Statistical genetic analyses (outlined above) were performed in R.

## Data Availability

Single-cell RNAseq data generated for this study is available from NCBI Gene Expression Omnibus (GSE299310).

## Funding

NK - R01HL127349, R01HL141852, U01HL145567, UH3 TR002445, R21HL161723

NSAH - SNSF PM_210847

XY – NLM R01LM014087

EPM – NIA R03AG074063 GEMSSTAR, VA VISN1 CDA, Dr. Manning was supported by a Pilot Grant from the Claude D. Pepper Older Americans Independence Center at Yale School of Medicine (P30AG021342)

ABR – NSF DMS 2436623

## Contributions of Authors

RDM, ZC – performed single cell RNA sequencing analyses and prepared figures and manuscripts

PD – performed 2-photon imaging analyses

NG – performed echocardiography and analyses

LL – performed pulmonary function tests and analyses

AJ – performed single cell RNA sequencing

NSAH – performed single cell RNA sequencing

CC – performed 2-photon imaging and supervised 2-photon imaging analyses

MSR – designed and supervised NICHES intercellular communication analyses

PH – analyzed RV pressure tracings

IS – edited manuscript

XY – supervised single cell RNA sequencing analyses

MJK – edited manuscript, emphasis PFT aging

DB – edited manuscript, emphasis RV aging

PL – edited manuscript, emphasis senescence

GT – supervised experiments, edited manuscript, large vessel aging and immunobiology

JDH – supervised experiments, edited manuscript, biomechanics and mechanobiology

NK – supervised experiments, edited manuscript, transcriptomics and aging

ABR – performed biomechanical experiments, prepared figures, edited manuscript

EPM – participated in all experiments, supervised experiments, edited and prepared figures, edited manuscript

## Notes

https://www.ncbi.nlm.nih.gov/geo/query/acc.cgi?acc=GSE299310

### Competing Interest Statement

The authors have declared no competing interest.

