## Supplemental Information for "Proximal Pulmonary Artery Stiffening as a Biomarker of Cardiopulmonary Aging"

Ruben De Man*

Zhongyu Cai*

Pramath Doddaballapur

Nicole Guerrera

Liqin Lin

Aurelien Justet

Nebal S. Abu Hussein

Cristina Cavinato

Micha Sam B. Raredon

Paul Heerdt

Inderjit Singh

Xiting Yan

Min-Jong Kang

Danielle R. Bruns

Patty J. Lee

George Tellides

Jay D. Humphrey

Naftali Kaminski

Abhay B. Ramachandra+

Edward P. Manning+

*,+authors contributed equally

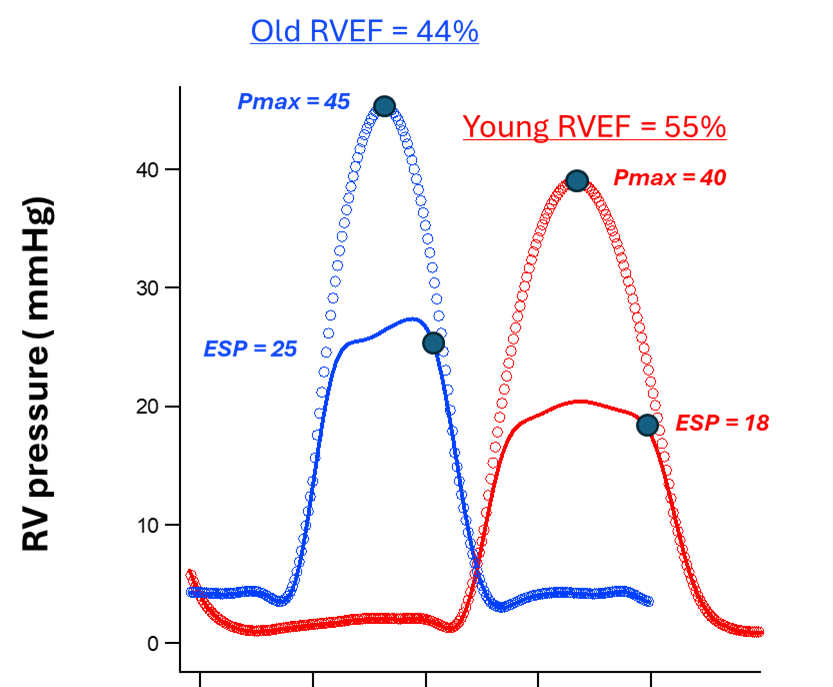

**Supplemental Figure S1:** Right ventricle (RV) mechanical characteristics. Standardized sampling and filters to create an example comparison of young (n=3) and old (n=3) RV pressure waveforms from which ejection fraction was calculated using our pressure-based method^1^, where EF is calculated as (Pmax-ESP)/Pmax, or 1-(ESP/Pmax). Peak and end-systolic pressures are elevated in older mice. Pmax = maximum pressure; ESP = end-systolic pressure.

[1] Elassal, A., et al. (2021). "Pressure-based estimation of right ventricular ejection fraction: Validation as a clinically relevant target for drug development in a rodent model of pulmonary hypertension." *Journal of Pharmacological and Toxicological Methods* 112: 107102.

**Supplemental Figure S2:** additional associations of changes in right ventricular and lung mechanics and changes in circumferential stiffness of the proximal pulmonary artery

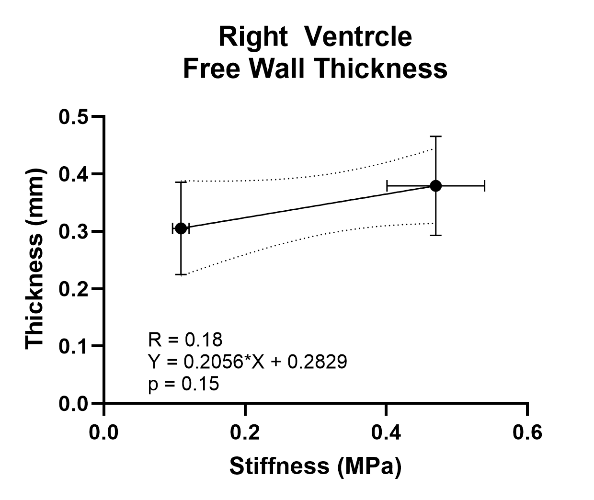

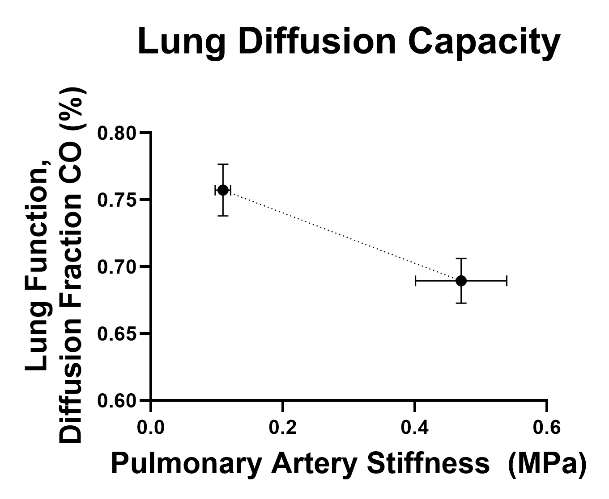

**Supplemental Figure S3:** Uniform manifold approximation and projection (UMAP) embedded as cell type, age grouping, and individual samples.

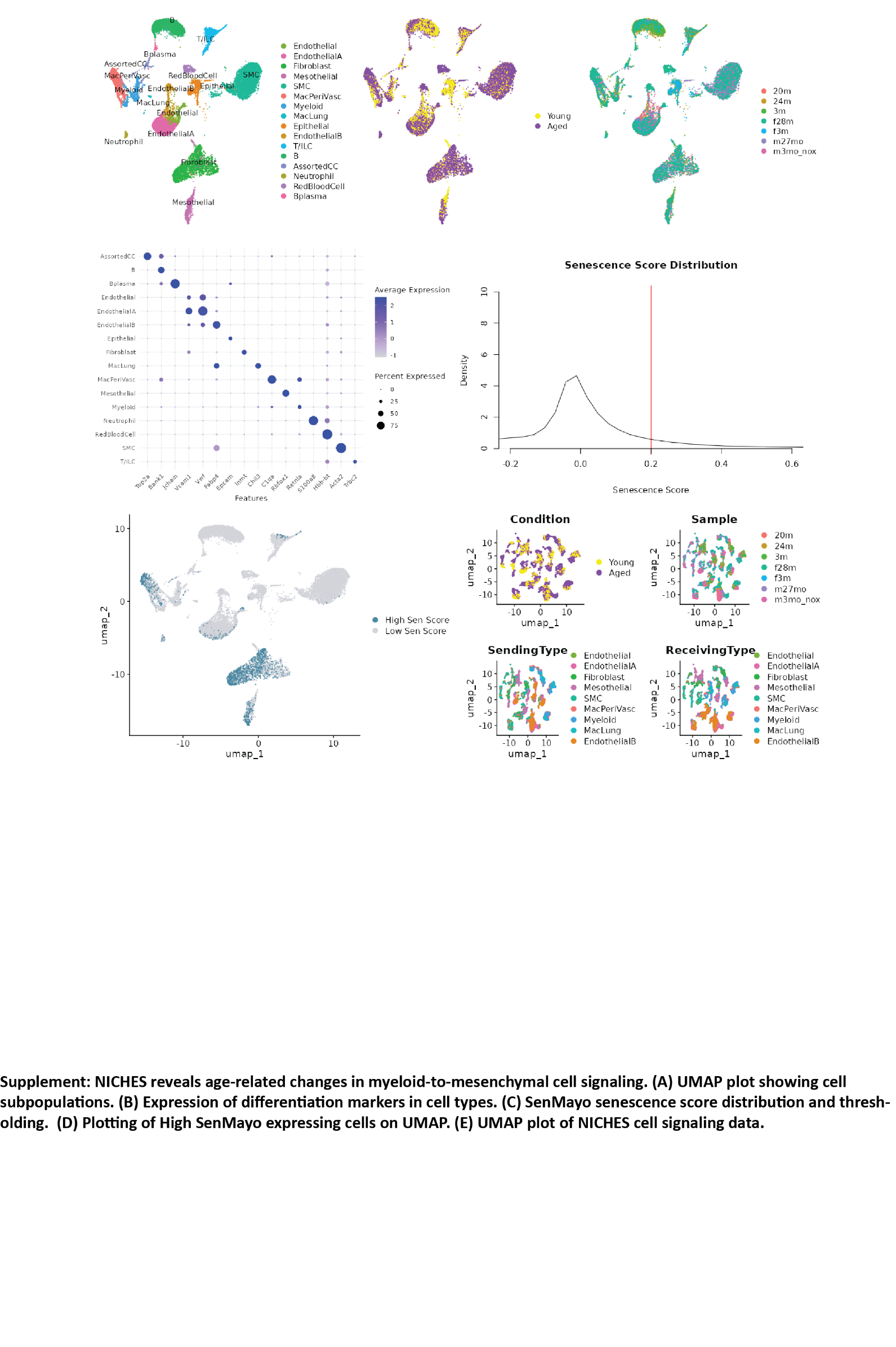

A heatmap demonstrates gene markers used to define cell types.

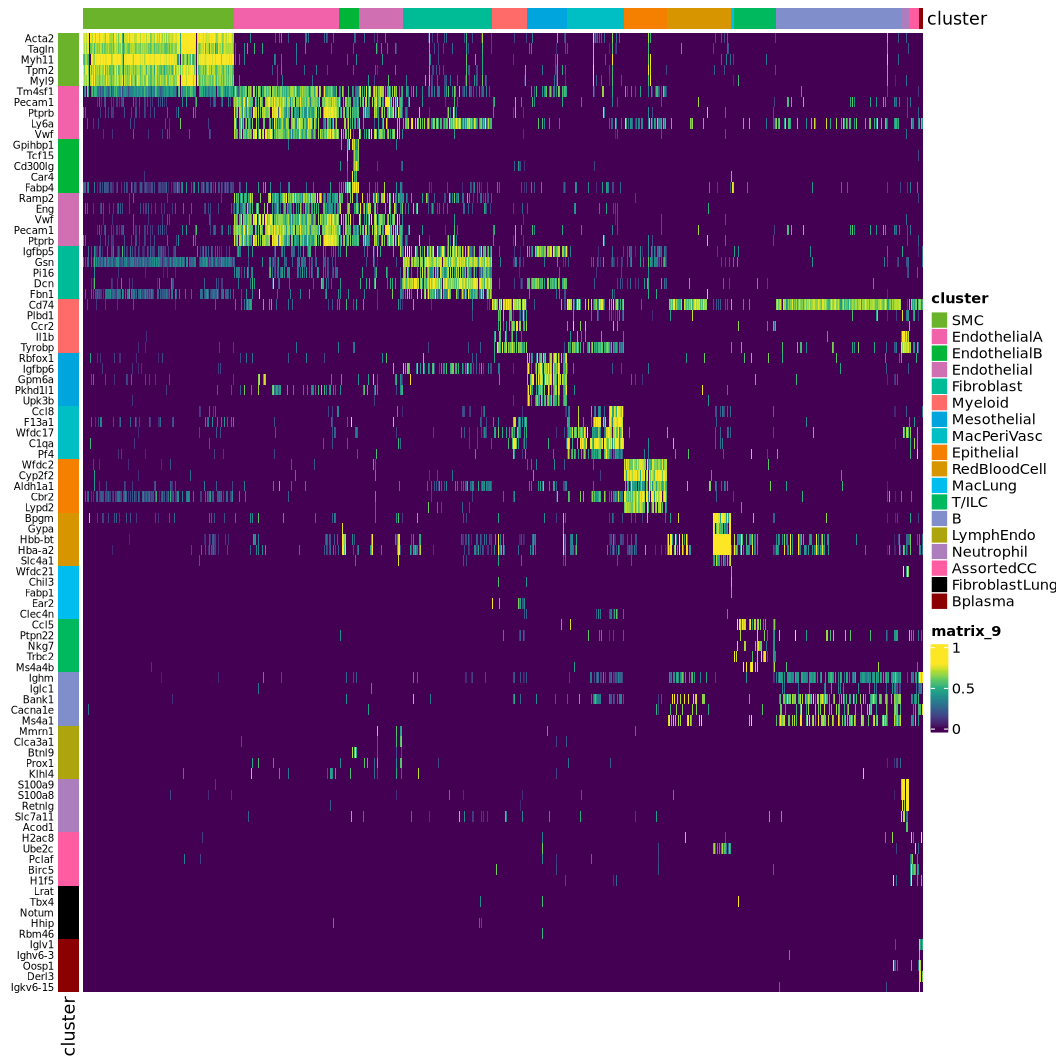

Left: Expression of differentiation markers in cell types. Right: Cell proportions are similar between young and old age groups.

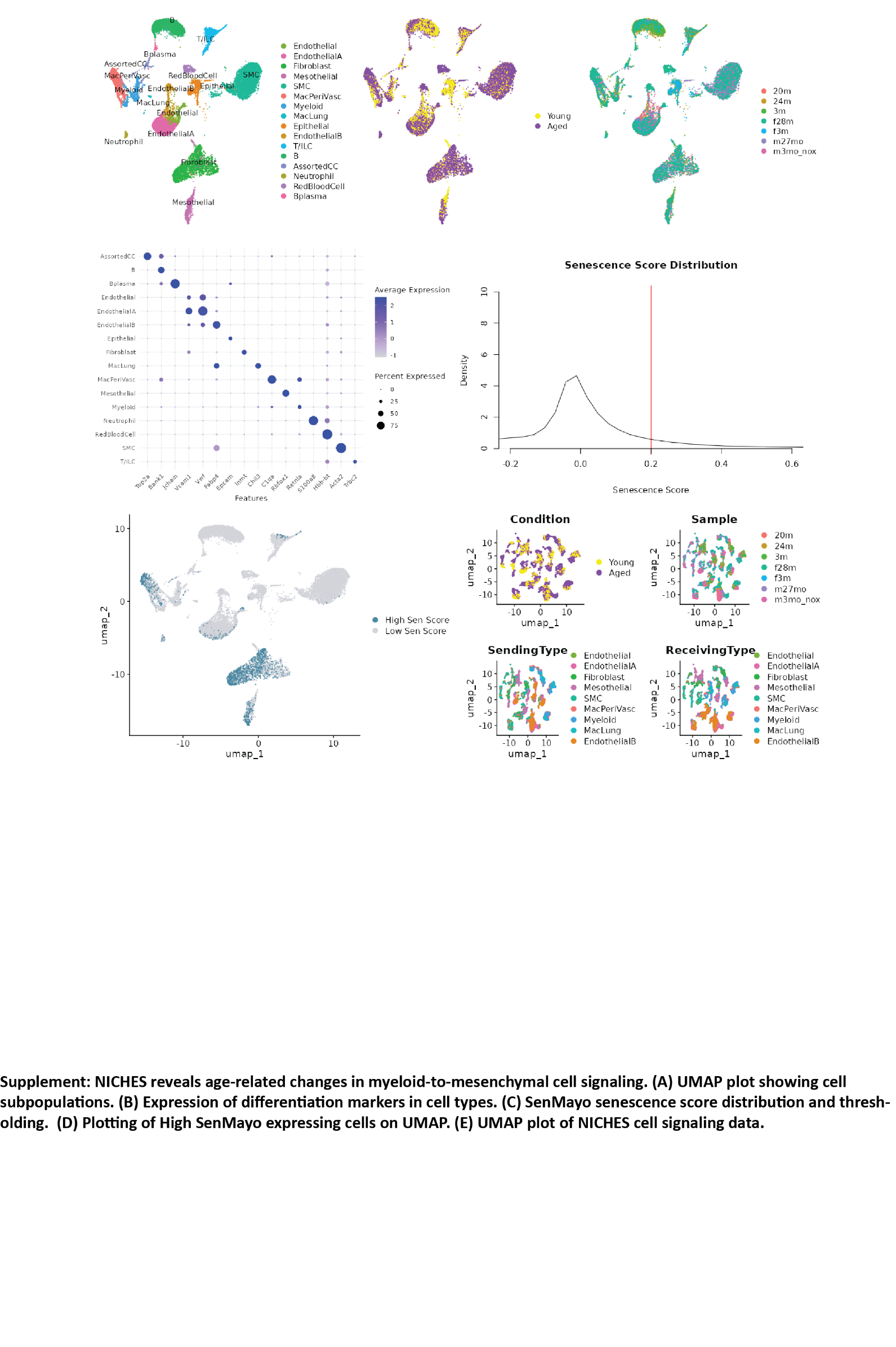

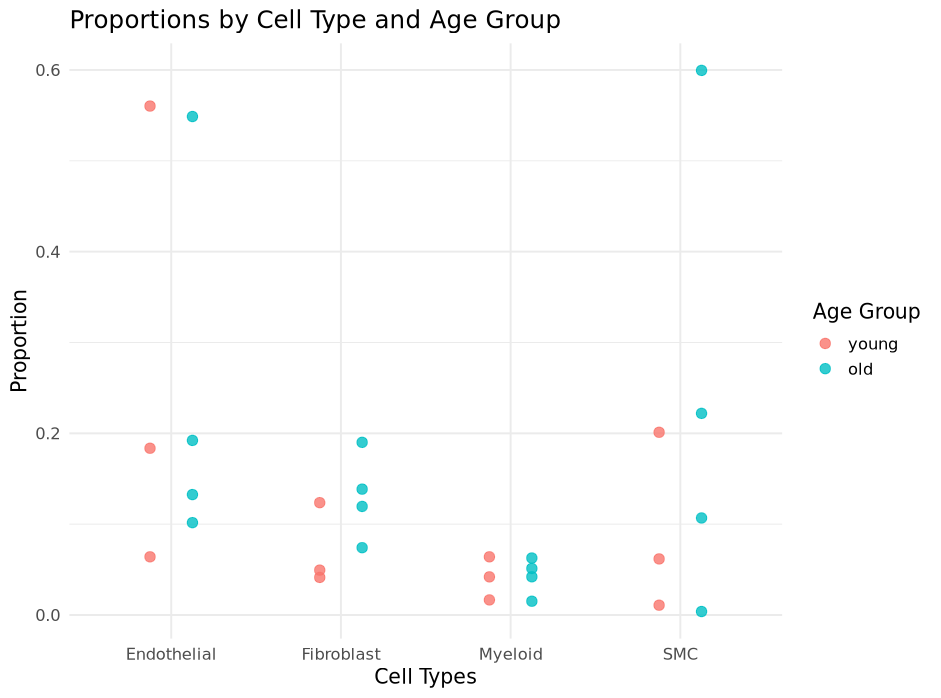

Differentiating general monocytic cells (Myeloid), macrophages that reside in vascular wall (MacPeriVasc), and macrophages that tend to reside in alveoli (MacLung).

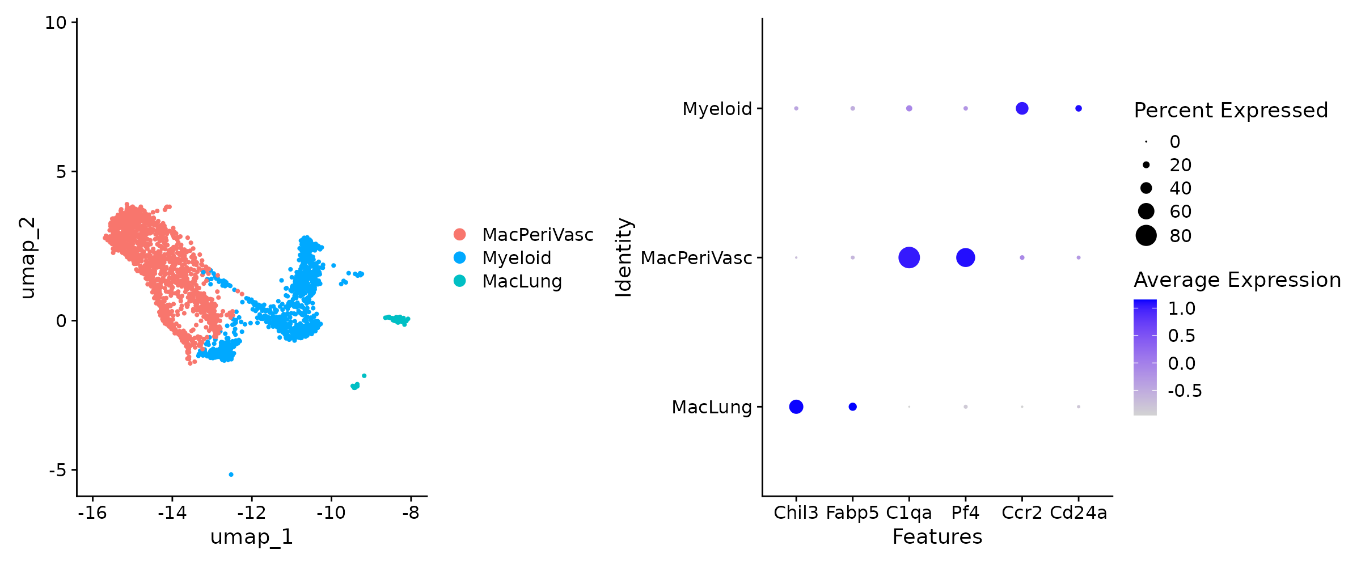

SenMayo senescence score distribution and thresholding.

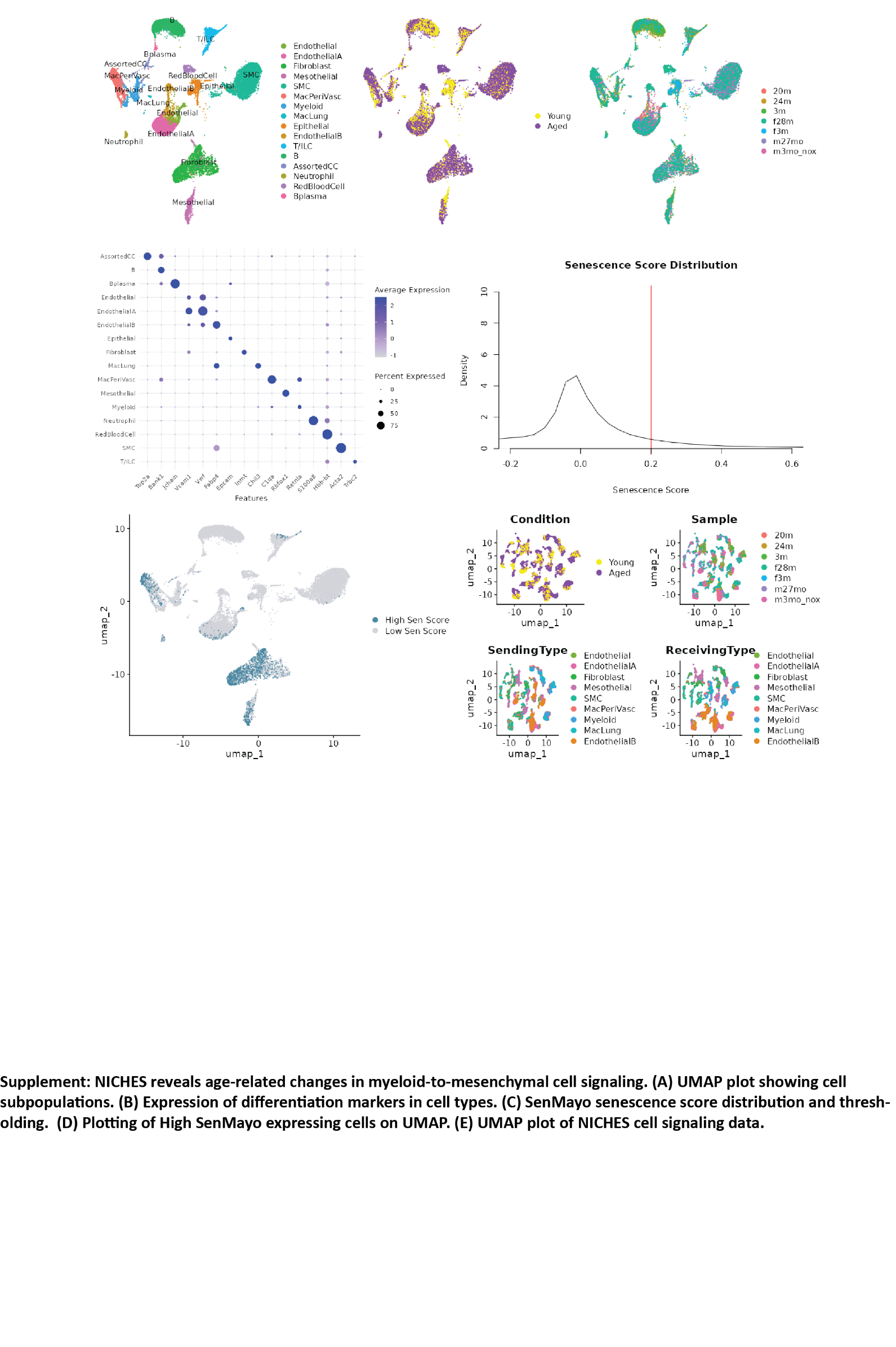

Left: Plotting of High SenMayo expressing cells on UMAP. Right: UMAP plot of NICHES cell signaling data.

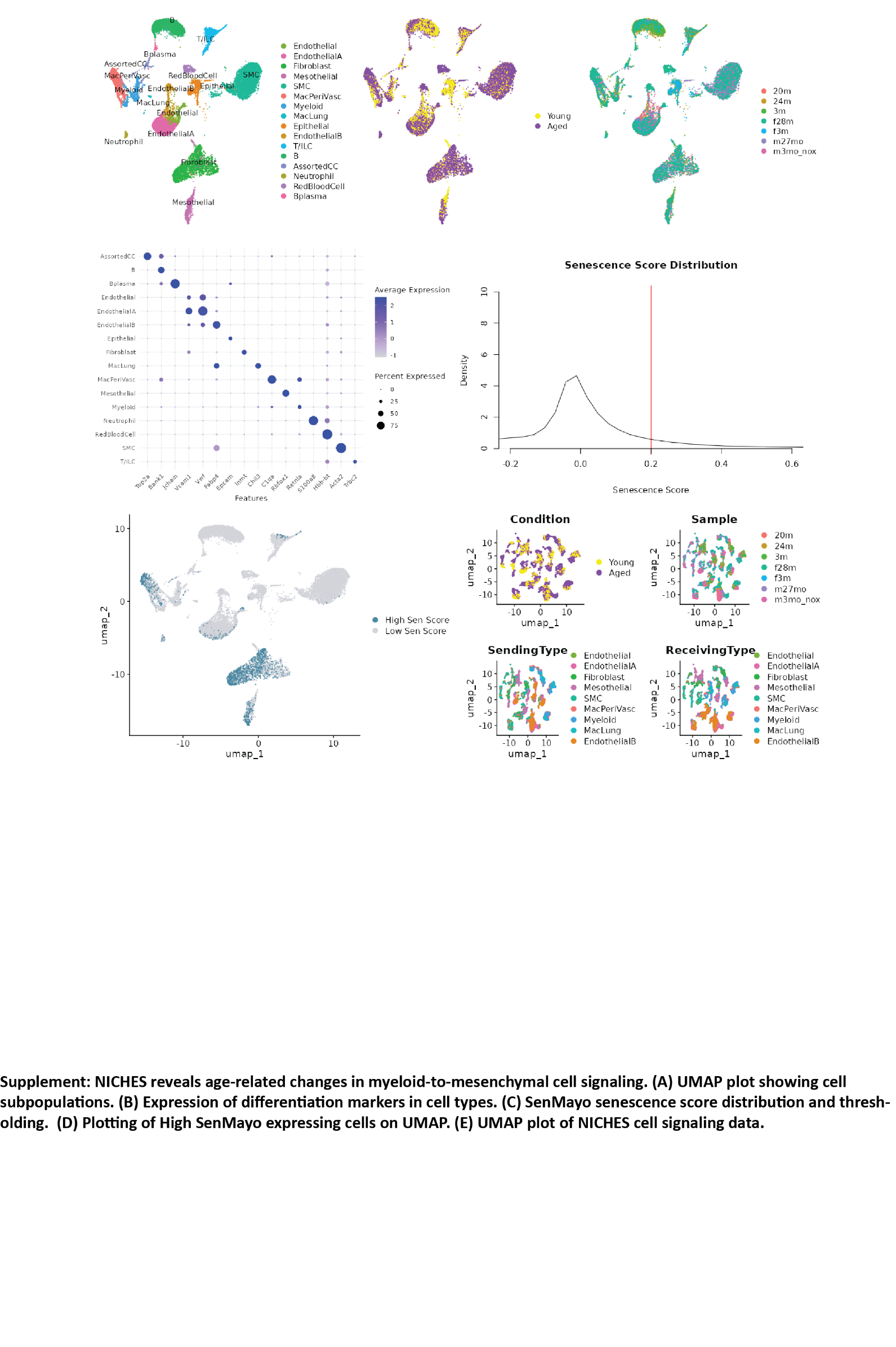

**Supplemental Figure S4**

Gene Enrichment Analysis – Biological Processes (Panther)

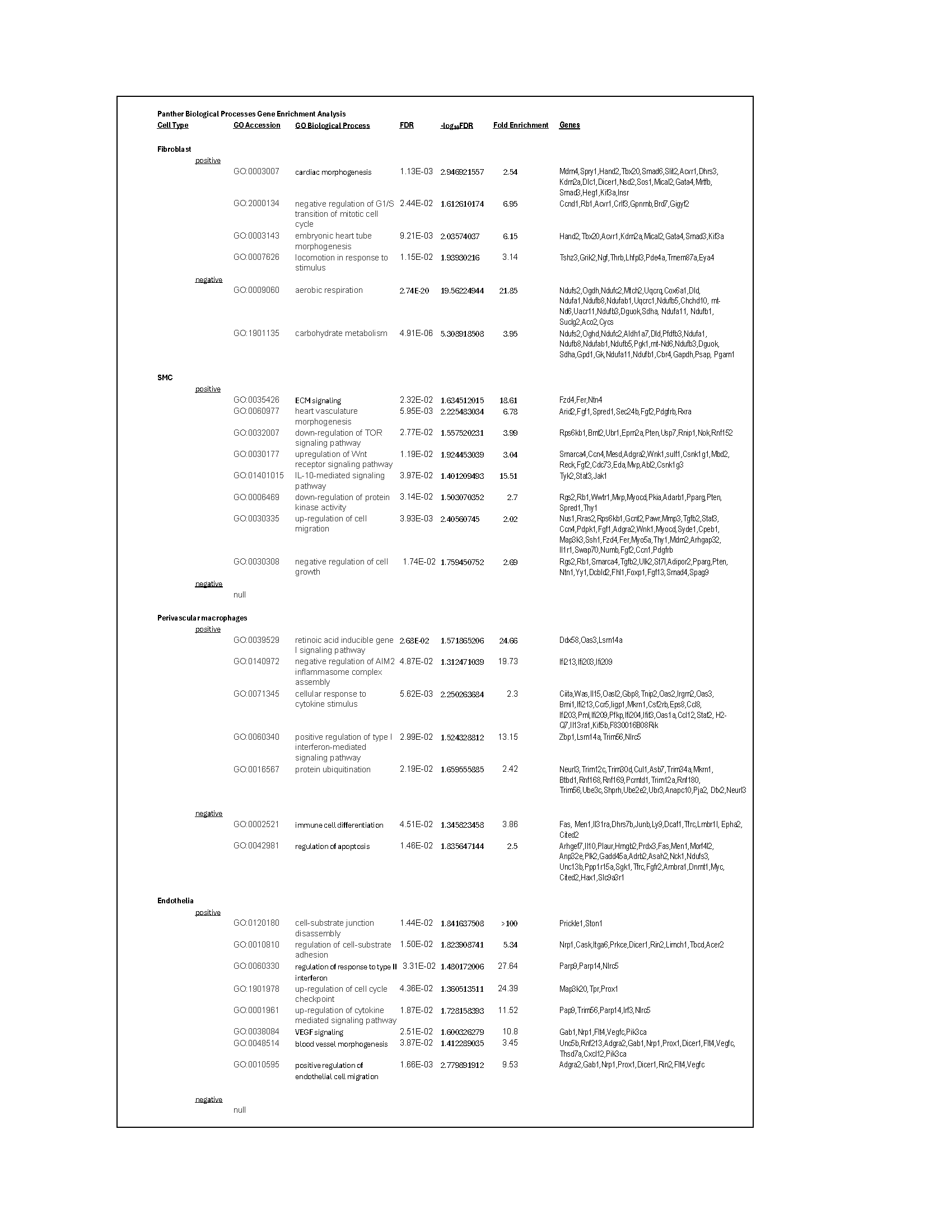

Gene Enrichment Analysis – Pathways (Panther & WikiPathways Mouse)

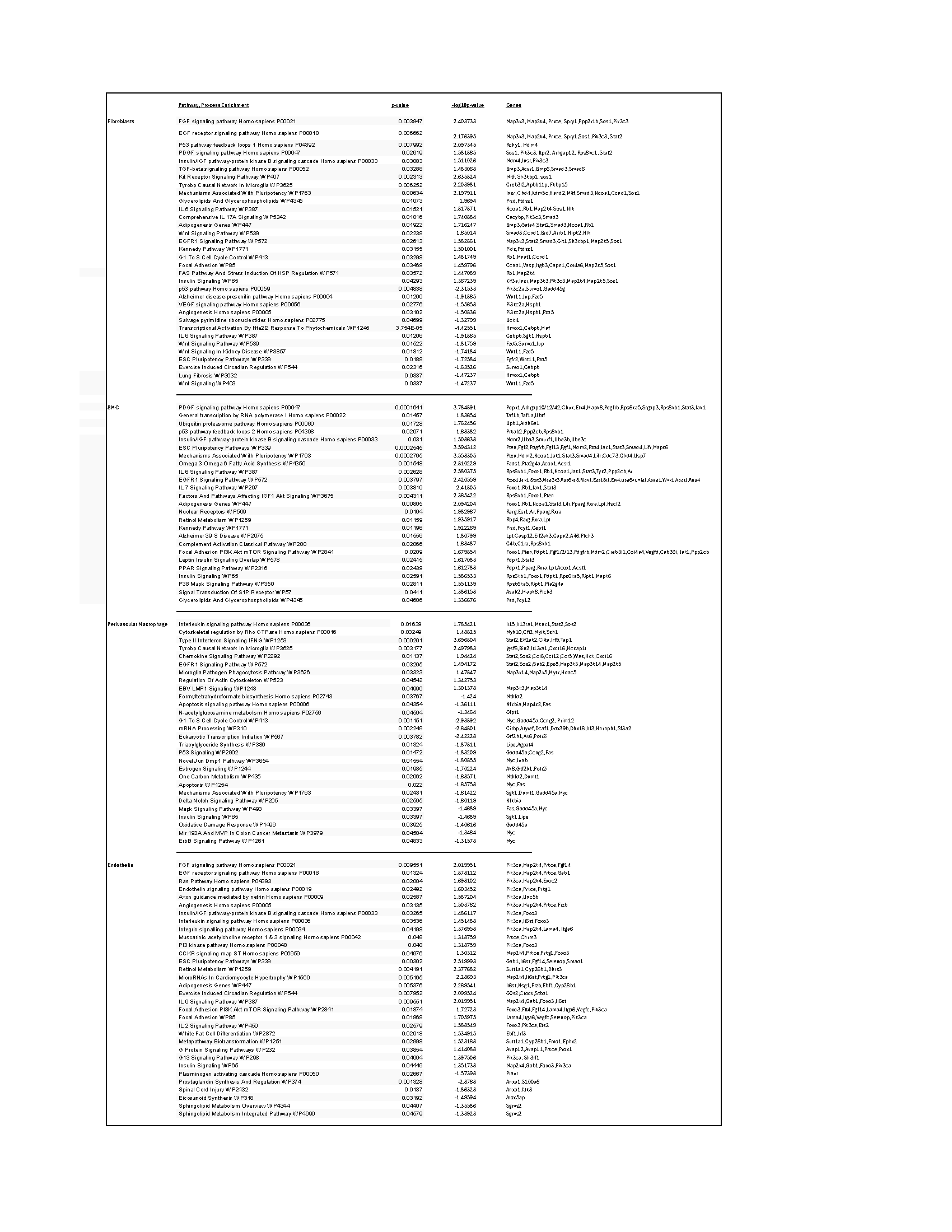

Fibroblast positive gene enrichment analysis using Enrichr:

Panther 2016:

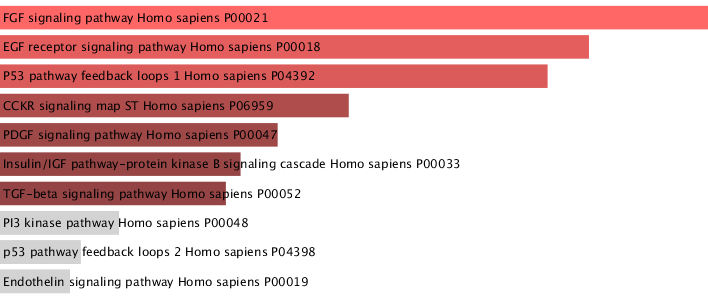

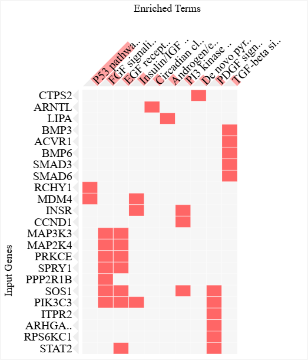

WikiPathways 2024 Mouse:

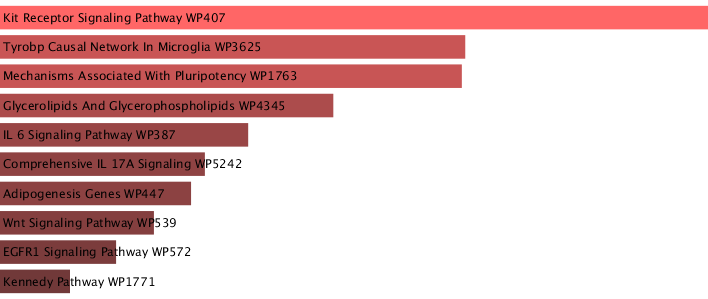

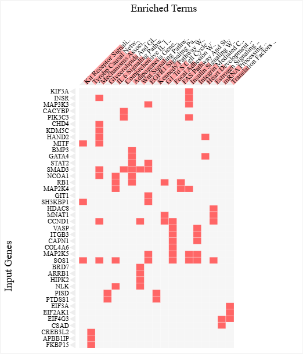

Fibroblast negative gene enrichment analysis using Enrichr:

Panther 2016:

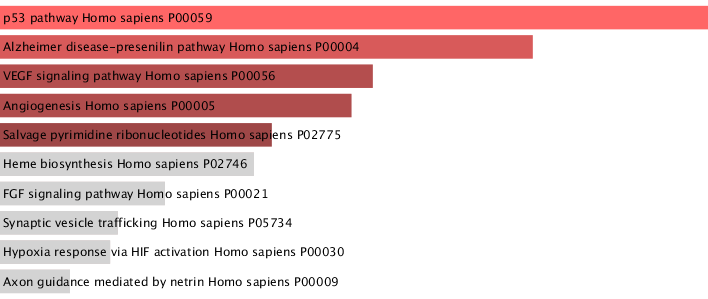

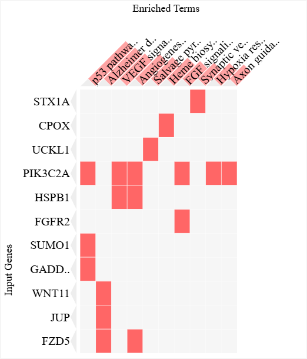

WikiPathways 2024 Mouse

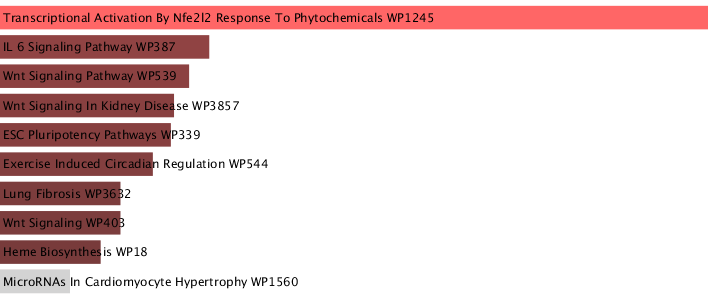

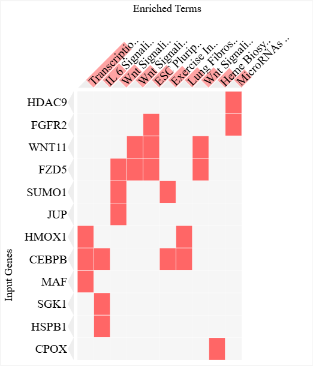

SMC positive gene enrichment analysis using Enrichr:

Panther 2016:

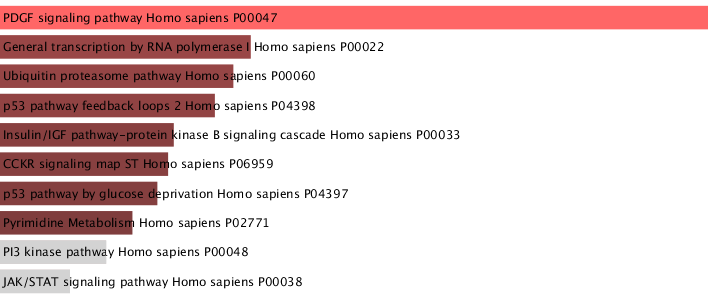

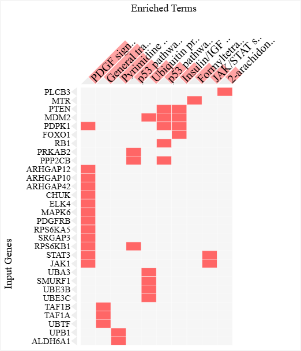

WikiPathways Mouse 2024

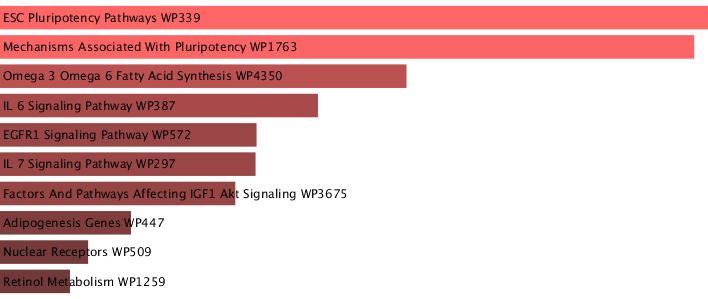

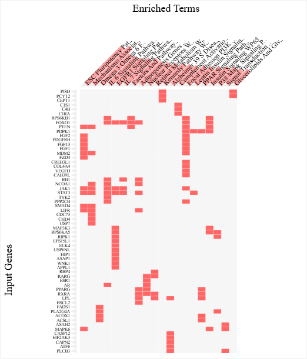

Perivascular Macrophage positive gene enrichment analysis using Enrichr:

Panther 2016:

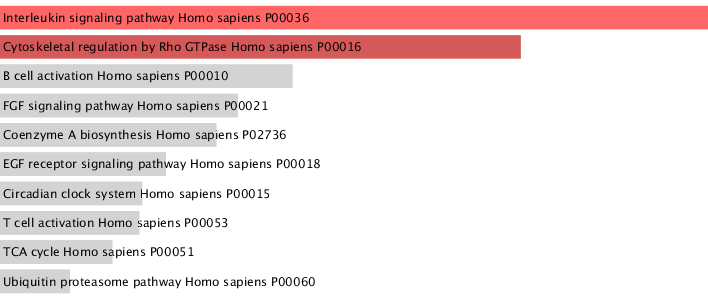

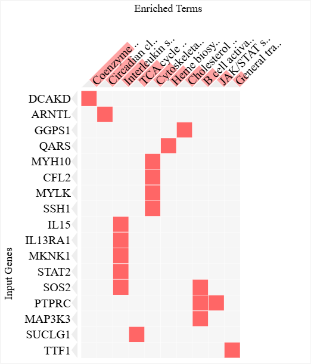

Mouse WikiPathway 2024:

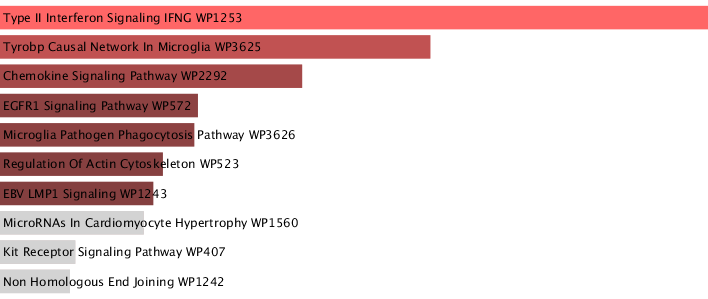

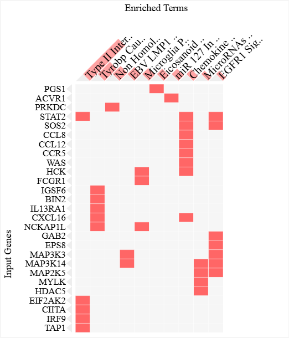

Perivascular Macrophage negative gene enrichment analysis using Enrichr:

Panther 2016:

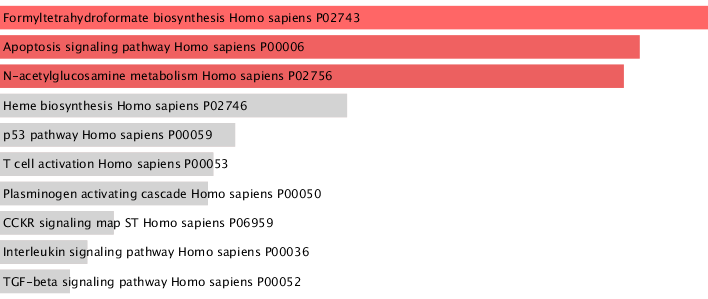

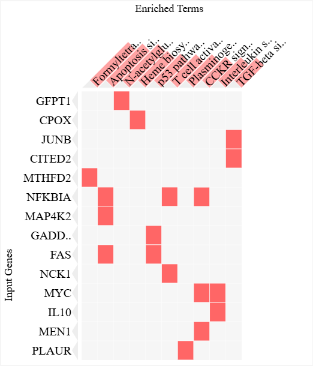

WikiPathways Mouse 2024:

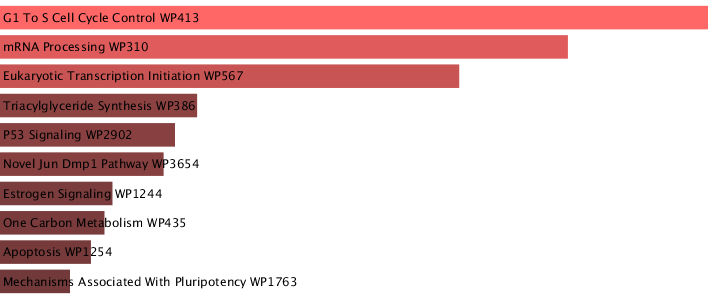

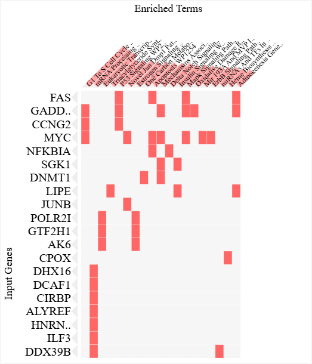

EC positive gene enrichment analysis using Enrichr:

Panther 2016:

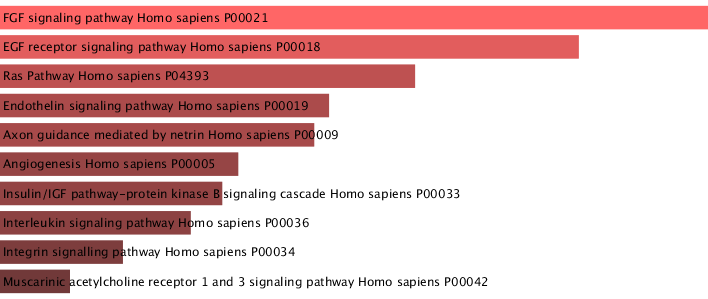

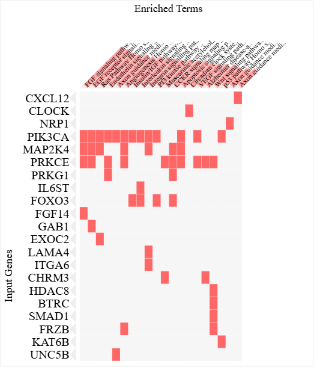

WikiPathways 2024 Mouse:

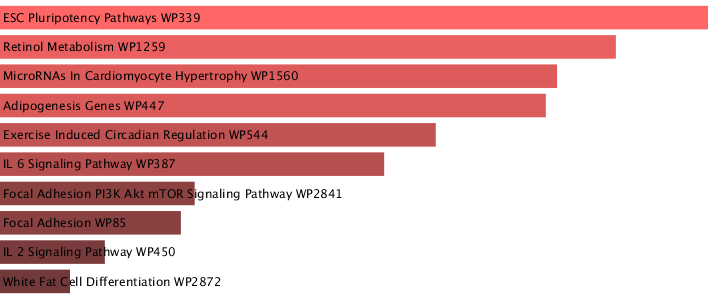

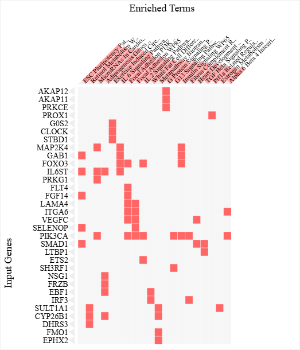

EC negative gene enrichment analysis using Enrichr:

Panther 2016:

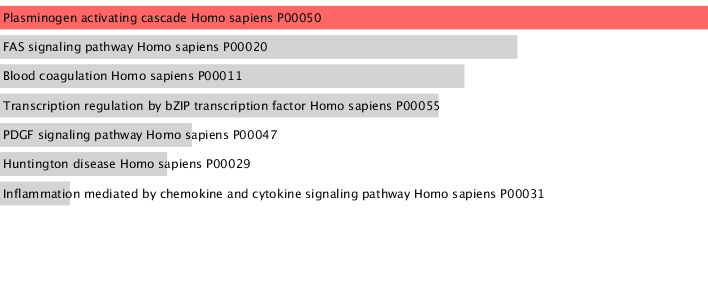

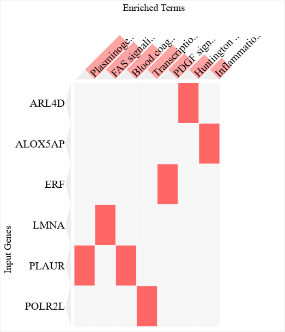

WikiPathways 2024 Mouse:

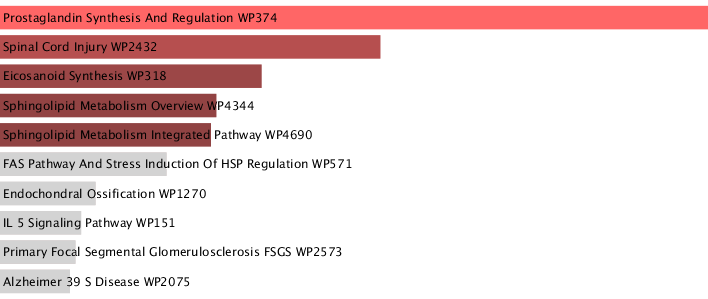

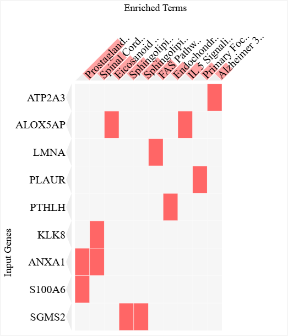

**Supplemental Figure S5**

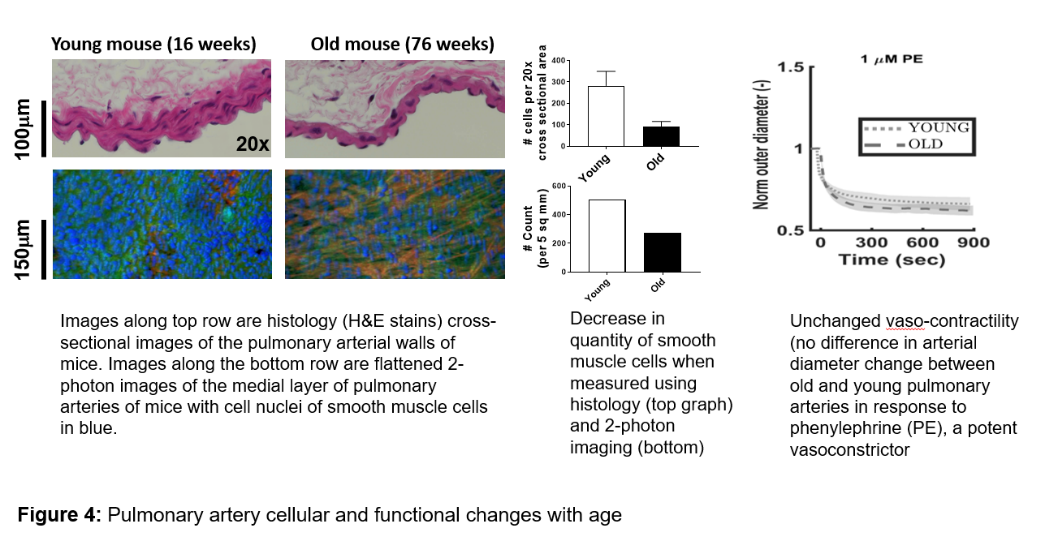

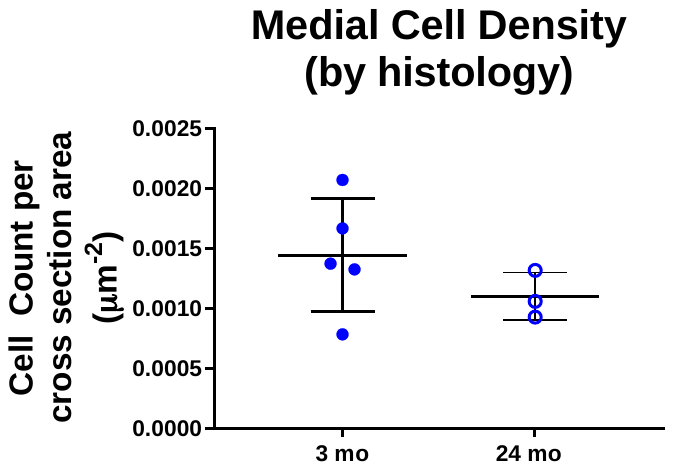

Left: Images on the top row are 5um thick, histology slices of formalin fixed, paraffin-embedded cross-sections of proximal pulmonary arteries from mice that are stained with H&E. Images on the bottom row are representative z-slices of the medial layer of flattened 2-photon images of proximal pulmonary arteries from mice in which the cell nucleir of SMC’s are stained with syto 17. Right: There is not a significant decrease in the density of smooth muscle cells of proximal PA’s from young and old mice when quantified by histology (H&E). This is similar to our findings when using 2-photon imaging (cf. Figure 2D).

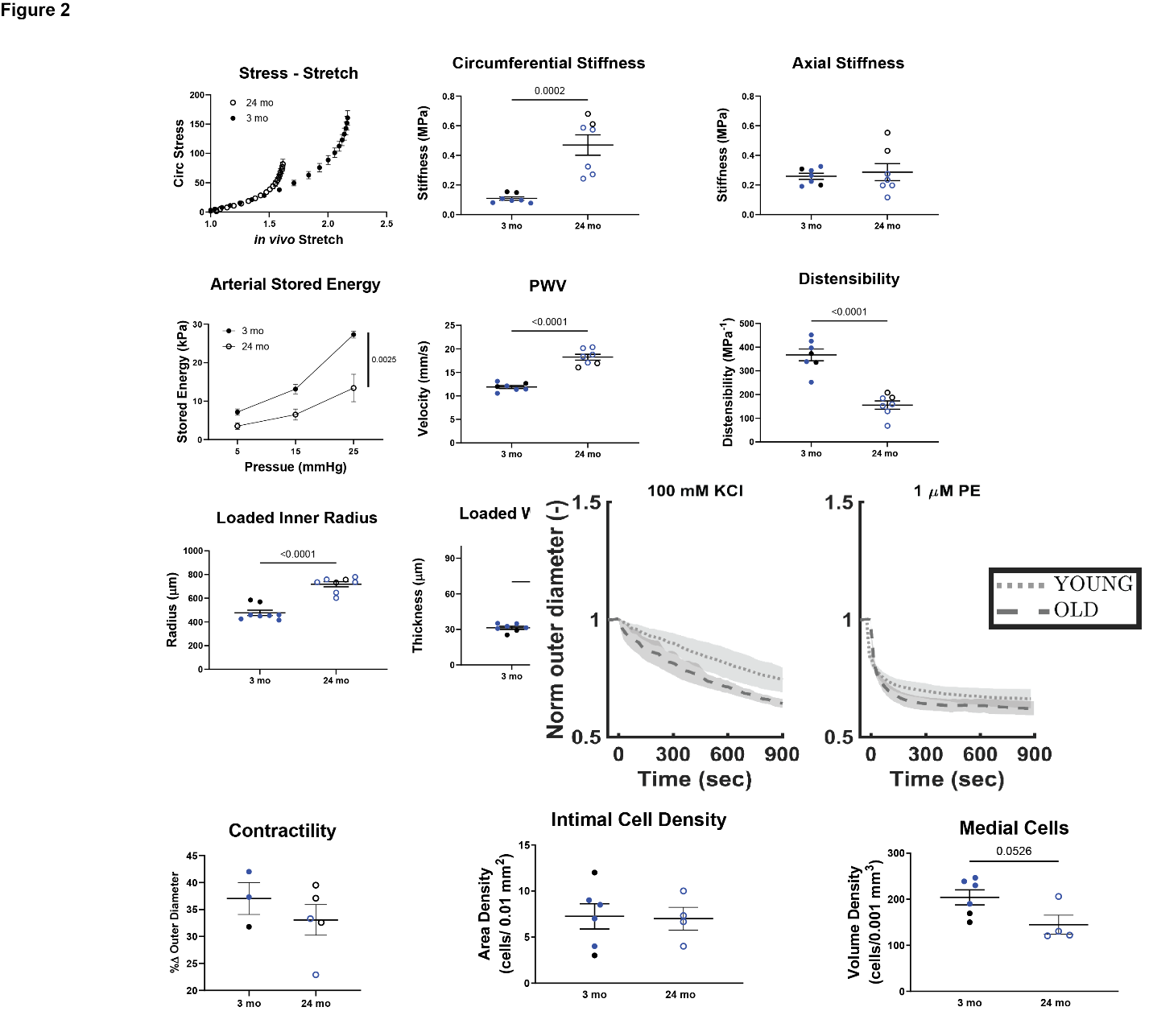

We found no significant difference in the vaso-contractility (based on change in of arterial diameter from baseline) in response to potassium chloride (KCl) or phenylephrine (PE).

**Supplemental Table S1:** additional cardiac RV data

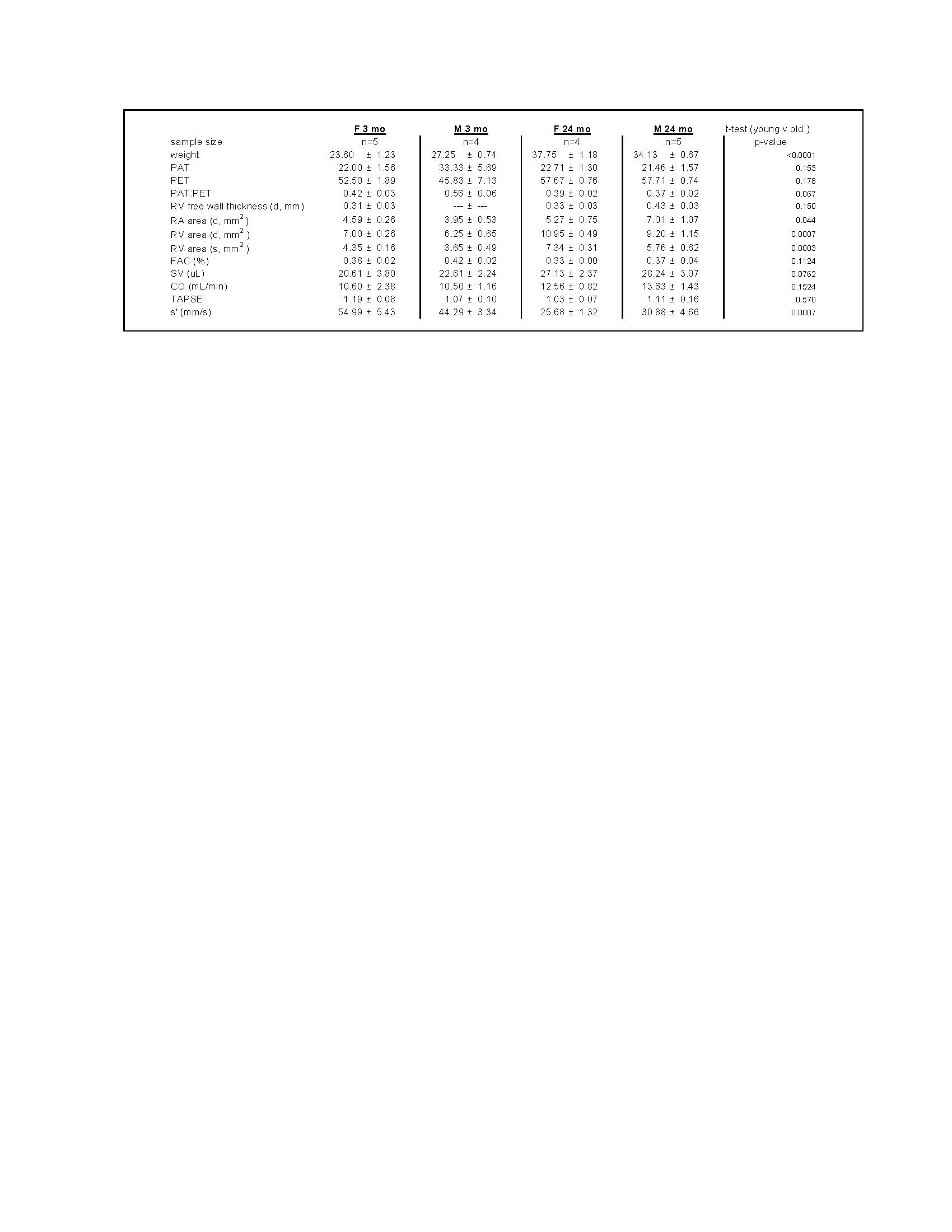

**Supplemental Table S2:** additional lung mechanics

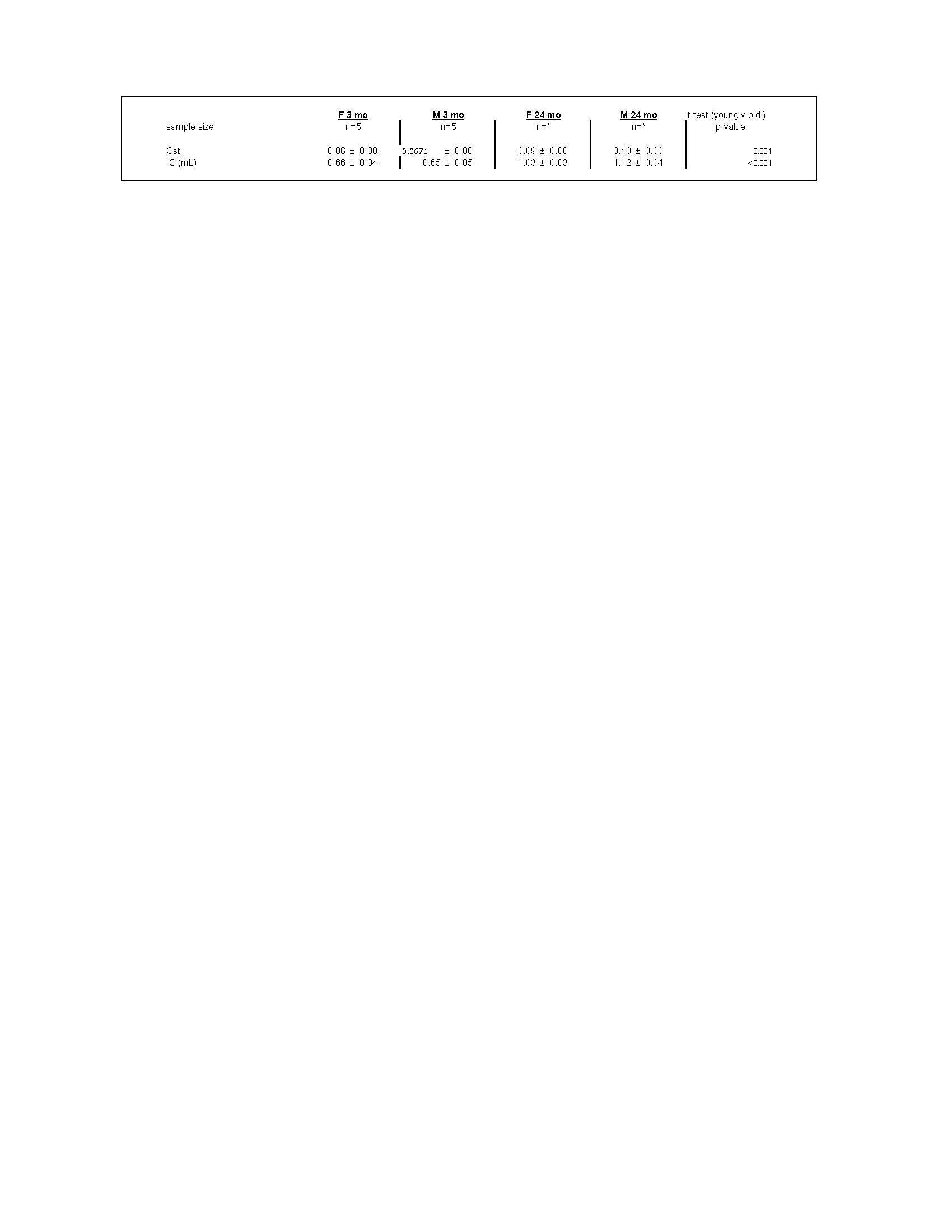

**Supplemental Table S4:** biomechanical table (common and MAP) and material properties

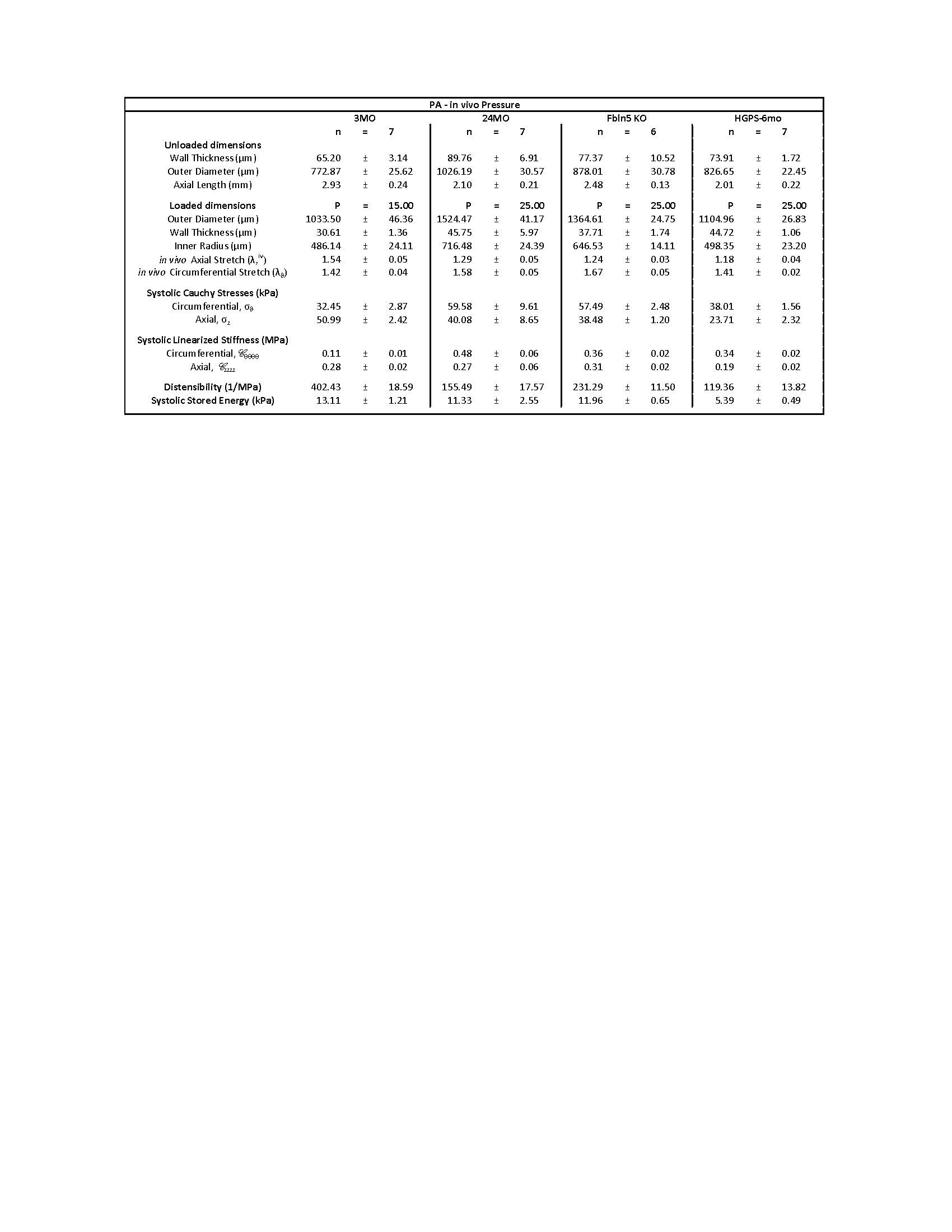

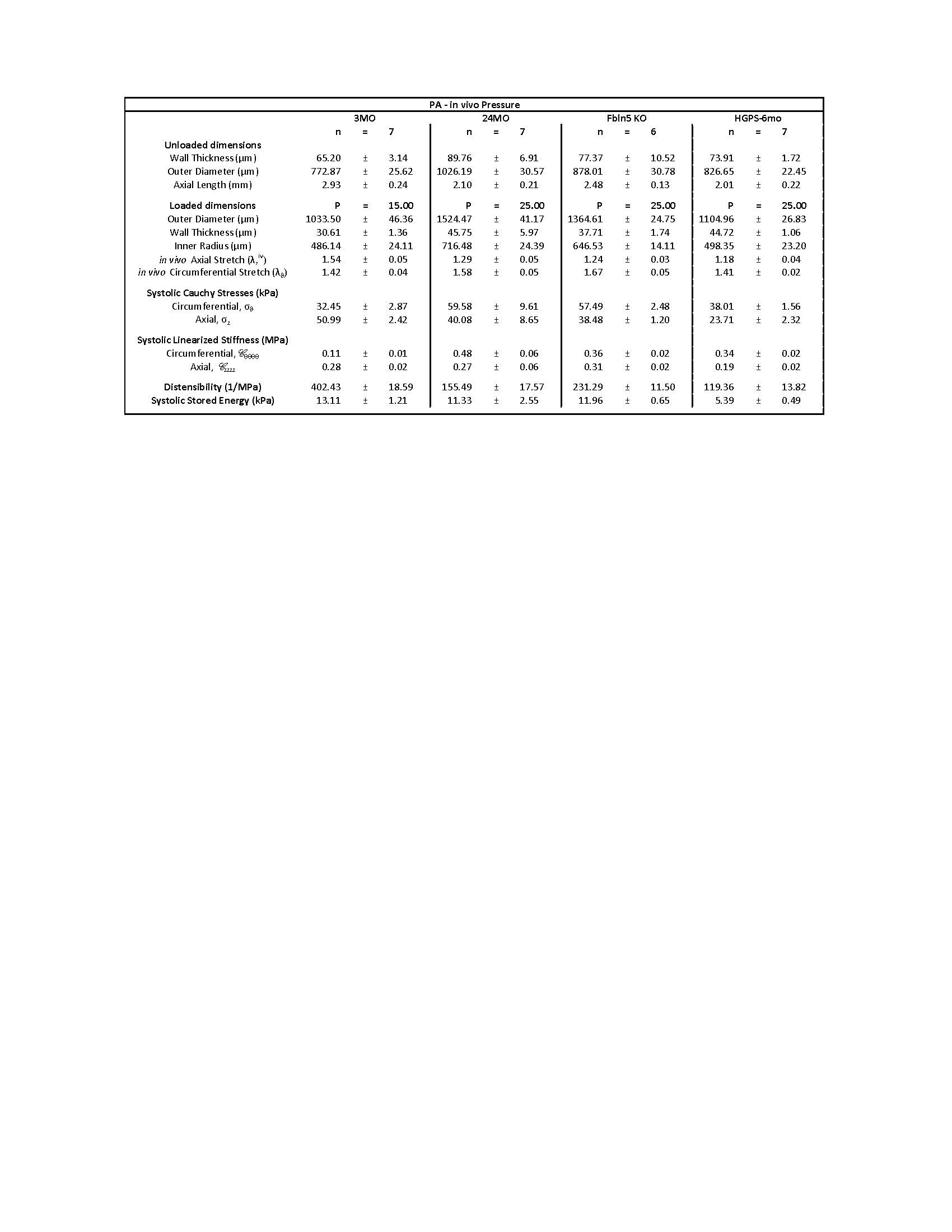

**Supplemental Table S5:** Top 50 genes differentially expressed genes from GLME model by cell type. Gene name is the first column. Β_0_ = intercept coefficient. Β_1_ = slope coefficient. Disp = dispersion parameter, Sigma = standard deviation, P.Beta = p-value, AIC = Akaike information criterion, fdrP = FDR-corrected p-value.

**Fibroblast: gene expression, positive correlation (β_1_) with age**

| **gene** | **Beta0** | **Beta1** | **Disp** | **Sigma** | **P.Beta** | **AIC** | **fdrP** |
| --- | --- | --- | --- | --- | --- | --- | --- |
| Itih5 | -9.2953 | 0.731 | 0.7305 | 7.02E-05 | 5.97E-08 | 4587.9208 | 3.52E-05 |
| Vps13d | -10.4628 | 1.1157 | 8.1092 | 2.70E-05 | 8.71E-08 | 2858.1634 | 4.97E-05 |
| C130026I21Rik | -10.1638 | 1.2109 | 1.2616 | 1.22E-01 | 1.21E-07 | 3724.8345 | 5.87E-05 |
| Prelp | -9.1327 | 0.7997 | 1.6873 | 1.00E-01 | 1.50E-07 | 5210.2008 | 7.07E-05 |
| Pknox2 | -11.4461 | 1.705 | 1.2443 | 6.29E-05 | 5.65E-07 | 2279.0031 | 2.22E-04 |
| Kdm2a | -9.7268 | 0.7347 | 4.9688 | 1.94E-02 | 1.22E-06 | 3565.9903 | 4.28E-04 |
| Mllt10 | -9.5271 | 0.7147 | 4.5736 | 6.15E-02 | 3.08E-06 | 3951.8103 | 8.48E-04 |
| Mdm4 | -10.8094 | 1.1451 | 4.4466 | 3.32E-05 | 3.89E-06 | 2400.4257 | 9.98E-04 |
| Trim8 | -10.384 | 0.93 | 4.7436 | 4.20E-05 | 4.07E-06 | 2736.8369 | 9.98E-04 |
| Ngf | -10.5408 | 1.0882 | 0.4094 | 6.06E-05 | 4.15E-06 | 2739.5772 | 9.98E-04 |
| Lipa | -10.5112 | 0.9936 | 2.5004 | 5.12E-05 | 4.16E-06 | 2659.9914 | 9.98E-04 |
| Klhl29 | -10.0326 | 0.8381 | 0.7139 | 3.16E-05 | 4.36E-06 | 3236.8469 | 1.03E-03 |
| Rab7b | -11.0042 | 1.2496 | 3.138 | 2.85E-05 | 4.78E-06 | 2159.1945 | 1.11E-03 |
| Gypc | -11.8501 | 1.8889 | 10.4464 | 2.00E-02 | 5.37E-06 | 1928.2471 | 1.23E-03 |
| Ilrun | -10.55 | 0.9862 | 663.4363 | 3.54E-14 | 5.53E-06 | 2526.967 | 1.25E-03 |
| Map2k5 | -11.095 | 1.3961 | 3.7014 | 1.09E-01 | 5.77E-06 | 2406.5062 | 1.29E-03 |
| Uba6 | -10.8086 | 1.1133 | 4.7893 | 3.33E-05 | 7.14E-06 | 2307.3531 | 1.55E-03 |
| Ncoa1 | -9.3136 | 0.6958 | 2.7014 | 8.82E-02 | 9.06E-06 | 4525.6458 | 1.85E-03 |
| Zfp950 | -10.7511 | 1.1781 | 2.259 | 6.62E-02 | 9.24E-06 | 2577.1546 | 1.86E-03 |
| Zscan26 | -11.3383 | 1.4025 | 14.191 | 1.55E-03 | 1.20E-05 | 2014.4047 | 2.26E-03 |
| Bmp6 | -10.765 | 1.1766 | 0.2241 | 5.05E-05 | 1.36E-05 | 2385.6943 | 2.51E-03 |
| Kdm5b | -10.2417 | 0.813 | 3.4207 | 4.09E-05 | 1.77E-05 | 2793.8488 | 3.05E-03 |
| Pou2f1 | -10.6466 | 0.98 | 4.1355 | 9.01E-05 | 1.98E-05 | 2408.7022 | 3.35E-03 |
| Matn2 | -10.738 | 1.1242 | 0.2826 | 3.11E-05 | 2.22E-05 | 2428.9485 | 3.63E-03 |
| Insr | -10.5026 | 0.9189 | 2.0363 | 3.57E-05 | 2.27E-05 | 2602.4958 | 3.68E-03 |
| Csnk1g3 | -9.9774 | 0.7 | 8.7619 | 1.37E-02 | 2.75E-05 | 3011.2309 | 4.14E-03 |
| G3bp2 | -10.0296 | 0.7099 | 6.7698 | 2.40E-05 | 2.95E-05 | 2979.3657 | 4.35E-03 |
| Tlk2 | -10.2761 | 0.804 | 3.2147 | 2.13E-05 | 3.04E-05 | 2715.7473 | 4.46E-03 |
| Dennd4c | -10.1093 | 0.8373 | 3.1187 | 3.98E-02 | 3.54E-05 | 3083.1892 | 4.96E-03 |
| Creb3l2 | -9.7201 | 0.7272 | 3.7087 | 8.25E-02 | 3.96E-05 | 3633.6731 | 5.28E-03 |
| Tshz3 | -10.1449 | 0.739 | 3.1904 | 3.80E-05 | 4.62E-05 | 2841.7856 | 5.88E-03 |
| Nlk | -10.4242 | 0.8548 | 1.4126 | 3.78E-05 | 4.91E-05 | 2592.7975 | 6.10E-03 |
| Trabd2b | -11.8958 | 1.9617 | 0.2965 | 2.28E-01 | 5.36E-05 | 2109.9989 | 6.52E-03 |
| Pdlim4 | -10.1842 | 0.75 | 2.0264 | 3.44E-05 | 5.46E-05 | 2720.3569 | 6.59E-03 |
| Fgd5 | -11.4683 | 1.6298 | 0.9704 | 2.05E-01 | 5.89E-05 | 2302.1267 | 7.05E-03 |
| Myoc | -11.2976 | 1.8002 | 0.0312 | 7.01E-05 | 6.05E-05 | 1485.7066 | 7.20E-03 |
| Hectd4 | -10.9339 | 1.055 | 8.4818 | 3.15E-05 | 6.19E-05 | 2028.3943 | 7.31E-03 |
| Gm10125 | -11.4442 | 1.3808 | 0.6395 | 3.90E-05 | 6.58E-05 | 1864.4436 | 7.72E-03 |
| Zmym2 | -10.1154 | 0.7057 | 3.7801 | 2.99E-05 | 7.64E-05 | 2860.2696 | 8.84E-03 |
| Dop1b | -12.032 | 1.7859 | 3.7786 | 3.29E-05 | 7.77E-05 | 1604.5663 | 8.92E-03 |
| Acvr1 | -10.2774 | 0.7606 | 3.5165 | 3.59E-05 | 7.93E-05 | 2680.5406 | 8.93E-03 |
| Dnajc13 | -10.4639 | 0.8598 | 6.1451 | 2.53E-02 | 8.01E-05 | 2512.5002 | 8.95E-03 |
| Chm | -10.8098 | 0.98 | 4.3158 | 5.48E-05 | 8.14E-05 | 2180.2088 | 9.03E-03 |
| Sparcl1 | -10.1182 | 1.0653 | 0.1639 | 1.50E-01 | 8.20E-05 | 3066.6461 | 9.04E-03 |
| Sacm1l | -10.3088 | 0.757 | 7.9843 | 9.70E-05 | 9.92E-05 | 2568.4062 | 1.05E-02 |
| Arrb1 | -10.8684 | 0.9954 | 4.6147 | 3.00E-05 | 1.02E-04 | 2188.6169 | 1.05E-02 |
| Slc10a6 | -10.3223 | 0.8464 | 0.4016 | 4.42E-05 | 1.04E-04 | 2702.0686 | 1.05E-02 |
| Ap3d1 | -10.8079 | 0.9586 | 129.3556 | 1.94E-14 | 1.05E-04 | NA | 1.05E-02 |
| Dicer1 | -10.643 | 0.8895 | 4.3507 | 3.30E-05 | 1.10E-04 | 2294.5008 | 1.09E-02 |
| Rabl6 | -10.6496 | 0.8906 | 3.6543 | 3.02E-05 | 1.10E-04 | 2297.6298 | 1.09E-02 |

**Fibroblast: gene expression, negative correlation (β_1_) with age**

| **gene** | **Beta0** | **Beta1** | **Disp** | **Sigma** | **P.Beta** | **AIC** | **fdrP** |
| --- | --- | --- | --- | --- | --- | --- | --- |
| Ttll12 | -8.6382 | -2.6566 | 0.0508 | 8.62E-05 | 4.16E-23 | 1049.425 | 6.87E-19 |
| Itm2a | -9.3468 | -1.4885 | 0.2626 | 4.56E-05 | 9.30E-16 | 1478.323 | 7.69E-12 |
| Dmac2l | -9.8757 | -1.6895 | 0.3593 | 6.85E-05 | 1.73E-15 | 908.7069 | 7.96E-12 |
| Cpox | -9.4389 | -1.2957 | 0.5889 | 5.05E-05 | 1.92E-15 | 1527.788 | 7.96E-12 |
| Slco2a1 | -9.803 | -2.1622 | 0.0708 | 8.67E-04 | 2.39E-13 | 711.7543 | 7.90E-10 |
| H4f16 | -10.692 | -2.7873 | 0.1687 | 6.66E-05 | 3.91E-12 | 307.1085 | 9.24E-09 |
| Nudt1 | -9.6817 | -2.2645 | 0.043 | 4.53E-05 | 6.75E-12 | 657.9736 | 1.40E-08 |
| Fnbp1l | -9.6907 | -1.6372 | 0.0915 | 5.73E-05 | 6.18E-11 | 964.9275 | 1.14E-07 |
| 1700003G18Rik | -10.274 | -3.1133 | 0.0205 | 1.01E-04 | 4.43E-10 | 290.3885 | 6.66E-07 |
| Bphl | -9.6899 | -0.9612 | 78.086 | 6.41E-08 | 6.25E-10 | 1520.678 | 8.61E-07 |
| Tnfrsf12a | -9.0814 | -0.7681 | 1.0615 | 7.69E-05 | 9.07E-10 | 2501.312 | 1.15E-06 |
| Osgep | -9.1346 | -0.8463 | 1.0297 | 4.66E-02 | 1.28E-09 | 2336.160 | 1.51E-06 |
| Cdh18 | -10.493 | -2.0258 | 0.1079 | 6.55E-05 | 2.09E-09 | 484.2409 | 2.31E-06 |
| A2ml1 | -11.174 | -2.8484 | 0.4034 | 4.56E-05 | 4.78E-09 | 204.1599 | 4.70E-06 |
| Grid2 | -10.422 | -1.3774 | 3524199.86 | 4.31E-12 | 4.83E-09 | NA | 4.70E-06 |
| Dpagt1 | -9.6713 | -1.1054 | 0.2521 | 4.55E-05 | 5.15E-09 | 1378.401 | 4.70E-06 |
| Gm45510 | -11.243 | -2.6514 | 4545327.03 | 5.44E-05 | 1.16E-08 | 213.7092 | 9.42E-06 |
| Otx2os1 | -10.851 | -3.5019 | 0.0322 | 7.17E-05 | 1.20E-08 | 190.442 | 9.42E-06 |
| Ptprn | -10.470 | -4.4676 | 1.3723 | 3.40E-01 | 1.48E-08 | 264.7124 | 1.11E-05 |
| Pik3c2a | -8.9811 | -0.6972 | 1.3703 | 2.54E-02 | 1.69E-08 | 2764.118 | 1.19E-05 |
| Ercc4 | -10.015 | -1.0887 | 2.8428 | 3.24E-02 | 1.73E-08 | 1112.549 | 1.19E-05 |
| Slc9a3r1 | -10.168 | -1.5825 | 0.3318 | 7.78E-02 | 1.93E-08 | 771.8676 | 1.28E-05 |
| 4930512B01Rik | -10.497 | -2.8067 | 0.0196 | 6.20E-05 | 4.77E-08 | 280.4491 | 3.04E-05 |
| Cldn10 | -10.447 | -1.3724 | 0.5746 | 3.37E-05 | 5.86E-08 | 729.5842 | 3.52E-05 |
| Larp7 | -9.5924 | -0.8176 | 1.8224 | 2.60E-05 | 9.39E-08 | 1790.066 | 5.01E-05 |
| Dstyk | -9.1945 | -0.7301 | 0.7726 | 7.15E-05 | 9.40E-08 | 2368.988 | 5.01E-05 |
| Gap43 | -10.451 | -2.0713 | 0.0797 | 1.83E-01 | 1.06E-07 | 474.3637 | 5.37E-05 |
| Nwd2 | -11.443 | -2.7384 | 1311581.877 | 2.09E-46 | 2.04E-07 | 169.6892 | 9.37E-05 |
| Gps2 | -9.4373 | -0.6985 | 11.2547 | 3.52E-05 | 2.23E-07 | 2097.352 | 9.96E-05 |
| Rpl37-ps1 | -11.443 | -3.4315 | 2786497.26 | 9.92E-05 | 2.64E-07 | 132.4164 | 1.15E-04 |
| Tmprss9 | -11.356 | -2.5308 | 0.2295 | 4.03E-05 | 2.74E-07 | 200.4893 | 1.16E-04 |
| Tmem108 | -10.414 | -2.8558 | 0.0133 | 1.72E-04 | 3.22E-07 | 269.5217 | 1.33E-04 |
| Ank3 | -9.5384 | -1.8611 | 0.0252 | 1.22E-04 | 4.84E-07 | 744.6009 | 1.95E-04 |
| Tulp2 | -11.075 | -1.9082 | 0.1366 | 6.05E-05 | 1.10E-06 | 323.7713 | 4.05E-04 |
| Dcps | -9.986 | -1.0185 | 0.3151 | 5.73E-04 | 1.36E-06 | 1193.286 | 4.69E-04 |
| Cgref1 | -11.134 | -1.9863 | 0.1283 | 4.50E-05 | 1.52E-06 | 307.8059 | 5.13E-04 |
| Gm26555 | -11.175 | -2.1371 | 0.0824 | 1.34E-04 | 1.60E-06 | 268.7858 | 5.31E-04 |
| Endog | -10.144 | -0.9316 | 3476670.23 | 6.96E-26 | 1.64E-06 | 1144.791 | 5.31E-04 |
| Gm15567 | -11.382 | -3.1842 | 0.0365 | 7.47E-05 | 1.81E-06 | 148.91 | 5.75E-04 |
| Dcc | -11.561 | -2.4665 | 1758633.03 | 3.83E-15 | 1.88E-06 | 185.0384 | 5.87E-04 |
| Gm14798 | -10.934 | -1.5757 | 0.306 | 3.21E-02 | 2.48E-06 | 435.8414 | 7.59E-04 |
| Ccdc171 | -9.3632 | -0.7073 | 0.5999 | 9.61E-05 | 2.90E-06 | 2205.983 | 8.48E-04 |
| Ezh2 | -9.793 | -0.9919 | 0.4369 | 9.12E-02 | 2.99E-06 | 1404.069 | 8.48E-04 |
| Nedd9 | -8.8019 | -1.049 | 0.2528 | 1.43E-01 | 3.00E-06 | 2529.770 | 8.48E-04 |
| Dhrs7b | -9.8435 | -0.8112 | 1.2263 | 3.02E-05 | 3.06E-06 | 1490.822 | 8.48E-04 |
| 1700112J16Rik | -11.561 | -2.3329 | 6895532.53 | 3.25E-18 | 3.07E-06 | 185.0453 | 8.48E-04 |
| Ak7 | -11.443 | -2.0452 | 2300573.49 | 1.73E-15 | 3.51E-06 | 235.0312 | 9.39E-04 |
| Sumf2 | -10.056 | -0.8676 | 4.6224 | 5.86E-05 | 3.55E-06 | 1254.0842 | 9.39E-04 |
| Acd | -10.307 | -1.0095 | 1.7319 | 4.17E-05 | 3.58E-06 | 982.9097 | 9.39E-04 |
| Fzd5 | -10.297 | -1.1468 | 1.0125 | 1.03E-01 | 4.04E-06 | 908.0828 | 9.98E-04 |

**Smooth Muscle Cell: gene expression, positive correlation (β_1_) with age**

| **gene** | **Beta0** | **Beta1** | **Disp** | **Sigma** | **P.Beta** | **AIC** | **fdrP** |
| --- | --- | --- | --- | --- | --- | --- | --- |
| Hsph1 | -10.0942 | 1.1491 | 4.5698 | 1.76E-05 | 6.57E-18 | 9213.3985 | 3.86E-15 |
| Mmp3 | -11.4854 | 2.4229 | 0.1109 | 1.14E-02 | 4.91E-16 | 6658.2179 | 2.04E-13 |
| Pawr | -9.2407 | 0.7762 | 1.899 | 2.74E-02 | 4.20E-15 | 11976.2017 | 1.52E-12 |
| Ccn2 | -7.1986 | 0.9221 | 2.0758 | 1.15E-01 | 6.38E-14 | 26112.9533 | 1.84E-11 |
| Rgs17 | -11.3877 | 1.9747 | 0.255 | 4.90E-02 | 6.95E-13 | 6816.0351 | 1.74E-10 |
| Kifap3 | -10.7697 | 1.2681 | 4.9598 | 1.07E-04 | 7.61E-12 | 6930.7296 | 1.58E-09 |
| Dnah7a | -11.4622 | 1.745 | 1.5384 | 1.72E-02 | 3.96E-11 | 6140.9425 | 6.96E-09 |
| Pcnx | -10.1104 | 0.8664 | 5.8275 | 2.40E-05 | 1.11E-10 | 7952.8409 | 1.81E-08 |
| Agfg1 | -10.5882 | 1.0804 | 9.0228 | 2.17E-02 | 2.78E-10 | 6875.1303 | 3.97E-08 |
| Rpa1 | -10.7716 | 1.1717 | 2.1219 | 2.93E-05 | 3.82E-10 | 6691.0415 | 5.30E-08 |
| Arhgap32 | -11.0753 | 1.3583 | 1.3214 | 2.22E-05 | 5.40E-10 | 6262.4913 | 7.24E-08 |
| Skap2 | -9.9663 | 0.7704 | 4.3267 | 9.37E-05 | 9.35E-10 | 8249.8697 | 1.19E-07 |
| Wdr33 | -10.0601 | 0.7994 | 6.5829 | 7.19E-05 | 1.03E-09 | 7947.7933 | 1.30E-07 |
| Tmem164 | -10.4833 | 0.9962 | 1.83 | 2.58E-05 | 1.17E-09 | 7160.9317 | 1.46E-07 |
| Acsl1 | -9.9953 | 0.8749 | 3.6813 | 3.52E-02 | 1.30E-09 | 8646.1828 | 1.58E-07 |
| Taok3 | -11.397 | 1.5326 | 3.6343 | 2.96E-05 | 1.33E-09 | 5636.9627 | 1.59E-07 |
| Pdgfrb | -9.528 | 0.6938 | 17.0688 | 4.86E-02 | 1.66E-09 | 9616.1365 | 1.89E-07 |
| Me1 | -10.5331 | 0.9895 | 7.0019 | 2.51E-05 | 1.95E-09 | 6800.1894 | 2.18E-07 |
| Med1 | -11.2807 | 1.4206 | 10.1408 | 2.58E-05 | 2.28E-09 | 5614.313 | 2.51E-07 |
| C1s1 | -10.963 | 1.4497 | 0.8784 | 1.11E-01 | 2.96E-09 | 7209.8645 | 3.16E-07 |
| Tlk2 | -10.451 | 0.958 | 4.7638 | 2.99E-02 | 4.57E-09 | 7045.1912 | 4.61E-07 |
| Wscd2 | -10.4549 | 0.9353 | 3.8488 | 2.83E-05 | 4.78E-09 | 6975.1837 | 4.79E-07 |
| Numb | -10.8752 | 1.1453 | 3.8529 | 4.27E-05 | 4.82E-09 | 6157.379 | 4.81E-07 |
| Rad21 | -10.3644 | 0.8741 | 4843269.996 | 1.32E-19 | 6.62E-09 | NA | 6.48E-07 |
| Sdc2 | -8.993 | 0.8364 | 3.8016 | 9.48E-02 | 7.90E-09 | 13537.4223 | 7.65E-07 |
| St3gal3 | -11.6819 | 1.6696 | 1.5939 | 2.46E-05 | 1.17E-08 | 5211.8994 | 1.11E-06 |
| Nsmce2 | -10.1842 | 0.8072 | 1.8127 | 2.97E-05 | 1.44E-08 | 7641.883 | 1.34E-06 |
| Fam91a1 | -11.1267 | 1.2412 | 29.0392 | 2.54E-04 | 1.70E-08 | 5553.7461 | 1.54E-06 |
| Topors | -10.951 | 1.1656 | 4.274 | 2.15E-02 | 2.13E-08 | 5991.8156 | 1.90E-06 |
| Prpf19 | -10.6447 | 0.9584 | 2960181.737 | 8.73E-41 | 3.15E-08 | NA | 2.68E-06 |
| Zcchc14 | -9.8908 | 0.707 | 8.9349 | 3.19E-02 | 4.50E-08 | 8249.36 | 3.67E-06 |
| Invs | -11.175 | 1.2516 | 1.1614 | 3.91E-05 | 5.18E-08 | 5547.4978 | 4.16E-06 |
| Ptpn9 | -10.3852 | 0.8404 | 4.0787 | 2.14E-05 | 5.23E-08 | 6881.0554 | 4.18E-06 |
| Samd9l | -11.1745 | 1.334 | 2.3888 | 7.89E-02 | 5.53E-08 | 5685.486 | 4.36E-06 |
| Vwa5a | -11.0355 | 1.1384 | 25.7236 | 1.85E-05 | 6.35E-08 | 5529.6679 | 4.92E-06 |
| Ubtf | -10.3429 | 0.8068 | 17.8244 | 3.71E-05 | 6.95E-08 | 6866.3722 | 5.34E-06 |
| Zyg11b | -10.364 | 0.814 | 11.2453 | 2.17E-05 | 7.76E-08 | 6772.2462 | 5.88E-06 |
| Phc3 | -10.0603 | 0.7046 | 5.3029 | 3.51E-05 | 8.30E-08 | 7614.5226 | 6.23E-06 |
| Zhx2 | -10.5085 | 0.8784 | 3.3328 | 2.33E-05 | 8.66E-08 | 6558.7874 | 6.44E-06 |
| Slc7a2 | -10.9904 | 1.116 | 1.8805 | 3.64E-05 | 9.29E-08 | 5749.0497 | 6.85E-06 |
| Appl1 | -10.0957 | 0.7124 | 5.7761 | 2.36E-05 | 9.35E-08 | 7464.5391 | 6.87E-06 |
| Slc4a3 | -10.3646 | 0.8064 | 18.2961 | 2.12E-05 | 9.74E-08 | 6772.4809 | 7.13E-06 |
| Lin52 | -11.6056 | 1.4952 | 1.3866 | 2.62E-05 | 1.15E-07 | 4911.9606 | 8.23E-06 |
| Ttc39b | -11.081 | 1.1471 | 3.6207 | 2.58E-05 | 1.17E-07 | 5472.6274 | 8.33E-06 |
| Plekhh2 | -11.3914 | 1.3883 | 0.8113 | 2.39E-02 | 1.71E-07 | 5250.9805 | 1.16E-05 |
| Lima1 | -10.7372 | 0.9773 | 0.76 | 2.82E-05 | 1.82E-07 | 6149.0762 | 1.23E-05 |
| Brd3 | -10.5332 | 0.8603 | 6.2492 | 2.24E-05 | 1.91E-07 | 6393.391 | 1.28E-05 |
| Rps6ka5 | -11.4002 | 1.4333 | 1.6473 | 8.11E-02 | 2.19E-07 | 5544.9222 | 1.45E-05 |
| Dstyk | -10.993 | 1.0692 | 11.9833 | 2.87E-05 | 2.23E-07 | 5412.9481 | 1.46E-05 |
| Prickle2 | -11.6072 | 1.4625 | 1.2772 | 3.14E-05 | 2.24E-07 | 4848.4971 | 1.46E-05 |

**Smooth Muscle Cell: gene expression, negative correlation (β_1_) with age**

| **gene** | **Beta0** | **Beta1** | **Disp** | **Sigma** | **P.Beta** | **AIC** | **fdrP** |
| --- | --- | --- | --- | --- | --- | --- | --- |
| Ercc4 | -9.1096 | -1.9886 | 29.9402 | 1.02E-06 | 3.80E-99 | 3217.440 | 6.47E-95 |
| Sdk2 | -8.877 | -1.7655 | 0.947 | 1.60E-04 | 1.11E-76 | 4308.432 | 9.43E-73 |
| Dync1i1 | -10.2585 | -4.03 | 29.6169 | 8.87E-05 | 1.30E-46 | 572.9016 | 7.39E-43 |
| Cpox | -9.3722 | -1.561 | 0.6554 | 3.62E-05 | 8.14E-41 | 3450.984 | 3.47E-37 |
| Bex3 | -8.8533 | -0.9944 | 4.1559 | 3.34E-05 | 1.42E-36 | 6495.329 | 4.03E-33 |
| Ubb-ps | -10.5286 | -2.8321 | 2.3085 | 2.86E-05 | 3.03E-36 | 750.8064 | 7.37E-33 |
| Tpcn2 | -9.219 | -1.8777 | 0.1761 | 3.91E-05 | 3.89E-36 | 3016.925 | 8.28E-33 |
| Gap43 | -10.5615 | -3.38 | 0.2612 | 3.64E-05 | 7.45E-34 | 562.0157 | 1.41E-30 |
| Gm45609 | -10.6352 | -3.7133 | 0.2288 | 4.06E-05 | 2.38E-31 | 466.4338 | 4.05E-28 |
| Pfkfb4 | -10.0129 | -1.6925 | 0.664 | 2.61E-05 | 1.30E-28 | 2047.793 | 1.85E-25 |
| Ttll12 | -9.8035 | -2.0581 | 0.1174 | 4.63E-05 | 4.79E-27 | 1811.1 | 6.26E-24 |
| Snord104 | -10.4143 | -2.2396 | 0.2928 | 3.52E-05 | 5.14E-27 | 1111.947 | 6.26E-24 |
| Gfod2 | -9.6023 | -1.8147 | 0.1162 | 3.56E-05 | 1.32E-25 | 2338.706 | 1.50E-22 |
| Spa17 | -10.2059 | -1.667 | 1.1488 | 3.86E-05 | 8.05E-25 | 1829.469 | 8.57E-22 |
| Rny1 | -10.9479 | -3.6121 | 0.2385 | 4.04E-05 | 3.57E-24 | 373.8124 | 3.58E-21 |
| Dmac2l | -10.1428 | -1.5821 | 0.8129 | 8.17E-05 | 1.70E-23 | 1996.135 | 1.61E-20 |
| Slc2a3 | -10.0127 | -2.0805 | 0.0956 | 6.47E-05 | 1.98E-22 | 1529.567 | 1.78E-19 |
| Racgap1 | -11.1265 | -3.798 | 147746.415 | 1.94E-04 | 1.53E-21 | 304.1932 | 1.24E-18 |
| H4c14 | -10.8557 | -2.5031 | 0.2636 | 3.45E-05 | 2.02E-21 | 684.8197 | 1.57E-18 |
| Il12rb1 | -10.996 | -4.1804 | 0.2067 | 3.94E-05 | 3.60E-21 | 299.7066 | 2.67E-18 |
| Gm50455 | -11.0781 | -2.7103 | 1.1999 | 7.78E-07 | 5.66E-21 | 523.2945 | 4.02E-18 |
| Ptprn | -10.4292 | -2.3117 | 2.1372 | 1.12E-01 | 3.03E-20 | 1069.805 | 1.98E-17 |
| Dennd6b | -10.0497 | -2.3532 | 0.0477 | 4.11E-05 | 1.17E-19 | 1169.102 | 7.36E-17 |
| Gm6787 | -11.1505 | -3.4806 | 0.1085 | 4.06E-05 | 5.01E-19 | 336.8781 | 3.05E-16 |
| Batf | -11.2746 | -3.5438 | 0.1701 | 3.92E-05 | 7.42E-18 | 301.4315 | 4.22E-15 |
| H3c6 | -11.3447 | -3.1385 | 0.6863 | 3.44E-05 | 9.77E-18 | 331.6523 | 5.37E-15 |
| Mbd4 | -10.3323 | -1.7395 | 0.1399 | 3.93E-05 | 1.25E-17 | 1516.636 | 6.68E-15 |
| Slc7a14 | -11.2068 | -4.1248 | 0.1603 | 3.49E-05 | 1.52E-17 | 259.5425 | 7.87E-15 |
| Slc9a3r1 | -9.867 | -1.0749 | 2.3704 | 2.76E-05 | 7.30E-17 | 3238.361 | 3.55E-14 |
| Pcbd2 | -8.6217 | -1.1767 | 1.1261 | 6.74E-02 | 1.95E-16 | 6735.580 | 8.97E-14 |
| Gm45442 | -11.463 | -3.4615 | 2924086.52 | 9.44E-05 | 2.22E-16 | 268.1609 | 9.94E-14 |
| Dlgap2 | -11.3282 | -2.749 | 0.4224 | 3.83E-05 | 2.40E-16 | 420.1169 | 1.05E-13 |
| Mrln | -11.2782 | -2.5101 | 0.6924 | 3.65E-04 | 3.51E-16 | 490.0011 | 1.49E-13 |
| Zfp935 | -11.464 | -3.2581 | 0.3126 | 3.80E-05 | 1.10E-15 | 292.9145 | 4.46E-13 |
| 4921531C22Rik | -10.7452 | -1.6833 | 0.744 | 4.87E-05 | 1.74E-15 | 1198.277 | 6.88E-13 |
| Ddb2 | -9.826 | -1.374 | 0.1846 | 4.46E-05 | 3.78E-15 | 2706.212 | 1.40E-12 |
| H4c1 | -11.3975 | -2.6327 | 0.4857 | 4.41E-05 | 4.68E-15 | 415.6648 | 1.66E-12 |
| 1700064H15Rik | -11.1019 | -3.368 | 0.0336 | 3.95E-05 | 4.82E-15 | 345.3293 | 1.66E-12 |
| Gm48943 | -11.1753 | -4.8478 | 5766350.8 | 6.86E-05 | 4.88E-15 | NA | 1.66E-12 |
| Fhod3 | -10.3456 | -1.7814 | 0.0769 | 7.78E-05 | 1.12E-14 | 1391.297 | 3.74E-12 |
| 4930513L16Rik | -11.4094 | -4.3256 | 0.5193 | 3.85E-05 | 1.63E-14 | 209.7112 | 5.33E-12 |
| Ticrr | -11.5337 | -3.1013 | 0.2725 | 3.20E-05 | 1.76E-14 | 299.0011 | 5.67E-12 |
| Plxdc1 | -9.6222 | -4.8768 | 0.0062 | 8.85E-05 | 1.94E-14 | 318.1192 | 6.13E-12 |
| U2af1 | -9.4348 | -0.7545 | 450.3833 | 2.16E-13 | 2.18E-14 | 5161.631 | 6.75E-12 |
| Znrf2 | -9.6563 | -0.8895 | 1.8 | 3.82E-05 | 2.43E-14 | 4180.499 | 7.31E-12 |
| Edn1 | -10.9649 | -2.5304 | 0.0492 | 4.79E-05 | 2.45E-14 | 584.179 | 7.31E-12 |
| Fgd1 | -9.8162 | -1.0077 | 0.7964 | 3.18E-05 | 2.53E-14 | 3534.584 | 7.43E-12 |
| Gm27151 | -11.039 | -1.8923 | 0.462 | 4.28E-05 | 6.90E-14 | 837.5217 | 1.96E-11 |
| AC113970.1 | -11.4556 | -4.2791 | 0.2379 | 3.95E-05 | 7.11E-14 | 208.296 | 1.99E-11 |
| Uckl1 | -9.4214 | -1.2288 | 1.3454 | 9.01E-02 | 1.01E-13 | 4023.754 | 2.78E-11 |

**Myeloid: gene expression, positive correlation (β_1_) with age**

| **gene** | **Beta0** | **Beta1** | **Disp** | **Sigma** | **P.Beta** | **AIC** | **fdrP** |
| --- | --- | --- | --- | --- | --- | --- | --- |
| Itih5 | -9.2953 | 0.731 | 0.7305 | 7.02E-05 | 5.97E-08 | 4587.9208 | 3.52E-05 |
| Vps13d | -10.4628 | 1.1157 | 8.1092 | 2.70E-05 | 8.71E-08 | 2858.1634 | 4.97E-05 |
| C130026I21Rik | -10.1638 | 1.2109 | 1.2616 | 1.22E-01 | 1.21E-07 | 3724.8345 | 5.87E-05 |
| Prelp | -9.1327 | 0.7997 | 1.6873 | 1.00E-01 | 1.50E-07 | 5210.2008 | 7.07E-05 |
| Pknox2 | -11.4461 | 1.705 | 1.2443 | 6.29E-05 | 5.65E-07 | 2279.0031 | 2.22E-04 |
| Kdm2a | -9.7268 | 0.7347 | 4.9688 | 1.94E-02 | 1.22E-06 | 3565.9903 | 4.28E-04 |
| Mllt10 | -9.5271 | 0.7147 | 4.5736 | 6.15E-02 | 3.08E-06 | 3951.8103 | 8.48E-04 |
| Mdm4 | -10.8094 | 1.1451 | 4.4466 | 3.32E-05 | 3.89E-06 | 2400.4257 | 9.98E-04 |
| Trim8 | -10.384 | 0.93 | 4.7436 | 4.20E-05 | 4.07E-06 | 2736.8369 | 9.98E-04 |
| Ngf | -10.5408 | 1.0882 | 0.4094 | 6.06E-05 | 4.15E-06 | 2739.5772 | 9.98E-04 |
| Lipa | -10.5112 | 0.9936 | 2.5004 | 5.12E-05 | 4.16E-06 | 2659.9914 | 9.98E-04 |
| Klhl29 | -10.0326 | 0.8381 | 0.7139 | 3.16E-05 | 4.36E-06 | 3236.8469 | 1.03E-03 |
| Rab7b | -11.0042 | 1.2496 | 3.138 | 2.85E-05 | 4.78E-06 | 2159.1945 | 1.11E-03 |
| Gypc | -11.8501 | 1.8889 | 10.4464 | 2.00E-02 | 5.37E-06 | 1928.2471 | 1.23E-03 |
| Ilrun | -10.55 | 0.9862 | 663.4363 | 3.54E-14 | 5.53E-06 | 2526.967 | 1.25E-03 |
| Map2k5 | -11.095 | 1.3961 | 3.7014 | 1.09E-01 | 5.77E-06 | 2406.5062 | 1.29E-03 |
| Uba6 | -10.8086 | 1.1133 | 4.7893 | 3.33E-05 | 7.14E-06 | 2307.3531 | 1.55E-03 |
| Ncoa1 | -9.3136 | 0.6958 | 2.7014 | 8.82E-02 | 9.06E-06 | 4525.6458 | 1.85E-03 |
| Zfp950 | -10.7511 | 1.1781 | 2.259 | 6.62E-02 | 9.24E-06 | 2577.1546 | 1.86E-03 |
| Zscan26 | -11.3383 | 1.4025 | 14.191 | 1.55E-03 | 1.20E-05 | 2014.4047 | 2.26E-03 |
| Bmp6 | -10.765 | 1.1766 | 0.2241 | 5.05E-05 | 1.36E-05 | 2385.6943 | 2.51E-03 |
| Kdm5b | -10.2417 | 0.813 | 3.4207 | 4.09E-05 | 1.77E-05 | 2793.8488 | 3.05E-03 |
| Pou2f1 | -10.6466 | 0.98 | 4.1355 | 9.01E-05 | 1.98E-05 | 2408.7022 | 3.35E-03 |
| Matn2 | -10.738 | 1.1242 | 0.2826 | 3.11E-05 | 2.22E-05 | 2428.9485 | 3.63E-03 |
| Insr | -10.5026 | 0.9189 | 2.0363 | 3.57E-05 | 2.27E-05 | 2602.4958 | 3.68E-03 |
| Csnk1g3 | -9.9774 | 0.7 | 8.7619 | 1.37E-02 | 2.75E-05 | 3011.2309 | 4.14E-03 |
| G3bp2 | -10.0296 | 0.7099 | 6.7698 | 2.40E-05 | 2.95E-05 | 2979.3657 | 4.35E-03 |
| Tlk2 | -10.2761 | 0.804 | 3.2147 | 2.13E-05 | 3.04E-05 | 2715.7473 | 4.46E-03 |
| Dennd4c | -10.1093 | 0.8373 | 3.1187 | 3.98E-02 | 3.54E-05 | 3083.1892 | 4.96E-03 |
| Creb3l2 | -9.7201 | 0.7272 | 3.7087 | 8.25E-02 | 3.96E-05 | 3633.6731 | 5.28E-03 |
| Tshz3 | -10.1449 | 0.739 | 3.1904 | 3.80E-05 | 4.62E-05 | 2841.7856 | 5.88E-03 |
| Nlk | -10.4242 | 0.8548 | 1.4126 | 3.78E-05 | 4.91E-05 | 2592.7975 | 6.10E-03 |
| Trabd2b | -11.8958 | 1.9617 | 0.2965 | 2.28E-01 | 5.36E-05 | 2109.9989 | 6.52E-03 |
| Pdlim4 | -10.1842 | 0.75 | 2.0264 | 3.44E-05 | 5.46E-05 | 2720.3569 | 6.59E-03 |
| Fgd5 | -11.4683 | 1.6298 | 0.9704 | 2.05E-01 | 5.89E-05 | 2302.1267 | 7.05E-03 |
| Myoc | -11.2976 | 1.8002 | 0.0312 | 7.01E-05 | 6.05E-05 | 1485.7066 | 7.20E-03 |
| Hectd4 | -10.9339 | 1.055 | 8.4818 | 3.15E-05 | 6.19E-05 | 2028.3943 | 7.31E-03 |
| Gm10125 | -11.4442 | 1.3808 | 0.6395 | 3.90E-05 | 6.58E-05 | 1864.4436 | 7.72E-03 |
| Zmym2 | -10.1154 | 0.7057 | 3.7801 | 2.99E-05 | 7.64E-05 | 2860.2696 | 8.84E-03 |
| Dop1b | -12.032 | 1.7859 | 3.7786 | 3.29E-05 | 7.77E-05 | 1604.5663 | 8.92E-03 |
| Acvr1 | -10.2774 | 0.7606 | 3.5165 | 3.59E-05 | 7.93E-05 | 2680.5406 | 8.93E-03 |
| Dnajc13 | -10.4639 | 0.8598 | 6.1451 | 2.53E-02 | 8.01E-05 | 2512.5002 | 8.95E-03 |
| Chm | -10.8098 | 0.98 | 4.3158 | 5.48E-05 | 8.14E-05 | 2180.2088 | 9.03E-03 |
| Sparcl1 | -10.1182 | 1.0653 | 0.1639 | 1.50E-01 | 8.20E-05 | 3066.6461 | 9.04E-03 |
| Sacm1l | -10.3088 | 0.757 | 7.9843 | 9.70E-05 | 9.92E-05 | 2568.4062 | 1.05E-02 |
| Arrb1 | -10.8684 | 0.9954 | 4.6147 | 3.00E-05 | 1.02E-04 | 2188.6169 | 1.05E-02 |
| Slc10a6 | -10.3223 | 0.8464 | 0.4016 | 4.42E-05 | 1.04E-04 | 2702.0686 | 1.05E-02 |
| Ap3d1 | -10.8079 | 0.9586 | 129.3556 | 1.94E-14 | 1.05E-04 | NA | 1.05E-02 |
| Dicer1 | -10.643 | 0.8895 | 4.3507 | 3.30E-05 | 1.10E-04 | 2294.5008 | 1.09E-02 |
| Rabl6 | -10.6496 | 0.8906 | 3.6543 | 3.02E-05 | 1.10E-04 | 2297.6298 | 1.09E-02 |

**Myeloid: gene expression, negative correlation (β_1_) with age**

| **gene** | **Beta0** | **Beta1** | **Disp** | **Sigma** | **P.Beta** | **AIC** | **fdrP** |
| --- | --- | --- | --- | --- | --- | --- | --- |
| Ttll12 | -8.6382 | -2.6566 | 0.0508 | 8.62E-05 | 4.16E-23 | 1049.425 | 6.87E-19 |
| Itm2a | -9.3468 | -1.4885 | 0.2626 | 4.56E-05 | 9.30E-16 | 1478.323 | 7.69E-12 |
| Dmac2l | -9.8757 | -1.6895 | 0.3593 | 6.85E-05 | 1.73E-15 | 908.7069 | 7.96E-12 |
| Cpox | -9.4389 | -1.2957 | 0.5889 | 5.05E-05 | 1.92E-15 | 1527.788 | 7.96E-12 |
| Slco2a1 | -9.803 | -2.1622 | 0.0708 | 8.67E-04 | 2.39E-13 | 711.7543 | 7.90E-10 |
| H4f16 | -10.6921 | -2.7873 | 0.1687 | 6.66E-05 | 3.91E-12 | 307.1085 | 9.24E-09 |
| Nudt1 | -9.6817 | -2.2645 | 0.043 | 4.53E-05 | 6.75E-12 | 657.9736 | 1.40E-08 |
| Fnbp1l | -9.6907 | -1.6372 | 0.0915 | 5.73E-05 | 6.18E-11 | 964.9275 | 1.14E-07 |
| 1700003G18Rik | -10.2742 | -3.1133 | 0.0205 | 1.01E-04 | 4.43E-10 | 290.3885 | 6.66E-07 |
| Bphl | -9.6899 | -0.9612 | 78.086 | 6.41E-08 | 6.25E-10 | 1520.678 | 8.61E-07 |
| Tnfrsf12a | -9.0814 | -0.7681 | 1.0615 | 7.69E-05 | 9.07E-10 | 2501.312 | 1.15E-06 |
| Osgep | -9.1346 | -0.8463 | 1.0297 | 4.66E-02 | 1.28E-09 | 2336.160 | 1.51E-06 |
| Cdh18 | -10.4934 | -2.0258 | 0.1079 | 6.55E-05 | 2.09E-09 | 484.2409 | 2.31E-06 |
| A2ml1 | -11.1743 | -2.8484 | 0.4034 | 4.56E-05 | 4.78E-09 | 204.1599 | 4.70E-06 |
| Grid2 | -10.4222 | -1.3774 | 3524199.86 | 4.31E-12 | 4.83E-09 | NA | 4.70E-06 |
| Dpagt1 | -9.6713 | -1.1054 | 0.2521 | 4.55E-05 | 5.15E-09 | 1378.401 | 4.70E-06 |
| Gm45510 | -11.2432 | -2.6514 | 4545327.03 | 5.44E-05 | 1.16E-08 | 213.7092 | 9.42E-06 |
| Otx2os1 | -10.8512 | -3.5019 | 0.0322 | 7.17E-05 | 1.20E-08 | 190.442 | 9.42E-06 |
| Ptprn | -10.4707 | -4.4676 | 1.3723 | 3.40E-01 | 1.48E-08 | 264.7124 | 1.11E-05 |
| Pik3c2a | -8.9811 | -0.6972 | 1.3703 | 2.54E-02 | 1.69E-08 | 2764.118 | 1.19E-05 |
| Ercc4 | -10.0155 | -1.0887 | 2.8428 | 3.24E-02 | 1.73E-08 | 1112.549 | 1.19E-05 |
| Slc9a3r1 | -10.1688 | -1.5825 | 0.3318 | 7.78E-02 | 1.93E-08 | 771.8676 | 1.28E-05 |
| 4930512B01Rik | -10.4978 | -2.8067 | 0.0196 | 6.20E-05 | 4.77E-08 | 280.4491 | 3.04E-05 |
| Cldn10 | -10.447 | -1.3724 | 0.5746 | 3.37E-05 | 5.86E-08 | 729.5842 | 3.52E-05 |
| Larp7 | -9.5924 | -0.8176 | 1.8224 | 2.60E-05 | 9.39E-08 | 1790.066 | 5.01E-05 |
| Dstyk | -9.1945 | -0.7301 | 0.7726 | 7.15E-05 | 9.40E-08 | 2368.988 | 5.01E-05 |
| Gap43 | -10.4514 | -2.0713 | 0.0797 | 1.83E-01 | 1.06E-07 | 474.3637 | 5.37E-05 |
| Nwd2 | -11.4438 | -2.7384 | 1311581.87 | 2.09E-46 | 2.04E-07 | 169.6892 | 9.37E-05 |
| Gps2 | -9.4373 | -0.6985 | 11.2547 | 3.52E-05 | 2.23E-07 | 2097.352 | 9.96E-05 |
| Rpl37-ps1 | -11.4438 | -3.4315 | 2786497.26 | 9.92E-05 | 2.64E-07 | 132.4164 | 1.15E-04 |
| Tmprss9 | -11.3566 | -2.5308 | 0.2295 | 4.03E-05 | 2.74E-07 | 200.4893 | 1.16E-04 |
| Tmem108 | -10.4145 | -2.8558 | 0.0133 | 1.72E-04 | 3.22E-07 | 269.5217 | 1.33E-04 |
| Ank3 | -9.5384 | -1.8611 | 0.0252 | 1.22E-04 | 4.84E-07 | 744.6009 | 1.95E-04 |
| Tulp2 | -11.0756 | -1.9082 | 0.1366 | 6.05E-05 | 1.10E-06 | 323.7713 | 4.05E-04 |
| Dcps | -9.986 | -1.0185 | 0.3151 | 5.73E-04 | 1.36E-06 | 1193.286 | 4.69E-04 |
| Cgref1 | -11.1348 | -1.9863 | 0.1283 | 4.50E-05 | 1.52E-06 | 307.8059 | 5.13E-04 |
| Gm26555 | -11.1753 | -2.1371 | 0.0824 | 1.34E-04 | 1.60E-06 | 268.7858 | 5.31E-04 |
| Endog | -10.1446 | -0.9316 | 3476670.23 | 6.96E-26 | 1.64E-06 | 1144.791 | 5.31E-04 |
| Gm15567 | -11.3827 | -3.1842 | 0.0365 | 7.47E-05 | 1.81E-06 | 148.91 | 5.75E-04 |
| Dcc | -11.5616 | -2.4665 | 1758633.03 | 3.83E-15 | 1.88E-06 | 185.0384 | 5.87E-04 |
| Gm14798 | -10.9343 | -1.5757 | 0.306 | 3.21E-02 | 2.48E-06 | 435.8414 | 7.59E-04 |
| Ccdc171 | -9.3632 | -0.7073 | 0.5999 | 9.61E-05 | 2.90E-06 | 2205.983 | 8.48E-04 |
| Ezh2 | -9.793 | -0.9919 | 0.4369 | 9.12E-02 | 2.99E-06 | 1404.069 | 8.48E-04 |
| Nedd9 | -8.8019 | -1.049 | 0.2528 | 1.43E-01 | 3.00E-06 | 2529.770 | 8.48E-04 |
| Dhrs7b | -9.8435 | -0.8112 | 1.2263 | 3.02E-05 | 3.06E-06 | 1490.822 | 8.48E-04 |
| 1700112J16Rik | -11.5616 | -2.3329 | 6895532.5 | 3.25E-18 | 3.07E-06 | 185.0453 | 8.48E-04 |
| Ak7 | -11.4438 | -2.0452 | 2300573.4 | 1.73E-15 | 3.51E-06 | 235.0312 | 9.39E-04 |
| Sumf2 | -10.0564 | -0.8676 | 4.6224 | 5.86E-05 | 3.55E-06 | 1254.084 | 9.39E-04 |
| Acd | -10.307 | -1.0095 | 1.7319 | 4.17E-05 | 3.58E-06 | 982.9097 | 9.39E-04 |
| Fzd5 | -10.2974 | -1.1468 | 1.0125 | 1.03E-01 | 4.04E-06 | 908.0828 | 9.98E-04 |

**Perivascular Macrophage: gene expression, positive correlation (β_1_) with age**

| **gene** | **Beta0** | **Beta1** | **Disp** | **Sigma** | **P.Beta** | **AIC** | **fdrP** |
| --- | --- | --- | --- | --- | --- | --- | --- |
| Wdr33 | -10.2491 | 1.16E+14 | 7.743034 | 4.13E-05 | 7.91E-07 | 1741.861 | 0.000803 |
| C130026I21Rik | -9.82924 | 1.36E+14 | 2.324261 | 0.300124 | 1.34E-05 | 2719.183 | 0.006198 |
| Pacs1 | -9.85837 | 8.97E+14 | 0.88233 | 3.72E-05 | 1.50E-05 | 2009.327 | 0.006533 |
| Slc23a2 | -9.99414 | 9.85E+14 | 2.399098 | 0.137919 | 5.69E-05 | 1947.387 | 0.015221 |
| Lrp10 | -10.6268 | 1.14E+14 | 8.734029 | 5.66E-05 | 6.33E-05 | 1386.645 | 0.016607 |
| Cul1 | -9.71421 | 7.52E+12 | 1.987161 | 8.52E-05 | 6.60E-05 | 2026.611 | 0.017036 |
| Gm20732 | -11.804 | 2.01E+14 | 7.727827 | 3.43E-05 | 7.04E-05 | 1111.904 | 0.017856 |
| Pde7b | -9.73319 | 1.29E+13 | 0.595869 | 0.311457 | 7.21E-05 | 2781.896 | 0.017985 |
| Mier1 | -9.4094 | 7.24E+14 | 25.16253 | 0.071992 | 0.000102 | 2293.79 | 0.022156 |
| Hs2st1 | -10.7952 | 1.18E+14 | 3.939956 | 0.000121 | 0.000128 | 1296.501 | 0.025307 |
| Oas2 | -11.8703 | 2.2E+14 | 2.190779 | 0.364483 | 0.000207 | 1418.869 | 0.033151 |
| Exoc6 | -11.2554 | 1.43E+14 | 1.169288 | 5.55E-05 | 0.000238 | 1133.538 | 0.035821 |
| Lonp2 | -11.2591 | 1.6E+14 | 6.838344 | 0.212683 | 0.000257 | 1330.589 | 0.036205 |
| Dst | -9.91393 | 8.14E+14 | 0.428516 | 4.18E-05 | 0.000259 | 1810.248 | 0.036205 |
| Ecpas | -10.1956 | 1.04E+14 | 45.21069 | 0.143044 | 0.000289 | 1762.998 | 0.038428 |
| Ifi209 | -11.2522 | 1.47E+14 | 1.448757 | 0.119754 | 0.00029 | 1180.27 | 0.038428 |
| Pnn | -10.7949 | 1.2E+14 | 6.593619 | 0.129663 | 0.000332 | 1358.656 | 0.04013 |
| R3hdm1 | -10.3014 | 8.4E+14 | 4.039403 | 2.94E-05 | 0.000588 | 1474.345 | 0.060437 |
| Slfn5 | -9.86174 | 1.05E+14 | 1.578997 | 0.279388 | 0.000594 | 2315.354 | 0.060648 |
| Maea | -10.7935 | 1.05E+14 | 13.75695 | 0.001059 | 0.00067 | 1201.315 | 0.065349 |
| Fbxo42 | -12.4969 | 2.41E+14 | 3.761251 | 5.18E-05 | 0.000724 | 900.3149 | 0.067581 |
| Cemip2 | -10.4114 | 8.88E+14 | 1.420468 | 3.89E-05 | 0.000734 | 1407.342 | 0.067848 |
| Nrf1 | -10.2068 | 7.99E+14 | 1.470844 | 5.01E-05 | 0.000742 | 1521.642 | 0.068012 |
| Acox3 | -11.4017 | 1.4E+14 | 3.706061 | 4.00E-05 | 0.000765 | 1007.529 | 0.069255 |
| Iigp1 | -10.9597 | 1.67E+14 | 0.167758 | 0.393213 | 0.000774 | 1427.614 | 0.069255 |
| Phka2 | -11.2481 | 1.3E+14 | 3.418387 | 9.14E-05 | 0.000781 | 1043.493 | 0.069529 |
| Plscr4 | -10.8061 | 1.06E+14 | 1.006683 | 3.17E-05 | 0.000815 | 1213.145 | 0.070828 |
| Parp12 | -10.8946 | 1.16E+14 | 4.28709 | 0.12357 | 0.000844 | 1242.307 | 0.071465 |
| Cxcl16 | -9.76465 | 8.85E+14 | 0.565257 | 0.189241 | 0.000864 | 2052.268 | 0.071846 |
| Rsad2 | -12.0591 | 2.03E+14 | 0.135854 | 4.74E-05 | 0.000877 | 893.9832 | 0.07257 |
| Mdfic | -9.55998 | 8.68E+13 | 3.168202 | 0.233776 | 0.00089 | 2396.548 | 0.073233 |
| Palld | -11.5827 | 1.53E+14 | 0.712142 | 3.71E-05 | 0.000909 | 960.2112 | 0.073697 |
| Fyb | -9.17789 | 1.25E+14 | 2.565098 | 0.438688 | 0.000922 | 3687.861 | 0.073697 |
| Ash1l | -9.62112 | 9.63E+14 | 24.76211 | 0.282819 | 0.001008 | 2367.463 | 0.077836 |
| Chordc1 | -10.7082 | 9.76E+14 | 2.675481 | 5.23E-05 | 0.001078 | 1219.672 | 0.081218 |
| Odr4 | -10.7056 | 9.64E+14 | 11.35319 | 0.000217 | 0.001132 | 1198.247 | 0.084068 |
| Ube2e2 | -9.98725 | 8.04E+14 | 0.524304 | 0.103711 | 0.001145 | 1753.223 | 0.084582 |
| Btaf1 | -9.42934 | 7.9E+14 | 3.14584 | 0.219364 | 0.001179 | 2281.038 | 0.086249 |
| Ccl8 | -8.71364 | 2.27E+14 | 0.306255 | 0.878004 | 0.001211 | 6144.239 | 0.087736 |
| Ncoa7 | -10.2435 | 9.32E+14 | 4.943974 | 0.168955 | 0.001293 | 1613.352 | 0.08966 |
| Pja2 | -10.3001 | 7.81E+14 | 7.241713 | 3.44E-05 | 0.00135 | 1394.965 | 0.0912 |
| Ccdc6 | -11.8048 | 1.62E+14 | 14650655 | 1.17E-22 | 0.001374 | 859.6717 | 0.0912 |
| Gpr137b-ps | -11.1113 | 1.16E+14 | 2.506798 | 7.27E-05 | 0.001376 | 1043.583 | 0.0912 |
| Ddx58 | -10.7872 | 1.06E+13 | 1.996264 | 0.097915 | 0.001395 | 1265.26 | 0.091887 |
| Tbrg1 | -10.8885 | 1.03E+14 | 9701360 | 3.29E-27 | 0.00146 | NA | 0.093738 |
| Pld1 | -10.539 | 8.95E+14 | 1.154557 | 3.44E-05 | 0.001475 | 1351.005 | 0.094314 |
| Cpq | -9.52281 | 7.39E+14 | 2.737441 | 0.185975 | 0.001497 | 2256.159 | 0.094456 |
| Csnk1g1 | -10.6458 | 1.07E+14 | 4.44131 | 0.186143 | 0.001498 | 1410.502 | 0.094456 |
| Tgs1 | -10.5494 | 8.79E+14 | 3.112608 | 8.21E-05 | 0.001504 | 1287.172 | 0.094456 |
| Arap1 | -11.2451 | 1.23E+14 | 1.724643 | 3.05E-05 | 0.001537 | 995.6132 | 0.094456 |

**Perivascular Macrophage: gene expression, negative correlation (β_1_) with age**

| **gene** | **Beta0** | **Beta1** | **Disp** | **Sigma** | **P.Beta** | **AIC** | **fdrP** |
| --- | --- | --- | --- | --- | --- | --- | --- |
| Nr1d1 | -9.76518 | -1.9E+14 | 1.120959 | 3.52E-05 | 7.49E-14 | 534.7277 | 1.14E-09 |
| Gm12236 | -10.2667 | -3.2E+14 | 0.737129 | 9.52E-05 | 1.32E-11 | 223.5961 | 1.01E-07 |
| Cdkl4 | -10.0196 | -1.8E+13 | 0.70504 | 4.98E-05 | 4.08E-10 | 469.6472 | 2.07E-06 |
| Nedd4l | -8.73245 | -9.4E+13 | 0.405558 | 4.31E-05 | 4.17E-09 | 1597.167 | 1.16E-05 |
| Cirbp | -9.50218 | -1.1E+14 | 3863530 | 1.69E-21 | 4.56E-09 | NA | 1.16E-05 |
| Als2 | -9.45906 | -1.6E+14 | 0.113399 | 9.72E-05 | 5.38E-09 | 691.7965 | 1.17E-05 |
| Tmem189 | -10.6572 | -2.6E+14 | 0.2499 | 4.81E-05 | 5.81E-08 | 199.4201 | 0.000111 |
| Rapgef4 | -9.34593 | -5.1E+14 | 0.006382 | 7.01E-05 | 1.18E-07 | 150.4907 | 0.00018 |
| Entr1 | -9.47482 | -9.5E+14 | 8.606352 | 0.006897 | 1.71E-07 | 1026.62 | 0.000236 |
| Dut | -10.3083 | -1.8E+14 | 0.192186 | 0.000146 | 2.27E-07 | 354.0351 | 0.000288 |
| Tceal8 | -9.50926 | -9.6E+14 | 2.891246 | 3.28E-05 | 2.98E-07 | 958.3001 | 0.000348 |
| Anp32e | -8.92434 | -8.3E+14 | 1.229784 | 0.076743 | 9.75E-07 | 1536.561 | 0.000928 |
| Vtcn1 | -10.2134 | -2.4E+14 | 0.037738 | 5.37E-05 | 1.07E-06 | 273.7492 | 0.00096 |
| Myo15 | -10.7384 | -2.1E+14 | 0.219962 | 0.000158 | 1.51E-06 | 230.4769 | 0.001259 |
| Rexo1 | -9.47406 | -1E+14 | 3.668727 | 0.126172 | 1.57E-06 | 996.9921 | 0.001259 |
| 6030469F06Rik | -10.5343 | -2.8E+14 | 0.276984 | 0.39388 | 2.25E-06 | 208.0938 | 0.001711 |
| Pdpn | -10.498 | -1.9E+14 | 0.136644 | 5.27E-05 | 2.88E-06 | 303.0358 | 0.002088 |
| Gm43305 | -10.6197 | -1.8E+14 | 0.513826 | 6.64E-05 | 3.21E-06 | 308.6423 | 0.002219 |
| Tmem185b | -10.0597 | -1.2E+14 | 2.230677 | 4.25E-05 | 4.00E-06 | 625.9755 | 0.002646 |
| AI506816 | -10.0127 | -1.1E+14 | 20.96745 | 8.48E-05 | 4.53E-06 | 648.7212 | 0.002874 |
| Polr2i | -9.38496 | -7.7E+14 | 8.851386 | 3.59E-08 | 5.29E-06 | 1143.206 | 0.003218 |
| Plk2 | -8.18476 | -1E+14 | 1.136623 | 0.240145 | 6.40E-06 | 2129.752 | 0.003747 |
| Mcm7 | -10.2038 | -1.4E+14 | 0.190212 | 5.39E-05 | 8.71E-06 | 470.784 | 0.004907 |
| Ndst3 | -10.2887 | -1.6E+14 | 0.092934 | 5.83E-05 | 1.01E-05 | 384.6355 | 0.00511 |
| Dnmt1 | -9.71662 | -9.1E+14 | 2.825995 | 4.62E-05 | 1.14E-05 | 884.3698 | 0.005604 |
| Ints2 | -10.057 | -1.1E+14 | 7.094496 | 0.000169 | 1.25E-05 | 626.386 | 0.005967 |
| Suclg2 | -9.48424 | -8.7E+14 | 1.210842 | 0.032862 | 1.46E-05 | 1015.87 | 0.006533 |
| Cacnb4 | -10.6336 | -1.9E+14 | 0.081794 | 4.46E-05 | 1.66E-05 | 265.2866 | 0.007036 |
| Nkain3 | -10.961 | -2.2E+13 | 0.483763 | 5.74E-05 | 2.06E-05 | 207.1612 | 0.008482 |
| Cux2 | -10.9241 | -2.2E+14 | 0.077162 | 4.82E-05 | 2.22E-05 | 185.3154 | 0.008898 |
| Gpr158 | -10.9506 | -2.1E+14 | 0.149672 | 0.000114 | 2.38E-05 | 195.9501 | 0.008905 |
| Catsper2 | -9.80053 | -2.7E+14 | 0.011431 | 6.21E-05 | 2.42E-05 | 235.2916 | 0.008905 |
| Taf1c | -10.7048 | -1.5E+14 | 2.341834 | 3.21E-05 | 2.52E-05 | 296.4804 | 0.008905 |
| Zbtb24 | -10.3578 | -1.2E+14 | 5274390 | 1.03E-06 | 2.58E-05 | 454.9864 | 0.008939 |
| Gm15787 | -10.7904 | -1.7E+14 | 0.658435 | 4.13E-05 | 2.86E-05 | 272.1116 | 0.009475 |
| Csf3 | -10.9647 | -3.6E+14 | 0.03974 | 0.000113 | 3.04E-05 | 120.7375 | 0.009737 |
| Ndufs3 | -9.07606 | -1.3E+14 | 0.203509 | 0.217726 | 3.07E-05 | 1047.024 | 0.009737 |
| Dcaf15 | -10.6261 | -1.4E+14 | 4454851 | 2.15E-05 | 3.72E-05 | 355.762 | 0.01156 |
| Shmt1 | -9.96433 | -2.1E+14 | 0.078422 | 0.214357 | 3.92E-05 | 382.49 | 0.011944 |
| Gm5586 | -10.9938 | -1.8E+14 | 2414958 | 4.84E-28 | 4.20E-05 | NA | 0.012327 |
| Mir692-1 | -10.0645 | -4.6E+14 | 0.821117 | 0.971475 | 4.21E-05 | 342.19 | 0.012327 |
| Mir320 | -10.8328 | -4.4E+14 | 0.048367 | 4.39E-05 | 4.76E-05 | 115.5813 | 0.013654 |
| Rcl1 | -10.7062 | -1.4E+14 | 6240705 | 6.13E-27 | 5.05E-05 | NA | 0.01406 |
| Il31ra | -9.85344 | -8.9E+13 | 2.746239 | 0.000145 | 5.08E-05 | 842.9278 | 0.01406 |
| Pou6f2 | -11.1474 | -3E+14 | 0.041697 | 0.000232 | 7.53E-05 | 117.9871 | 0.018475 |
| Pbdc1 | -9.38878 | -1.1E+14 | 1.537566 | 0.244164 | 9.04E-05 | 967.4087 | 0.02106 |
| Pde11a | -11.3993 | -3.2E+14 | 2611950 | 1.92E-20 | 9.22E-05 | NA | 0.02106 |
| Fam207a | -10.0451 | -9.7E+14 | 1.09593 | 3.32E-05 | 9.25E-05 | 664.1323 | 0.02106 |
| Nipsnap3b | -9.63535 | -7.6E+13 | 2.342757 | 2.75E-05 | 9.27E-05 | 971.2015 | 0.02106 |

**Endothelial Cell: gene expression, positive correlation (β_1_) with age**

| **gene** | **Beta0** | **Beta1** | **Disp** | **Sigma** | **P.Beta** | **AIC** | **fdrP** |
| --- | --- | --- | --- | --- | --- | --- | --- |
| Ltbp1 | -9.8737 | 1.0862 | 0.2587 | 9.79E-05 | 8.38E-21 | 4292.4096 | 7.33E-17 |
| Chrm3 | -10.4814 | 1.7225 | 0.1218 | 1.14E-01 | 7.52E-17 | 3677.8082 | 2.63E-13 |
| Selenop | -8.5193 | 1.0215 | 0.6718 | 1.24E-01 | 8.04E-15 | 9149.1037 | 2.34E-11 |
| Slco3a1 | -10.7175 | 1.5002 | 0.2914 | 1.67E-01 | 4.20E-11 | 3162.0626 | 4.90E-08 |
| Stbd1 | -11.5977 | 1.309 | 0.5659 | 3.54E-04 | 1.06E-09 | 1553.2323 | 1.03E-06 |
| Ubash3b | -11.7315 | 1.5603 | 0.099 | 6.80E-02 | 4.27E-09 | 1545.0672 | 3.55E-06 |
| Prelp | -11.839 | 1.9135 | 0.1696 | 2.11E-01 | 6.72E-09 | 1781.6542 | 4.97E-06 |
| Stard8 | -10.9684 | 1.0216 | 1.5107 | 4.59E-02 | 6.82E-09 | 2119.0936 | 4.97E-06 |
| Emb | -11.0889 | 1.0495 | 0.205 | 2.85E-05 | 1.34E-08 | 1901.0334 | 8.68E-06 |
| Foxo3 | -10.4778 | 0.7462 | 0.9349 | 3.81E-05 | 1.46E-08 | 2640.3614 | 9.12E-06 |
| Gpd2 | -10.5518 | 0.7664 | 0.5957 | 5.28E-05 | 3.59E-08 | 2509.3174 | 1.90E-05 |
| Sh3tc1 | -11.7102 | 1.2588 | 0.4695 | 3.26E-05 | 3.97E-08 | 1407.8605 | 2.04E-05 |
| Lin52 | -11.0312 | 0.939 | 0.4785 | 5.25E-05 | 4.47E-08 | 1920.5568 | 2.24E-05 |
| Hipk2 | -10.9572 | 0.8698 | 1.3283 | 4.35E-02 | 2.26E-07 | 1966.5143 | 8.24E-05 |
| Cdc42ep1 | -10.4267 | 0.7173 | 5.401 | 6.25E-02 | 3.21E-07 | 2631.1639 | 1.10E-04 |
| Gja1 | -11.5829 | 1.672 | 0.1116 | 2.27E-01 | 3.48E-07 | 1849.261 | 1.15E-04 |
| Akap12 | -10.9498 | 1.6299 | 0.335 | 2.86E-01 | 5.33E-07 | 2998.5377 | 1.64E-04 |
| Unc5b | -10.5635 | 0.6998 | 0.6243 | 1.82E-04 | 5.69E-07 | 2360.8947 | 1.72E-04 |
| Thy1 | -13.703 | 3.0131 | 0.0469 | 6.65E-05 | 7.61E-07 | 904.0665 | 2.22E-04 |
| Il33 | -11.6919 | 1.2797 | 0.073 | 4.80E-05 | 8.99E-07 | 1305.154 | 2.52E-04 |
| Fmnl2 | -10.4992 | 1.4175 | 0.2286 | 2.57E-01 | 9.06E-07 | 3561.3818 | 2.52E-04 |
| Dlg5 | -11.4131 | 0.9988 | 0.2912 | 4.50E-05 | 1.15E-06 | 1470.6879 | 3.06E-04 |
| Pcp4l1 | -12.0747 | 1.3889 | 0.1086 | 4.30E-05 | 1.60E-06 | 1123.0321 | 3.91E-04 |
| Atoh8 | -12.1887 | 1.5334 | 0.6037 | 9.18E-02 | 1.61E-06 | 1207.2186 | 3.91E-04 |
| Hivep3 | -11.2598 | 1.1941 | 0.2655 | 1.35E-01 | 3.03E-06 | 1867.9604 | 6.71E-04 |
| Npr2 | -11.1479 | 0.9032 | 1.0461 | 6.97E-02 | 3.28E-06 | 1769.0849 | 7.01E-04 |
| Smo | -11.4111 | 0.904 | 3.2394 | 1.49E-04 | 4.34E-06 | 1430.6875 | 8.82E-04 |
| Tbx20 | -11.5629 | 1.5241 | 0.1173 | 2.17E-01 | 4.42E-06 | 1706.8904 | 8.83E-04 |
| Maf | -11.1393 | 1.1805 | 0.0335 | 5.75E-05 | 4.54E-06 | 1548.4756 | 8.92E-04 |
| Zfp74 | -12.7691 | 1.8977 | 0.473 | 1.67E-01 | 4.73E-06 | 994.8996 | 9.09E-04 |
| Klhl29 | -11.2074 | 0.9195 | 0.1684 | 6.38E-05 | 5.29E-06 | 1626.8505 | 9.91E-04 |
| Pard6g | -13.5426 | 2.93 | 0.0872 | 3.62E-01 | 6.97E-06 | 1138.3367 | 1.23E-03 |
| Ccl21a | -14.9344 | 5.371 | 0.0027 | 1.76E-04 | 8.29E-06 | 500.9525 | 1.37E-03 |
| Pvt1 | -11.1261 | 1.0346 | 0.6061 | 1.30E-01 | 8.42E-06 | 1902.3245 | 1.38E-03 |
| Comp | -12.7877 | 1.9181 | 0.0377 | 5.49E-05 | 1.03E-05 | 844.313 | 1.64E-03 |
| Ifit1 | -12.2564 | 1.6398 | 0.0762 | 1.91E-01 | 1.84E-05 | 1120.375 | 2.56E-03 |
| Dusp2 | -11.9079 | 1.3348 | 0.0431 | 8.29E-05 | 1.89E-05 | 1098.5902 | 2.60E-03 |
| Parp14 | -10.46 | 0.9668 | 1.6224 | 1.84E-01 | 2.14E-05 | 2913.8645 | 2.87E-03 |
| Kdm4a | -12.0858 | 1.2965 | 2813854.359 | 1.43E-01 | 2.43E-05 | 1076.4189 | 3.20E-03 |
| Nlrc5 | -11.0623 | 0.7172 | 1.1636 | 4.01E-05 | 2.76E-05 | 1691.9019 | 3.52E-03 |
| Duox2 | -13.7284 | 2.5604 | 0.0453 | 8.10E-05 | 3.01E-05 | 667.2265 | 3.79E-03 |
| Nectin2 | -11.3727 | 1.2736 | 0.3227 | 2.27E-01 | 3.17E-05 | 1918.9692 | 3.93E-03 |
| Lcn2 | -12.035 | 1.3827 | 0.0315 | 4.63E-05 | 3.71E-05 | 995.7003 | 4.41E-03 |
| Fmo1 | -9.3223 | 0.7651 | 1.6547 | 1.25E-01 | 3.84E-05 | 5285.7991 | 4.42E-03 |
| Cp | -9.6521 | 1.494 | 0.4069 | 4.21E-01 | 4.04E-05 | 5712.9131 | 4.52E-03 |
| Etfbkmt | -12.2125 | 1.2458 | 0.1318 | 9.76E-05 | 4.21E-05 | 928.4544 | 4.60E-03 |
| Meox1 | -13.4521 | 2.2002 | 0.0765 | 6.94E-02 | 4.24E-05 | 676.7075 | 4.61E-03 |
| Rtp4 | -11.4818 | 0.8728 | 0.3514 | 8.06E-05 | 4.45E-05 | 1329.3381 | 4.72E-03 |
| Gas7 | -13.1641 | 2.1675 | 0.0718 | 1.89E-01 | 4.58E-05 | 816.2447 | 4.79E-03 |
| 9930111J21Rik2 | -11.6267 | 0.8891 | 2.767 | 7.16E-05 | 5.02E-05 | 1232.5102 | 5.08E-03 |

**Endothelial Cell: gene expression, negative correlation (β_1_) with age**

| gene | Beta0 | Beta1 | Disp | Sigma | P.Beta | AIC | fdrP |
| --- | --- | --- | --- | --- | --- | --- | --- |
| Meg3 | -8.9562 | -1.5149 | 0.1679 | 7.61E-05 | 4.70E-32 | 2790.465 | 8.21E-28 |
| Dse | -10.1896 | -1.7585 | 0.2135 | 4.44E-05 | 6.22E-18 | 1115.608 | 3.28E-14 |
| Gap43 | -10.485 | -2.5123 | 1.6542 | 3.49E-05 | 7.50E-18 | 754.1488 | 3.28E-14 |
| Mdga1 | -10.6608 | -1.9318 | 0.3932 | 3.77E-05 | 9.47E-14 | 774.9051 | 1.84E-10 |
| Spag1 | -9.7824 | -1.6218 | 0.0334 | 5.91E-05 | 7.12E-12 | 1220.209 | 1.00E-08 |
| Gm16104 | -10.5292 | -1.3382 | 0.6369 | 6.72E-05 | 1.54E-11 | 1016.566 | 1.93E-08 |
| Srrm3 | -11.0673 | -2.1301 | 0.1121 | 5.07E-05 | 2.64E-09 | 532.0859 | 2.31E-06 |
| Gm43684 | -10.1896 | -4.0471 | 0.005 | 7.60E-05 | 7.64E-09 | 372.3594 | 5.34E-06 |
| Gm31615 | -11.2316 | -2.7864 | 0.2217 | 4.55E-05 | 8.79E-09 | 428.2712 | 5.92E-06 |
| Igf2bp3 | -10.9452 | -1.3142 | 0.3791 | 8.03E-05 | 6.80E-08 | 749.2149 | 3.13E-05 |
| Rian | -10.4354 | -1.0951 | 0.1169 | 4.98E-05 | 9.27E-08 | 1134.834 | 4.05E-05 |
| Klb | -11.2667 | -3.2667 | 0.1104 | 4.30E-05 | 9.83E-08 | 371.3677 | 4.19E-05 |
| Ticam1 | -10.4329 | -1.2247 | 0.0648 | 6.52E-05 | 1.04E-07 | 1025.685 | 4.32E-05 |
| CT030142.7 | -11.4134 | -2.136 | 0.1493 | 4.73E-05 | 2.14E-07 | 414.804 | 8.13E-05 |
| Gm15457 | -10.4653 | -3.593 | 0.0045 | 6.46E-05 | 2.64E-07 | 329.2919 | 9.42E-05 |
| Stra6 | -10.8711 | -1.1789 | 0.2289 | 4.04E-05 | 3.64E-07 | 803.5254 | 1.18E-04 |
| Nudt17 | -11.1315 | -1.2457 | 5.1553 | 1.12E-04 | 7.23E-07 | 649.5678 | 2.14E-04 |
| Mbd4 | -9.789 | -0.9233 | 0.0582 | 4.71E-04 | 8.73E-07 | 1680.194 | 2.50E-04 |
| Vamp9 | -11.5449 | -1.7863 | 0.4355 | 3.97E-06 | 1.96E-06 | 404.2321 | 4.70E-04 |
| Eno2 | -11.445 | -3.5007 | 0.0893 | 4.67E-05 | 2.49E-06 | 320.0683 | 5.73E-04 |
| Ank3 | -9.5097 | -1.5983 | 0.0184 | 2.09E-01 | 3.90E-06 | 1203.792 | 8.22E-04 |
| Kif11 | -11.0057 | -1.6489 | 0.0228 | 6.95E-05 | 4.26E-06 | 545.6294 | 8.76E-04 |
| Ret | -11.0593 | -1.1056 | 0.2873 | 3.61E-05 | 6.05E-06 | 719.1942 | 1.10E-03 |
| Fam189a1 | -11.5805 | -1.9438 | 0.0632 | 4.33E-05 | 7.78E-06 | 372.0498 | 1.33E-03 |
| Gm13708 | -11.7549 | -1.9349 | 2545151.63 | 2.87E-08 | 8.24E-06 | NA | 1.37E-03 |
| Frmd8os | -11.5183 | -1.6904 | 0.0866 | 4.23E-05 | 9.21E-06 | 411.6224 | 1.48E-03 |
| Acpp | -10.9401 | -0.9424 | 0.5096 | 3.05E-05 | 1.25E-05 | 808.7255 | 1.91E-03 |
| Mirg | -11.6553 | -2.2018 | 0.03 | 5.07E-05 | 1.60E-05 | 307.2475 | 2.36E-03 |
| Gm17655 | -10.3516 | -1.3224 | 0.0169 | 1.11E-04 | 2.65E-05 | 826.1289 | 3.44E-03 |
| Zfp935 | -10.7324 | -3.1933 | 0.0991 | 6.32E-01 | 2.72E-05 | 804.5036 | 3.50E-03 |
| Gm13427 | -11.765 | -1.7784 | 0.1355 | 3.88E-05 | 3.23E-05 | 331.156 | 3.98E-03 |
| S100a4 | -9.5423 | -1.8841 | 0.399 | 5.12E-01 | 3.25E-05 | 1796.671 | 3.98E-03 |
| Car7 | -10.88 | -1.6114 | 0.0141 | 6.18E-05 | 3.28E-05 | 530.819 | 3.98E-03 |
| Gm16845 | -10.8646 | -0.8585 | 0.3723 | 8.18E-05 | 3.47E-05 | 904.1943 | 4.18E-03 |
| Ngef | -11.8015 | -1.637 | 3338814.61 | 5.36E-05 | 3.97E-05 | 360.6693 | 4.51E-03 |
| Ip6k3 | -11.1053 | -0.9672 | 0.5368 | 3.70E-05 | 4.06E-05 | 740.3172 | 4.52E-03 |
| Borcs5 | -10.8283 | -0.7906 | 0.8318 | 3.27E-05 | 4.76E-05 | 944.5517 | 4.90E-03 |
| Gpr19 | -10.5417 | -1.0473 | 0.0373 | 1.01E-04 | 4.92E-05 | 927.5052 | 5.00E-03 |
| Gm18227 | -11.2408 | -4.3464 | 0.0118 | 5.61E-05 | 5.07E-05 | 278.1743 | 5.09E-03 |
| Gm45030 | -11.8043 | -2.6841 | 0.0263 | 5.07E-05 | 5.37E-05 | 255.3514 | 5.37E-03 |
| F630040K05Rik | -11.826 | -1.7347 | 0.1538 | 6.03E-05 | 5.96E-05 | 328.6836 | 5.84E-03 |
| Carf | -10.2848 | -1.0342 | 0.0285 | 6.07E-05 | 6.70E-05 | 1033.324 | 6.34E-03 |
| Ninj2 | -11.3841 | -1.3715 | 0.0495 | 9.25E-05 | 7.83E-05 | 494.5388 | 7.04E-03 |
| 4933439C10Rik | -10.9096 | -0.955 | 0.0917 | 4.66E-05 | 7.84E-05 | 817.6342 | 7.04E-03 |
| Efcab12 | -11.6709 | -1.5618 | 0.0649 | 3.79E-05 | 7.84E-05 | 367.9292 | 7.04E-03 |
| Igsf23 | -11.7105 | -1.3603 | 2548673.02 | 9.99E-23 | 8.85E-05 | 404.8744 | 7.70E-03 |
| Gm16157 | -11.9345 | -2.9914 | 0.0546 | 5.24E-05 | 9.25E-05 | 232.8885 | 7.93E-03 |
| Fkbp6 | -12.0942 | -2.4349 | 0.2227 | 1.78E-12 | 1.30E-04 | 221.0568 | 1.00E-02 |
| Gm35215 | -11.9926 | -2.9312 | 0.0486 | 5.10E-05 | 1.38E-04 | 220.3579 | 1.06E-02 |
| 4930506C21Rik | -12.0782 | -2.1644 | 0.1406 | 3.59E-05 | 1.45E-04 | 237.2944 | 1.10E-02 |

**Supplemental Tables S7:** NICHES table. Common gene signaling pairs followed by significant gene pairs associated with age-related adventitial remodeling in specific cell-type sender-receiving analyses.

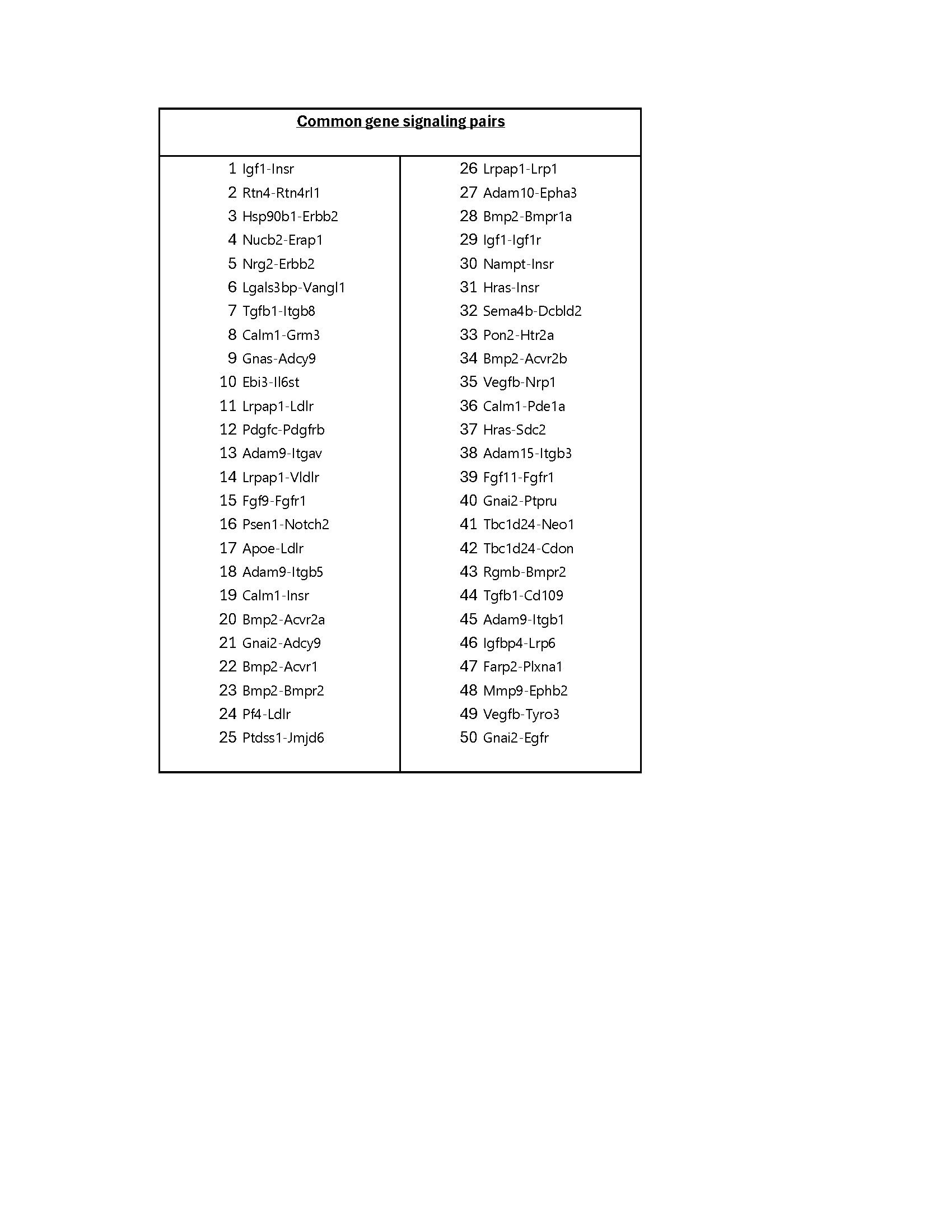

| **Signaling cell types** | **Signaling gene pair** | **p_val** | **avg_log2FC** | **pct.1** | **pct.2** | **p_val_adj** |
| --- | --- | --- | --- | --- | --- | --- |
| MacPeriVasc-Fibroblast | Calm1-Grm3 | 2.18E-59 | 4.50668425 | 0.716 | 0.048 | 2.47E-56 |
| MacPeriVasc-Fibroblast | Lgals3bp-Vangl1 | 4.42E-58 | 3.69389279 | 0.749 | 0.143 | 5.02E-55 |
| MacPeriVasc-Fibroblast | Il18-Il1rl2 | 4.81E-52 | 4.19162667 | 0.652 | 0.028 | 5.46E-49 |
| MacPeriVasc-Fibroblast | Nucb2-Erap1 | 8.47E-50 | 2.91340507 | 0.72 | 0.159 | 9.62E-47 |
| MacPeriVasc-Fibroblast | Igf1-Insr | 5.21E-49 | 3.6012284 | 0.655 | 0.048 | 5.91E-46 |
| MacPeriVasc-Fibroblast | Rtn4-Rtn4rl1 | 4.18E-48 | 3.54725637 | 0.643 | 0.071 | 4.75E-45 |
| MacPeriVasc-Fibroblast | Hsp90b1-Erbb2 | 2.17E-43 | 7.16589959 | 0.54 | 0.008 | 2.46E-40 |
| MacPeriVasc-Fibroblast | Vegfb-Nrp1 | 2.28E-42 | 3.10889271 | 0.637 | 0.103 | 2.59E-39 |
| MacPeriVasc-Fibroblast | Tgfb1-Itgb8 | 6.32E-42 | 3.69395814 | 0.585 | 0.04 | 7.18E-39 |
| MacPeriVasc-Fibroblast | Lrpap1-Vldlr | 1.68E-41 | 1.68279563 | 0.813 | 0.341 | 1.91E-38 |
| MacPeriVasc-Fibroblast | Ptdss1-Jmjd6 | 1.77E-41 | 3.43108261 | 0.614 | 0.091 | 2.01E-38 |
| MacPeriVasc-Fibroblast | Gnas-Adcy9 | 1.79E-41 | 2.68619074 | 0.673 | 0.103 | 2.03E-38 |
| MacPeriVasc-Fibroblast | Nrg2-Erbb2 | 4.19E-41 | 6.92423735 | 0.519 | 0.008 | 4.75E-38 |
| MacPeriVasc-Fibroblast | Pdgfc-Pdgfrb | 6.80E-41 | 2.05059091 | 0.738 | 0.278 | 7.72E-38 |
| MacPeriVasc-Fibroblast | Lrpap1-Lrp1 | 6.93E-41 | 1.37169841 | 0.871 | 0.417 | 7.87E-38 |
| MacPeriVasc-Fibroblast | Bmp2-Acvr2b | 3.52E-40 | 5.04032424 | 0.535 | 0.032 | 4.00E-37 |
| MacPeriVasc-Fibroblast | Igf1-Igf1r | 1.54E-39 | 1.59673705 | 0.801 | 0.345 | 1.74E-36 |
| MacPeriVasc-Fibroblast | Fgf9-Fgfr1 | 3.38E-38 | 2.62422958 | 0.639 | 0.147 | 3.83E-35 |
| MacPeriVasc-Fibroblast | Ebi3-Il6st | 2.43E-37 | 1.46867229 | 0.853 | 0.437 | 2.76E-34 |
| MacPeriVasc-Fibroblast | Adam9-Itgb5 | 4.90E-37 | 1.24213965 | 0.851 | 0.512 | 5.56E-34 |
| MacPeriVasc-SMC | Pon2-Htr2a | 1.11E-55 | 12.3695293 | 0.835 | 0 | 1.26E-52 |
| MacPeriVasc-SMC | Psen1-Notch3 | 9.87E-51 | 1.87000412 | 0.946 | 0.64 | 1.12E-47 |
| MacPeriVasc-SMC | Rtn4-Rtn4rl1 | 1.82E-50 | 3.98633107 | 0.838 | 0.093 | 2.07E-47 |
| MacPeriVasc-SMC | Igf1-Insr | 5.96E-48 | 2.01221948 | 0.937 | 0.379 | 6.77E-45 |
| MacPeriVasc-SMC | Hsp90b1-Erbb2 | 1.17E-47 | 3.72958346 | 0.793 | 0.019 | 1.33E-44 |
| MacPeriVasc-SMC | Nrg2-Erbb2 | 3.86E-47 | 6.36496756 | 0.763 | 0.006 | 4.38E-44 |
| MacPeriVasc-SMC | Gnai2-Ednra | 6.23E-46 | 2.03910658 | 0.931 | 0.236 | 7.07E-43 |
| MacPeriVasc-SMC | Gas6-Tyro3 | 1.24E-45 | 3.91559481 | 0.802 | 0.068 | 1.41E-42 |
| MacPeriVasc-SMC | Vegfb-Tyro3 | 1.34E-45 | 4.87777246 | 0.769 | 0.043 | 1.52E-42 |
| MacPeriVasc-SMC | Adam9-Itgav | 2.53E-45 | 2.41117178 | 0.877 | 0.273 | 2.87E-42 |
| MacPeriVasc-SMC | Psen1-Notch2 | 1.11E-44 | 2.42924246 | 0.862 | 0.304 | 1.25E-41 |
| MacPeriVasc-SMC | Pros1-Tyro3 | 1.64E-44 | 3.21236493 | 0.829 | 0.143 | 1.86E-41 |
| MacPeriVasc-SMC | Tgfb1-Itgb8 | 1.04E-43 | 2.38767586 | 0.865 | 0.13 | 1.19E-40 |
| MacPeriVasc-SMC | Nucb2-Erap1 | 1.49E-43 | 2.89834954 | 0.826 | 0.099 | 1.69E-40 |
| MacPeriVasc-SMC | Adam9-Itga3 | 2.63E-43 | 1.74367441 | 0.895 | 0.565 | 2.99E-40 |
| MacPeriVasc-SMC | Rtn4-Cntnap1 | 6.80E-43 | 4.39299447 | 0.757 | 0.056 | 7.72E-40 |
| MacPeriVasc-SMC | Lrpap1-Ldlr | 7.51E-42 | 4.72438182 | 0.73 | 0.037 | 8.52E-39 |
| MacPeriVasc-SMC | Lgals3bp-Vangl1 | 1.12E-41 | 11.2006616 | 0.691 | 0 | 1.27E-38 |
| MacPeriVasc-SMC | Ebi3-Il6st | 2.79E-40 | 2.67500225 | 0.826 | 0.087 | 3.16E-37 |
| MacPeriVasc-SMC | Gnas-Adcy1 | 5.63E-38 | 2.95380927 | 0.79 | 0.106 | 6.39E-35 |
| Myeloid-Fibroblast | Calm1-Grm3 | 1.12E-27 | 4.11495659 | 0.687 | 0.062 | 1.27E-24 |
| Myeloid-Fibroblast | Il18-Il1rl2 | 3.18E-22 | 2.62914435 | 0.662 | 0.069 | 3.61E-19 |
| Myeloid-Fibroblast | Sema4d-Erbb2 | 3.29E-21 | 12.2715709 | 0.503 | 0 | 3.74E-18 |
| Myeloid-Fibroblast | Rtn4-Rtn4rl1 | 5.83E-21 | 3.06125389 | 0.59 | 0.077 | 6.62E-18 |
| Myeloid-Fibroblast | Lrpap1-Lrp1 | 1.50E-20 | 2.21041281 | 0.662 | 0.115 | 1.70E-17 |
| Myeloid-Fibroblast | Lrpap1-Vldlr | 1.83E-20 | 2.68818771 | 0.631 | 0.1 | 2.07E-17 |
| Myeloid-Fibroblast | Tgfb1-Itgb6 | 9.12E-20 | -2.4542809 | 0.154 | 0.631 | 1.03E-16 |
| Myeloid-Fibroblast | Tgfb1-Itgb8 | 2.33E-19 | 3.06042542 | 0.564 | 0.046 | 2.65E-16 |
| Myeloid-Fibroblast | Hsp90b1-Erbb2 | 3.97E-19 | 11.5887875 | 0.462 | 0 | 4.50E-16 |
| Myeloid-Fibroblast | Lgals3bp-Vangl1 | 1.20E-18 | 4.95842559 | 0.492 | 0.031 | 1.36E-15 |
| Myeloid-Fibroblast | Hras-Insr | 2.34E-18 | 3.34291156 | 0.497 | 0.023 | 2.66E-15 |
| Myeloid-Fibroblast | Fn1-Itgb6 | 2.73E-18 | -3.4745921 | 0.077 | 0.485 | 3.10E-15 |
| Myeloid-Fibroblast | Ptdss1-Jmjd6 | 4.55E-17 | 3.75672188 | 0.508 | 0.077 | 5.16E-14 |
| Myeloid-Fibroblast | Gnas-Adcy9 | 6.51E-17 | 1.86430333 | 0.692 | 0.185 | 7.39E-14 |
| Myeloid-Fibroblast | Calm1-Insr | 2.74E-16 | 1.24128035 | 0.667 | 0.115 | 3.11E-13 |
| Myeloid-Fibroblast | Nampt-Insr | 7.34E-16 | 1.21592471 | 0.626 | 0.108 | 8.33E-13 |
| Myeloid-Fibroblast | Calm2-Insr | 1.36E-15 | 0.83951521 | 0.605 | 0.092 | 1.54E-12 |
| Myeloid-Fibroblast | Hras-Sdc2 | 1.28E-14 | 1.91065434 | 0.59 | 0.154 | 1.46E-11 |
| Myeloid-Fibroblast | Mfng-Notch2 | 1.35E-14 | 2.02276097 | 0.508 | 0.077 | 1.53E-11 |
| Myeloid-Fibroblast | S100a8-Tlr4 | 3.93E-14 | 4.39464445 | 0.405 | 0.031 | 4.46E-11 |
| Myeloid-SMC | Gnai2-Ednra | 2.69E-22 | 2.4809227 | 0.902 | 0.239 | 3.06E-19 |
| Myeloid-SMC | Pon2-Htr2a | 4.39E-22 | 11.1998938 | 0.75 | 0 | 4.99E-19 |
| Myeloid-SMC | Rtn4-Rtn4rl1 | 8.69E-21 | 4.07306333 | 0.774 | 0.099 | 9.86E-18 |
| Myeloid-SMC | Pros1-Tyro3 | 1.25E-20 | 3.68875145 | 0.78 | 0.056 | 1.42E-17 |
| Myeloid-SMC | Psen1-Notch3 | 1.17E-19 | 1.30242219 | 0.939 | 0.746 | 1.33E-16 |
| Myeloid-SMC | Sema4d-Erbb2 | 1.39E-19 | 3.6331958 | 0.744 | 0.028 | 1.58E-16 |
| Myeloid-SMC | Lrpap1-Sort1 | 2.43E-19 | 3.77776103 | 0.756 | 0.085 | 2.76E-16 |
| Myeloid-SMC | Tgfb1-Itgb8 | 6.48E-19 | 2.61098911 | 0.823 | 0.169 | 7.35E-16 |
| Myeloid-SMC | Rtn4-Cntnap1 | 7.72E-19 | 4.72944892 | 0.713 | 0.042 | 8.76E-16 |
| Myeloid-SMC | Hsp90b1-Erbb2 | 2.58E-18 | 4.82731371 | 0.689 | 0.014 | 2.92E-15 |
| Myeloid-SMC | Adam9-Itga3 | 2.72E-17 | 1.63797058 | 0.866 | 0.38 | 3.08E-14 |
| Myeloid-SMC | Adam9-Itgav | 5.55E-17 | 1.96653111 | 0.841 | 0.169 | 6.30E-14 |
| Myeloid-SMC | Lrpap1-Vldlr | 9.24E-17 | 2.89551516 | 0.756 | 0.113 | 1.05E-13 |
| Myeloid-SMC | Lrpap1-Sorl1 | 2.62E-16 | 2.74905962 | 0.756 | 0.127 | 2.97E-13 |
| Myeloid-SMC | Psen1-Notch2 | 2.83E-16 | 1.70210574 | 0.848 | 0.394 | 3.21E-13 |
| Myeloid-SMC | Lrpap1-Lrp1 | 4.31E-16 | 2.7650378 | 0.744 | 0.099 | 4.89E-13 |
| Myeloid-SMC | Gnas-Adcy1 | 5.47E-16 | 2.59588551 | 0.774 | 0.169 | 6.21E-13 |
| Myeloid-SMC | Ebi3-Il6st | 6.65E-16 | 1.97339823 | 0.732 | 0.07 | 7.55E-13 |
| Myeloid-SMC | Calm1-Grm3 | 7.04E-16 | 2.22203503 | 0.823 | 0.169 | 7.99E-13 |
| Myeloid-SMC | Lrpap1-Ldlr | 8.78E-16 | 5.21552699 | 0.616 | 0.014 | 9.97E-13 |
